# Ca^2+^-independent ZmCPK2 is inhibited by Ca^2+^-dependent ZmCPK17 during drought response in maize

**DOI:** 10.1101/2024.03.15.585280

**Authors:** Xiaoying Hu, Jinkui Cheng, Minmin Lu, Tingitng Fang, Yujuan Zhu, Zhen Li, Xiqing Wang, Yu Wang, Yan Guo, Shuhua Yang, Zhizhong Gong

## Abstract

Calcium oscillations are induced by different stresses. Calcium-dependent protein kinases (CDPKs/CPKs) are one major group of the plant calcium decoders that are involved in various processes including drought response. Some CPKs are calcium-independent. Here, we identified ZmCPK2 as a negative regulator of drought resistance by screening an overexpression transgenic maize pool. We found that ZmCPK2 does not bind calcium, and its activity is modulated likely in an oscillation manner during short time ABA treatment. Interestingly, ZmCPK2 interacts with and is inhibited by calcium-dependent ZmCPK17, a positive regulator of drought resistance, which is activated by ABA. ZmCPK17 activity exhibits opposite patterns relative to ZmCPK2 under ABA treatment, and could prevent the nuclear localization of ZmCPK2 when co-expressed. ZmCPK2 interacts with and phosphorylates ZmYAB15, a negative transcriptional factor for drought resistance. Our results suggest that drought stress-induced Ca^2+^ can be decoded by corresponding CPK17 likely in an oscillation manner, thereby promoting plant adaptation to water deficit.

## INTRODUCTION

Calcium is an important “language” that triggers signal transduction in plants, and calcium oscillations occur rapidly under biotic and abiotic stresses (Dong et al., 2022; Sanders et al., 2002). Different stimuli do not completely trigger uniform oscillatory parameters such as frequency and amplitude within the cell or microdomain (Smedler and Uhlen, 2014). Unique intracellular calcium signals induced by different environmental cues are recognized by various calcium sensors, thereby transmitting calcium signals and triggering a series of specific reactions in plants. Plants have four major calcium sensors, including calmodulin (CaM) and calmodulin dependent protein kinase (CaMK) (Means and Dedman, 1980; Watillon et al., 1993), calcineurin B-likes and its interacting protein kinases (CBLs-CIPKs) (Kudla et al., 1999; Shi et al., 1999), calcium/calmodulin dependent protein kinases (CCaMKs) (Patil et al., 1995), and calcium dependent protein kinases (CDPKs/CPKs) (Harmon et al., 1987). Among these sensors that rely on phosphorylation cascade to transmit signals, CPKs are special as they integrate the binding of calcium ion and the transmission of signals in the same molecule. Thus, CPKs are considered to be the combination of CaM and CaMK in evolution (Harper et al., 1991; Suen and Choi, 1991).

CPKs are a class of protein kinases widely found in plants and some protozoa with conserved structure, including a variable N-terminal domain, followed by a Ser/Thr kinase domain, a junction domain and a C-terminal calcium regulatory domain (Delormel and Boudsocq, 2019; Harmon et al., 2000). Different CPKs contain different numbers of EF-hands, which results in various binding abilities to calcium (Boudsocq et al., 2012; Cheng et al., 2002). By analyzing the structure of *Toxoplasma gondii* CPK, the key functional domains and amino acids of plant CPKs, a general model of calcium-dependent CPK activation mechanism was obtained (Liese and Romeis, 2013; Wernimont et al., 2010). In the resting state, the concentration of intracellular calcium ions is relatively low, and the activity of CPK kinase is inhibited by the junction domain, and it is inactivated; under biotic or abiotic stress conditions, intracellular calcium concentration increases, and C-terminal EFs bind to calcium, resulting in conformational changes, prompting the dissociation of the linking domain from the kinase domain, releasing kinase activity, and then transmitting signals through phosphorylation of downstream targets (Liese and Romeis, 2013; Wernimont et al., 2010). This mechanism is supported by different lines of evidence (Delormel and Boudsocq, 2019).

CPKs are involved in various signal transduction pathways (Delormel and Boudsocq, 2019; Edel and Kudla, 2015). Previous studies have found that both calcium-dependent and calcium-independent CPKs are involved in plant response to abiotic stress, but the response mechanism is still unclear (Delormel and Boudsocq, 2019). For example, in *Arabidopsis thaliana*, the ABA-activated AtCPK4 and AtCPK11 can positively regulate ABA and drought stress response by phosphorylating ABSCISIC ACID-RESPONSIVE ELEMENT BINDING FACTORs (ABFs), the key transcription factors in ABA signaling pathway (Zhu et al., 2007), or mediate ethylene biosynthesis through phosphorylating C-terminus of 1-aminocyclopropane-1-carboxylate synthase 6 and stabilizing it under ABA treatment (Luo et al., 2014). AtCPK32 can interact with and phosphorylate ABF4 to regulate ABA signal pathway (Choi et al., 2005). AtCPK3, AtCPK5, AtCPK6, AtCPK21, AtCPK23 in *Arabidopsis* and ZmCPK35, ZmCPK37 in maize can all phosphorylate and activate Slow Anion Channel 1 (SLAC1) and promote ABA-induced stomatal closure (Brandt et al., 2012; Brandt et al., 2015; Demir et al., 2013; Geiger et al., 2010; Li et al., 2022a; Mori et al., 2006; Scherzer et al., 2012). ZmCDPK7 enhances thermotolerance in maize (Zhao et al., 2021).

Some studies have shown that in addition to the classical activation mechanism of calcium, CPKs are also inhibited by phosphatases and may be regulated by upstream kinases. For example, CPK3/6/21/23 can be inhibited by the clade-A PP2C type ABA-insensitive 1 (ABI1) for their phosphorylation on SLAC1 (Geiger et al., 2010; Scherzer et al., 2012). Some kinases phosphorylated by other kinases, such as the MAPK cascade and RAFs-SnRK2s module, are common and play important signal transduction functions (Chen et al., 2021; Zhang and Zhang, 2022). In *Arabidopsis*, AtCPK11 interacts with and phosphorylates AtCPK24, and together they inhibit the Shaker potassium channel SPIK (AKT6) as no single kinase can inhibit SPIK (Zhao et al., 2013). Apart from this, there are few researches about upstream kinases on the regulation of calcium-independent CPKs (Delormel and Boudsocq, 2019).

Maize is one of the world’s most important crops, and its production is greatly threatened by drought, especially with the serious global warming (Yang and Qin, 2023). Thus, searching for maize drought resistance candidate genes and exploring maize drought resistance response mechanism are of great significance for breeding drought resilience maize species. In order to identify the candidate genes involved in drought response, we screened a number of maize transgenic lines. In this study, we found that ZmCPK2 is a negative drought regulator that can phosphorylate and activate the transcription regulator ZmYAB15. ZmYAB15 negatively regulates drought resistance. Another calcium-dependent member, ZmCPK17, is a positive regulator that phosphorylates and inhibits ZmCPK2 and ZmYAB15 in plant responses to drought stress. Interestingly, ZmCPK2 and ZmCPK17 activity showed opposite patterns likely in an oscillation manner during ABA treatment. Thus, these proteins form a complex module that regulates the balance between plant growth and drought stress.

## RESULTS

### ZmCPK2 is a negative regulator of drought resistance in maize

In order to search for genes associated with drought stress, we screened a pool of transgenic maize seedlings overexpressing more than 700 genes in inbred line LH244 background (generated by Center for Crop Functional Genomics and Molecular Breeding, CAU) by a soil drought treatment assay. We found that two independent *ZmCPK2* overexpression lines (*ZmCPK2-OE1* and *-OE2*) showed stronger drought sensitivity and had lower leaf relative water content compared with LH244 (Figure 1A, 1B). Please notice that CPK naming is a bit confusing (Kong et al., 2013; Weckwerth et al., 2015). Here we use the CPK naming in an earlier published paper (Kong et al., 2013). qRT-PCR assay showed the transcriptional levels of *ZmCPK2* were higher in *ZmCPK2-OE1* and *-OE2* than LH244 (Figure S1A). We further obtained four mutants of *ZmCPK2* using CRISPR/Cas9 technique: *zmcpk2-1* has a single base G deletion, thus resulting in premature termination of the protein; *zmcpk2-2* has three base deletion, one amino acid missing in the putative protein; *zmcpk2-3* and *zmcpk2-4* with truncated peptides at the C-terminus of ZmCPK2, deleting the 4 EF-hands after junction domain (Figure S1B). In the soil drought treatment assay, *zmcpk2-1* showed a drought-tolerant phenotype with higher leaf relative water content compared with LH244 (Figure 1C, 1D). In order to investigate the effect of ZmCPK2 on maize adult plants under drought stress, we carried out field soil drought treatment in a rain shelter to control the water supply, and found that *ZmCPK2-OE2* showed drought sensitive phenotypes such as young leaves with earlier wilting and old leaves with earlier senescence compared to LH244, while *zmcpk2-1* showed the opposite phenotypes of *ZmCPK2-OE2* (Figure S1C). Meanwhile, there were no significant different phenotypes between *zmcpk2-2* and wild type in soil drought test (Figure S2A, S2B), suggesting that lacking 3-bp in *zmcpk2-2* does not affect its activity. However, *zmcpk2-3* and *2-4* exhibited a more drought sensitive phenotype similar to *ZmCPK2-OE* lines compared to the wild type, suggesting that these two mutants are gain-of-function ones (Figure S2C). Consistently, the water loss assay of detached leaves indicated that *ZmCPK2-OE1/2* and *zmcpk2-3/-4* lost water faster, and *zmcpk2-1* slower compared to LH244 (Figure 1E, S2D). The leaf temperature can better reflect the transpiration rate: the lower the temperature, the higher the transpiration rate (Sun et al., 2022). We utilized a far-red CCD camera to compare the leaf temperatures among the wild type, *ZmCPK2-OE1/2* and *zmcpk2-1/-3/-4* mutants. As shown in Figure 1F, 1G, S2E and S2F, the wild type leaves had higher temperatures than *ZmCPK2-OE1/2* or *zmcpk2-3/-4,* and lower temperatures than *zmcpk2-1*, indicating that *zmcpk2-1* transpired less water, and *ZmCPK2-OE1/2* or *zmcpk2-3/-4* transpired more water than the wild type. We also examined the stomatal movement of *ZmCPK2-OE* and *zmcpk2* under ABA treatment. The results showed that the stomatal apertures of both *ZmCPK2-OE1/2* and *zmcpk2-1/-3/-4* were not significantly different from that of LH244 without ABA treatment (Mock). However, *ZmCPK2-OE1/2* and *zmcpk2-3/-4* were less sensitive, while *zmcpk2-1* was more sensitive to ABA-induced stomatal closure compared with LH244 (Figure 1H). We did not observe any different growth and development phenotypes of *ZmCPK2-OE1/2*, *zmcpk2-1* and LH244 seedlings under well-watered condition (Figure S2G). We also obtained the data of plant height, hundred grain weight and plot yield of mature plants of *ZmCPK2-OE1/2* in the field. The statistical analysis showed that the height of *ZmCPK2-OE* was no different from LH244 in some regions, and a little shorter than LH244 in other regions (Figure S2H), while the hundred grain weight and plot yield were similar to LH244 (Figure S2I, S2J). Thus, ZmCPK2 acts as a negative regulator in maize drought resistance.

**Figure 1.**
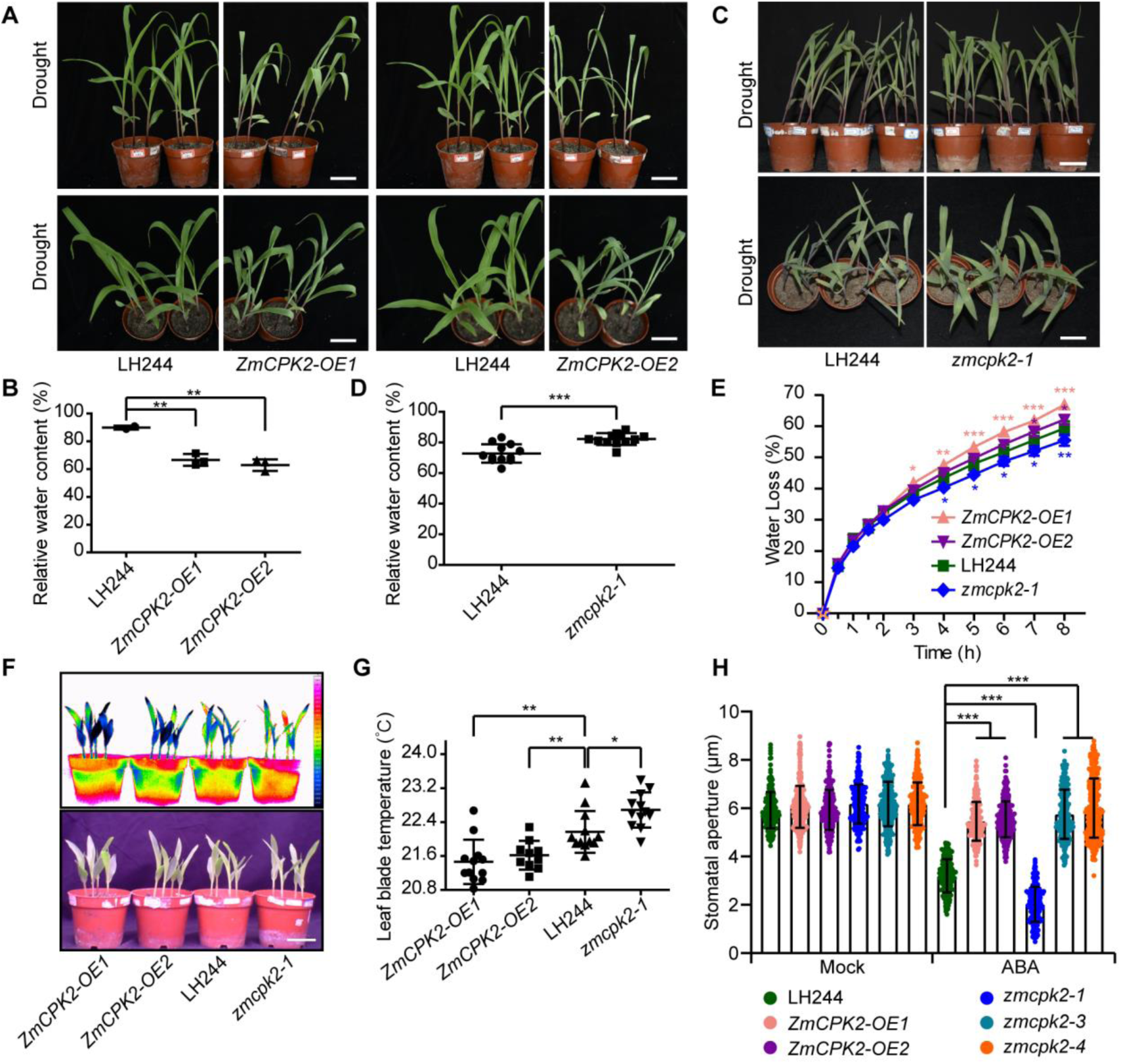
ZmCPK2 is a negative regulator for drought tolerance. **(A)** Transgenic *ZmCPK2-OE1/2* maize seedlings are more sensitive to drought treatment compared with LH244. The pictures were taken for 14-d-old seedlings treated with drought stress. Bar = 5 cm. **(B)** Relative water content of leaves was calculated in **(A)**. The fresh weight, saturated fresh weight and dry weight of all cut leaves were measured for calculating the relative water content. Each point represents a group of leaves from one seedling. Error bars represent ± SD from one experiment, and three experiments were performed with similar results. * Represents significant difference with LH244, ** p<0.01, Student’s *t*-test. **(C)** The CRISPR/Cas9 mutant *zmcpk2-1* is more tolerant to drought treatment compared with LH244. The pictures were taken for 12-d-old seedlings treated with drought stress. Bar = 5 cm. **(D)** Relative water content of leaves was calculated in **(C)**. Each point represents a group of leaves from one seedling. Error bars represent ± SD from one experiment, and three experiments were performed with similar results. * represents significant difference with LH244, *** p<0.001, Student’s *t*-test. **(E)** Comparison of water loss of *ZmCPK2-OE1/2*, *zmcpk2-1* and LH244. Error bars represent ± SD of three technical replicates, each with three leaves form the second expanded leaf of three seedlings. * Represents significant difference with LH244, * p<0.05, ** p<0.01, *** p<0.001, Student’s *t*-test. **(F)** Comparison of the leaf temperatures of *ZmCPK2-OE1/2*, the wild type and *zmcpk2-1* as determined by an infrared thermal imaging assay. 8-d-old seedlings grown in soil with well watering were used for temperature measurement with a far-red CCD camera. **(G)** Statistical analysis of leaf temperature in **(F)**. Data are means ± SD of three replicates (eight leaves from one pot were measured per replicate) in one experiment. Four experiments were done with similar results. * represents significant difference with LH244, *p < 0.05, **p < 0.01, Student’s *t*-test. **(H)** Measurement of stomatal aperture with (ABA) or without (Mock) 12 µM ABA treatment. Three hundred stomata of three leaves from three seedlings were measured in one experiment. Three experiments were done with similar results. Data are means ± SD. * Represents significant difference with LH244, *** p<0.001, Student’s *t*-test.

### ZmCPK2 is a calcium-independent CPK

Protein amino acid sequence alignment indicated that ZmCPK2 has the highest identity with ZmCPK7 in maize, and AtCPK5/6/26 in *Arabidopsis* (Figure S3A). Further sequence analysis showed that ZmCPK2 only has one intact EF hand (EF3) and three degenerated ones (Figure S3B). Previous studies reported that ZmCPK7 is a calcium-independent CPK, which has calcium-independent kinase activity and low calcium binding ability (Boudsocq et al., 2012; Weckwerth et al., 2015). We first tested the calcium binding ability of ZmCPK2. With the *in vitro* microscale thermophoresis (MST) assay, we found that ZmCPK2 had almost no calcium binding capacity with a very high Kd value (10914 ± 14580 nM), while the calcium-dependent CPK, AtCPK3, had a high calcium binding capacity with a relative lower Kd value (1533.9 ± 613.74 nM) (Figure 2A) (Boudsocq et al., 2012). The Kd value reflects the binding capacity, and the smaller the Kd value, the stronger the binding capacity. We used empty GST with no calcium-binding capacity as a negative control (Figure 2A). We then detected the kinase activity of ZmCPK2 by the *in vitro* kinase assay, and found that ZmCPK2 had autophosphorylation activity and could phosphorylate its target ZmYAB15 (see later for identification of ZmYAB15) (Figure 2B). To explore whether the calcium affects its activity, we added different concentrations of free Ca^2+^ to measure kinase activity. The results showed that the activity of ZmCPK2 was not affected by low calcium concentration, but became a little lower with addition of more calcium (Figure 2B, 2C), suggesting that ZmCPK2 was not activated by Ca^2+^. To further explore the Ca^2+^-independence of ZmCPK2, we compared the activity of truncated forms or EF hand mutated forms (Piazza et al., 2017) of ZmCPK2. It was noticed that the deletion of 4 EF-hands of ZmCPK2 (ZmCPK2-NKJ) or 4 EF-hands and junction domain of ZmCPK2 (ZmCPK2-NK) could release its autoinhibition effect, and its autophosphorylation activity was much higher than that of intact ZmCPK2, but the phosphorylation activity on ZmYAB15 was not different with intact ZmCPK2 (Figure 2D, S3B). Meanwhile, mutation in each EF hand did not affect the kinase activity of ZmCPK2, except for the nonfunctional EF1 which reduced its activity (Figure 2E, S3B). These results indicate that ZmCPK2 is a calcium-independent CPK.

**Figure 2.**
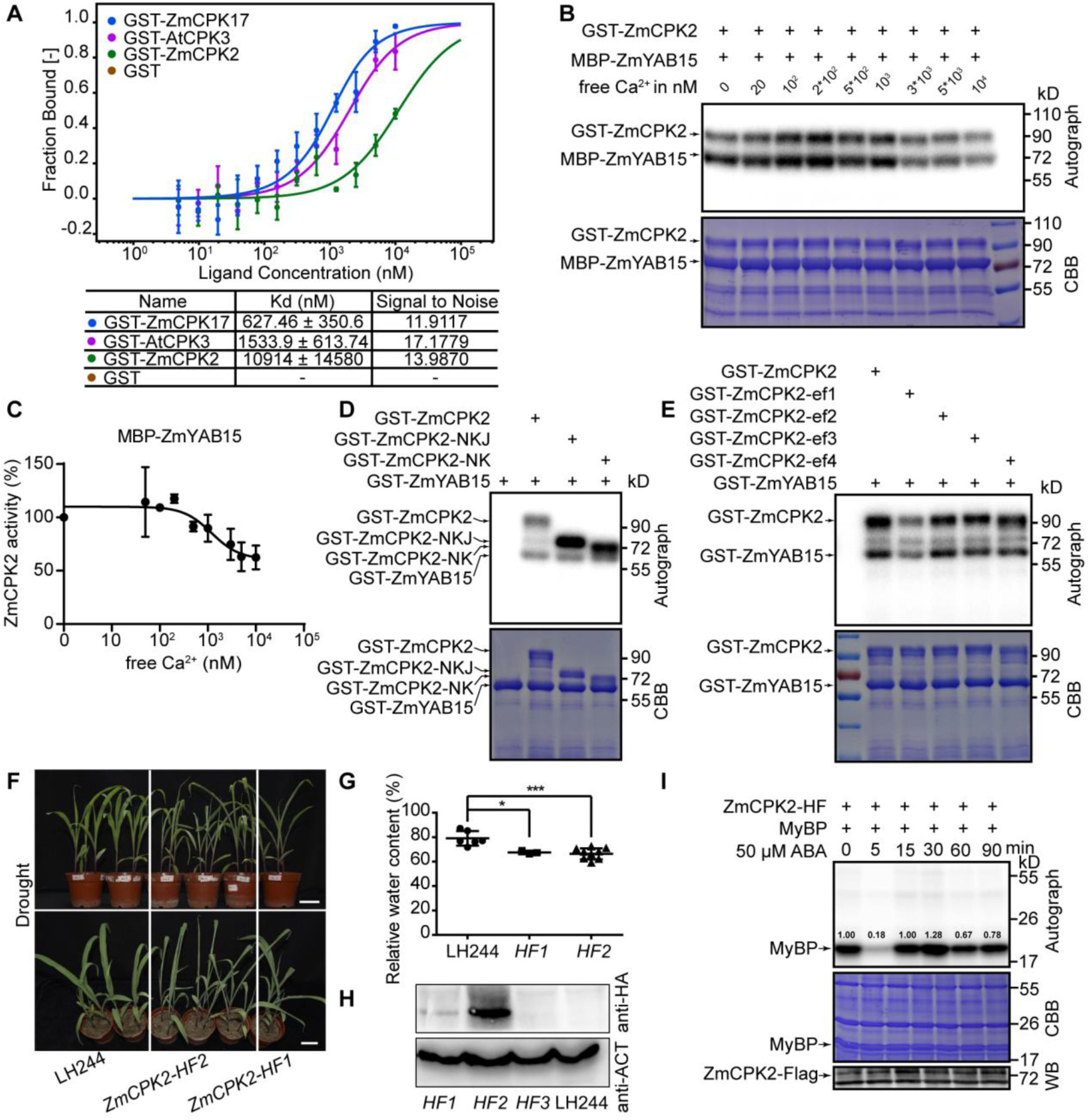
The kinase activity of ZmCPK2 is independent on Ca^2+^, and regulated by ABA. **(A)** ZmCPK2 does not bind calcium as shown by microscale thermophoresis (MST) assay. Recombinant GST-ZmCPK2 and GST-ZmCPK17 were used as receptor proteins and different concentrations of Ca^2+^ were used as ligand. GST-AtCPK3 was used as a positive control and GST was used as a negative control. Data were three independent experiments. **(B)** ZmCPK2 has calcium-independent kinase activity. Recombinant GST-ZmCPK2 were subjected to the phosphorylation assay with recombinant MBP-ZmYAB15 as the substrate in various Ca^2+^ concentrations. The autograph and the Coomassie Brilliant Blue-stained gel (CBB) were shown. **(C)** Quantitation of the relative signal of MBP-ZmYAB15 with Image J in **(B)** and analyzed by a Hill equation with a Hill factor of four in Graghpad Prism 8. ZmCPK2 exhibited high activity with no or low free Ca^2+^ concentration. Error bars represent ± SD from two experiments. **(D)** Comparing the kinase activity of different truncated forms of ZmCPK2 *in vitro*. Recombinant GST-ZmCPK2, GST-ZmCPK2-NKJ and GST-ZmCPK2-NK were subjected to the phosphorylation assay with GST-ZmYAB15 as the substrate. **(E)** Comparing the kinase activity of ZmCPK2 with different mutations in four EF hands. Recombinant GST-ZmCPK2 and GST-ZmCPK2-ef1/2/3/4 were subjected to the phosphorylation assay *in vitro* with GST-ZmYAB15 as the substrate. **(F)** Transgenic HA-Flag-tagged seedlings *ZmCPK2-HF1/2* are more sensitive to drought treatment compared with LH244. The pictures were taken from 14-d-old seedlings treated with drought stress. Bar = 5 cm. **(G)** Relative water content of leaves was calculated in **(F)**. Error bars represent ± SD from one experiment, and three experiments were performed with similar results. * represents significant difference with LH244, * p<0.05, *** p<0.001, Student’s *t*-test. **(H)** The protein levels of ZmCPK2 in *ZmCPK2-HF1/2/3*. Total proteins were extracted from 8-d-old seedlings and the protein levels of ZmCPK2-HA-Flag were detected with anti-HA antibodies. Actin was used as an internal control. **(I)** The kinase activity of ZmCPK2 is regulated by ABA. The total proteins were extracted from 8-d-old seedlings of *ZmCPK2-HF2* treated with 50 µM ABA for different times. ZmCPK2-HA-Flag (ZmCPK2-HF) was immunoprecipated (IPed) from total proteins with anti-Flag agarose. IPed proteins were subjected to *in vitro* phosphorylation assay using MyBP as a substrate. The autograph, CBB and the protein levels of ZmCPK2 in each sample (WB) were shown. The numbers represent the quantification of the phosphorylation signals of MyBP by the Image J software. Three experiments were done with similar expression patterns.

Considering that ZmCPK2 negatively regulates the drought/ABA response, we further investigated whether its activity is regulated by ABA. As we did not have ZmCPK2 antibodies, we generated two transgenic lines overexpressing *Ubi:ZmCPK2-HA-FLAG* (*ZmCPK2-HF1*, *-HF2*) for further study. The drought phenotype showed that they exhibited similar drought sensitivity as *ZmCPK2-OE1/2* with lower leaf relative water content (Figure 2F, 2G), and were overexpressed as detected by anti-HA antibodies (Figure 2H). We then treated the 8-d-old *ZmCPK2-HF2* seedlings with 50 μM ABA, and immunoprecipitated the total proteins with anti-Flag agarose, conducted *in vitro* phosphorylation assay using the general kinase substrate myelin basic protein (MyBP). Three independent biological experiments were performed with similar results. ABA treatment greatly inhibited the activity of ZmCPK2 at about 5 min, and then its activity reached to a normal level at 15 min and a higher level at 30 min, and then reduced to a lower level at later treatments (Figure 2I, S7A, S7B). These results suggest that ZmCPK2 activity is dynamically regulated during ABA treatment.

### ZmCPK2 interacts with ZmCPK17

Considering that ZmCPK2 is a calcium-independent CPK, its activity is dynamically regulated by ABA, we speculated that ZmCPK2 may be directly regulated by some other calcium-responsive upstream protein kinases under drought stress. To verify this possibility, we utilized ZmCPK2^D201A^ (D201 is a putative active center site of the kinase, and Asp201 mutated to Ala to form an inactive kinase as the active one has autoactivation in yeast cells) as a bait to screen its potential interaction proteins from other ZmCPKs by yeast two-hybrid assay. We found that ZmCPK2 indeed interacts with some ZmCPKs including ZmCPK6, ZmCPK9, ZmCPK14, ZmCPK17, but not with ZmCPK7, ZmCPK10, ZmCPK11, ZmCPK15, ZmCPK26 (Figure S3C). Meanwhile, *in vitro* phosphorylation assay showed that ZmCPK6/9/14/17 could all phosphorylate ZmCPK2 in a Ca^2+^-dependent manner (Figure S3D). Among these interacted CPKs, we chose ZmCPK17 as a representative for further study. Interaction assay with truncated ZmCPK2 proteins indicated that ZmCPK17 only interacted with ZmCPK2-NKJ but not with other truncated forms (Figure S4A, S4B). The interaction of *Arabidopsis* OST1 and ABI1 was used as a positive control (Deng et al., 2022). We also performed pull-down assay and confirmed the interaction between ZmCPK2 and ZmCPK17 (please see later in Figure S10E). The results showed that ZmCPK2 interacted with ZmCPK17 in vitro and the interaction was strengthened by Ca^2+^. We transiently expressed ZmCPK17-GFP in transgenic *ZmCPK2-HF1* protoplasts and did co-immunoprecipitation assay (Chen et al., 2008). The immunoprecipitated proteins with anti-GFP antibodies could be detected by anti-Flag antibodies, indicating the interaction of ZmCPK2 and ZmCPK17 *in vivo* (Figure S4C). We further performed Bimolecular Fluorescence Complementation (BiFC) assay, and found that ZmCPK2 interacted with ZmCPK17 (Figure S4D). GUS was used as a negative control for BiFC assay (Figure S4D). These results demonstrated that ZmCPK2 physically interacts with ZmCPK17.

### Calcium-dependent ZmCPK17 is a positive regulator of drought response

To explore whether ZmCPK17 can directly respond to calcium, we performed MST assay and found that ZmCPK17 had strong calcium binding capacity, similar to AtCPK3, with a lower Kd (627.46 ± 350.6 nM) (please see former Figure 2A). We then measured the kinase activity of ZmCPK17 *in vitro* using proteins purified from *Escherichia coli*. The result showed that ZmCPK17 had strong autophosphorylation activity and could phosphorylate MBP-ZmCPK2^D201A^, which was increased with presence of calcium. The Ca^2+^-dependent phosphorylation activity of ZmCPK17 is best fitted by a Hill equation with a Hill constant of four (Figure 3A, 3B). We also compared the activity of truncated forms or EF hand mutation forms of ZmCPK17. *In vitro* phosphorylation assay showed that ZmCPK17-NKJ and ZmCPK17-NK had little kinase activity (Figure S4E, S4F), and the mutation of each EF hand (ef) reduced the activity of ZmCPK17, especially the EF hand 3 and 4 (Figure S4E, S4G). Next, we checked drought phenotype of ZmCPK17 overexpression transgenic lines, *ZmCPK17-OE1* and *-OE2,* as well as *zmcpk17-1, -2* mutants created by CRISPR/Cas9 (Zeng et al., 2021). *zmcpk17-1* changes C120 [counting from the putative translation start codon A (1)TG] to TA, thus producing a frame shift and an early putative translation stop codon after Pro185. *zmcpk17-2* has a deletion of 4 bases (AGAC) after A121 and produces an early putative translation stop codon after Thr172 (Figure S5A). Both *ZmCPK17-OE1* and *-OE2* had a higher expression level of *ZmCPK17* than the wild type as checked by qRT-PCR (Figure S5B). We did not observe the difference in plant growth and development among these OE lines, mutants and the wild type under normal growth conditions (Figure S5C, S5D, S5E, S5F). From the field test data, we found that the height and hundred grain weight of *ZmCPK17-OE* were almost similar to LH244, except that *ZmCPK17-OE2* was a little higher in one the region, and *ZmCPK17-OE1* was a little shorter in two regions than LH244 (Figure S5G, S5H). The soil drought treatment experiments showed that seedlings of *ZmCPK17-OE1* and *-OE2* displayed stronger drought tolerance and higher leaf water content than those of the wild type LH244. In contrast, *zmcpk17-1,* and *-2* mutants had higher drought sensitivity and lower leaf water content than the wild type LH244 (Figure 3C, 3D, 3E, 3F). The stomatal aperture assay showed that t *ZmCPK17-OE1/2* and *zmcpk17-1/-2* had no significant difference from that of LH244 without ABA treatment (Mock), while *ZmCPK17-OE1/2* exhibited more sensitive, and *zmcpk17-1/-2* were less sensitive to ABA-induced stomatal closure than the wild type (Figure 3G). Consistently, the leaves of *ZmCPK17-OE1* and *-OE2* seedlings had higher temperatures, while *zmcpk17-1, -2* mutants had lower temperatures than those of the wild type (Figure 3H, 3I, 3J, 3K), indicating that *ZmCPK17-OE1* and *-OE2* transpire less water, while *zmcpk17-1, -2* transpire more water than the wild type. Thus, calcium-dependent ZmCPK17 is a positive regulator of drought resistance.

**Figure 3.**
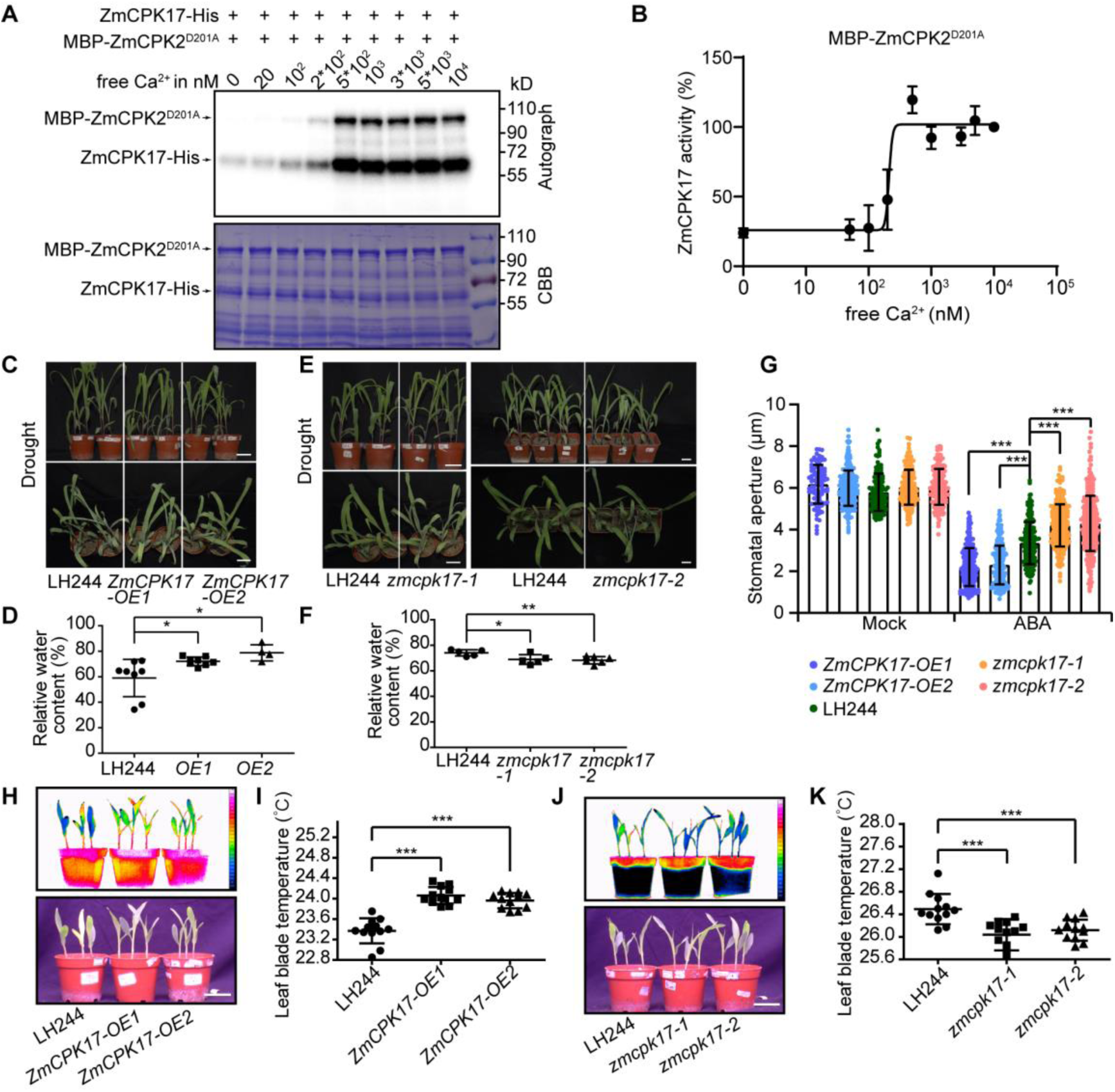
Calcium-dependent ZmCPK17 is a positive regulator for drought tolerance. **(A)** ZmCPK17 has calcium-dependent kinase activity. Recombinant ZmCPK17-His was subjected to the phosphorylation assay with recombinant kinase-dead MBP-ZmCPK2^D201A^ as the substrate in various free Ca^2+^ concentrations. The autograph and the Coomassie Brilliant Blue-stained gel (CBB) were shown. **(B)** Quantitation of the relative signal of MBP-ZmCPK2^D201A^ with Image J in **(A)** and analyzed by a Hill equation with a Hill factor of four in Graghpad Prism 8. The affinity toward Ca^2+^ of ZmCPK17 could be best described by the Hill equation. **(C)** Transgenic *ZmCPK17-OE1/2* seedlings are more tolerant to drought treatment compared with LH244. The pictures were taken from 14-d-old seedlings treated with drought stress. Bar = 5 cm. **(D)** Relative water content of leaves of *ZmCPK17-OE1/2* was calculated. Error bars represent ± SD from one experiment, and three experiments were performed with similar results. * represents significant difference with LH244, * p<0.05, Student’s *t*-test. **(E)** The CRISPR/Cas9 mutants *zmcpk17-1/-2* are more sensitive to drought treatment compared with LH244. The pictures were taken from 14-d-old seedlings treated with drought stress. Bar = 5 cm. **(F)** Relative water content of leaves of *zmcpk17-1/-2* was calculated. Error bars represent ± SD from one experiment, and three experiments were performed with similar results. * represents significant difference with LH244, * p<0.05, ** p<0.01, Student’s *t*-test. **(G)** Measurement of stomatal aperture with (ABA) or without (Mock) 12 µM ABA treatment. Three hundred stomata of three leaves from three seedlings were measured in one experiment, and three independent experiments were done with similar results. Data are means ± SD. * represents significant difference with LH244, *** p<0.001, Student’s *t*-test. **(H)** The leaf temperatures of *ZmCPK17-OE1/2* seedlings are higher than those of the wild type seedlings. 8-d-old seedlings with well watering were used for temperature measurement by a far-red CCD camera. **(I)** Leaf blade temperature in **(H)** was measured. Data are means ± SD of three replicates (eight leaves from one pot were measured per replicate) in one experiment. Four experiments were done with similar results. * represents significant difference with LH244, ***p < 0.001, Student’s *t*-test. **(J)** The leaf temperatures of *zmcpk17-1/-2* seedlings are lower than those of the wild type seedlings. 8-d-old seedlings with well watering were used for temperature measurement by a far-red CCD camera. **(K)** Leaf blade temperature in **(J)** was measured. Data are means ± SD of three replicates (eight leaves from one pot were measured per replicate) in one experiment. Four experiments were done with similar results. * represents significant difference with LH244, ***p < 0.001, Student’s *t*-test.

### ZmCPK17 inhibits ZmCPK2

Because ZmCPK17 and ZmCPK2 have an opposite role in drought stress, we speculated that ZmCPK17 may inhibit ZmCPK2 activity. We used the inactive ZmCPK2^D201A^, ZmCPK2^D201A^-NK, ZmCPK2-J as substrates, and found that ZmCPK17 phosphorylated all of them (Figure 4A). In contrast, ZmCPK2 did not phosphorylate the inactive form ZmCPK17^D173A^ (D173 is a putative active center site) (Figure 4B). We identified three phosphorylation sites at T60, T62 and T342 of ZmCPK2^D201A^ using LC-MS/MS (Figure S6A-C). ZmCPK17 phosphorylated ZmCPK2^D201AT60A^ and ZmCPK2^D201AT60AT62A^ to a less level than ZmCPK2^D201A^ and ZmCPK2^D201A^ ^T62A^ (Figure 4C), suggesting that T60 is a major phosphorylation site. Similarly, ZmCPK17 phosphorylated ZmCPK2^T342A^-J to a less level than ZmCPK2-J (Figure 4D). We found that ZmCPK2^T60A^ and ZmCPK2^T60D^ did not affect their phosphorylation activities on ZmYAB15, while ZmCPK2^T60A^ reduced a little autophosphorylation activity, and ZmCPK2^T60D^ increased its autophosphorylation activity (Figure 4E). ZmCPK2^T62A^ and ZmCPK2^T62D^ did not change phosphorylation activity (Figure 4E). Further analyses indicated that the phosphorylation activity of non-phosphorylable form ZmCPK2^T342A^ on MyBP was higher than ZmCPK2, and the phosphomimetic form ZmCPK2^T342D^ showed the lowest activity (Figure 4F). These results suggest that ZmCPK2 T342 is a key site inhibited by ZmCPK17 phosphorylation.

**Figure 4.**
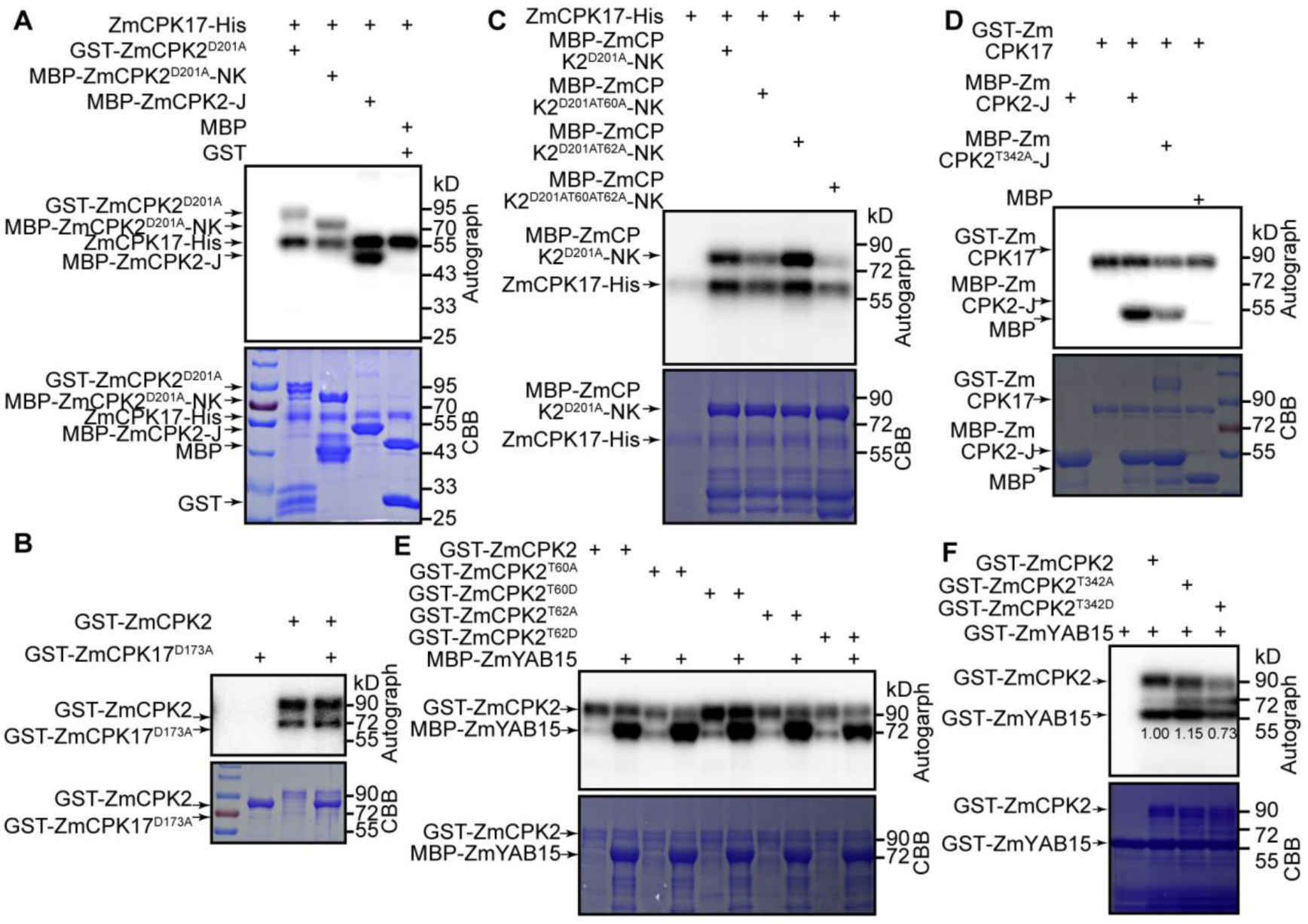
ZmCPK17 phosphorylates ZmCPK2 *in vitro*. **(A)** The full length and truncated forms of ZmCPK2 are phosphorylated by ZmCPK17. Recombinant ZmCPK17-His and the kinase dead GST-ZmCPK2^D201A^, MBP-ZmCPK2^D201A^-NK, MBP-ZmCPK2-J were subjected to phosphorylation assay. The empty GST and MBP were used as the negative substrates. The autograph and CBB were shown. **(B)** The full length ZmCPK17 is not phosphorylated by ZmCPK2. Recombinant GST-ZmCPK2 and the kinase dead GST-ZmCPK17^D173A^ were subjected to the phosphorylation assay. **(C)** ZmCPK2-NK T60 is one of key sites phosphorylated by ZmCPK17. Recombinant GST-ZmCPK17, MBP-ZmCPK2^D201A^-NK, MBP-ZmCPK2^D201A/T60A^-NK, MBP-ZmCPK2^D201A/T62A^-NK and MBP-ZmCPK2^D201AT60AT62A^-NK were subjected to the phosphorylation assay. **(D)** ZmCPK2-J T342 is one of key sites phosphorylated by ZmCPK17. Recombinant GST-ZmCPK17, MBP-ZmCPK2-J and MBP-ZmCPK2^T342A^-J were subjected to the phosphorylation assay. The empty MBP was used as a negative substrate. **(E)** Comparison of phosphorylation activities of ZmCPK2 T60A/D or T62A/D. The same amount of recombinant GST-ZmCPK2 with different phosphormimetic or non-phosphorylation forms (WT, T60A, T60D, T62A, T62D) were used for kinase activity assay with MBP-ZmYAB15 as the substrate. **(F)** The T342 phosphorylation negatively modulates the ZmCPK2 activity. The same amount of recombinant GST-ZmCPK2 with different mutant forms of T342 (WT, T342A or T342D) were used for kinase assay with GST-ZmYAB15 as the substrate.

In order to see whether ZmCPK17 affects ZmCPK2 activity in responding to ABA in plant cells, we first confirmed the dynamic activity of ZmCPK2 using *ZmCPK2-HF* maize protoplasts. Since we do not have ZmCPK17 antibodies and ZmCPK17-tagged transgenic plants, here we used protoplast transient assay in order to compare the results in a similar system. After treating protoplasts isolated from the 10-d-old *ZmCPK2-HF2* seedlings with 50 μM ABA for different time points, we extracted the total proteins and immunoprecipitated target protein with anti-Flag agarose, then conducted *in vitro* phosphorylation assay. The results showed that ABA treatment greatly inhibited the activity of ZmCPK2 at about 5 min, and then its activity was induced after 20 min, and gradually decreased later (Figure 5A). The quantitation of relative kinase activity was calculated using Image J. The dynamic change somehow exhibited a similar pattern with the observation using the maize seedlings as shown in Figure 2I.

**Figure 5.**
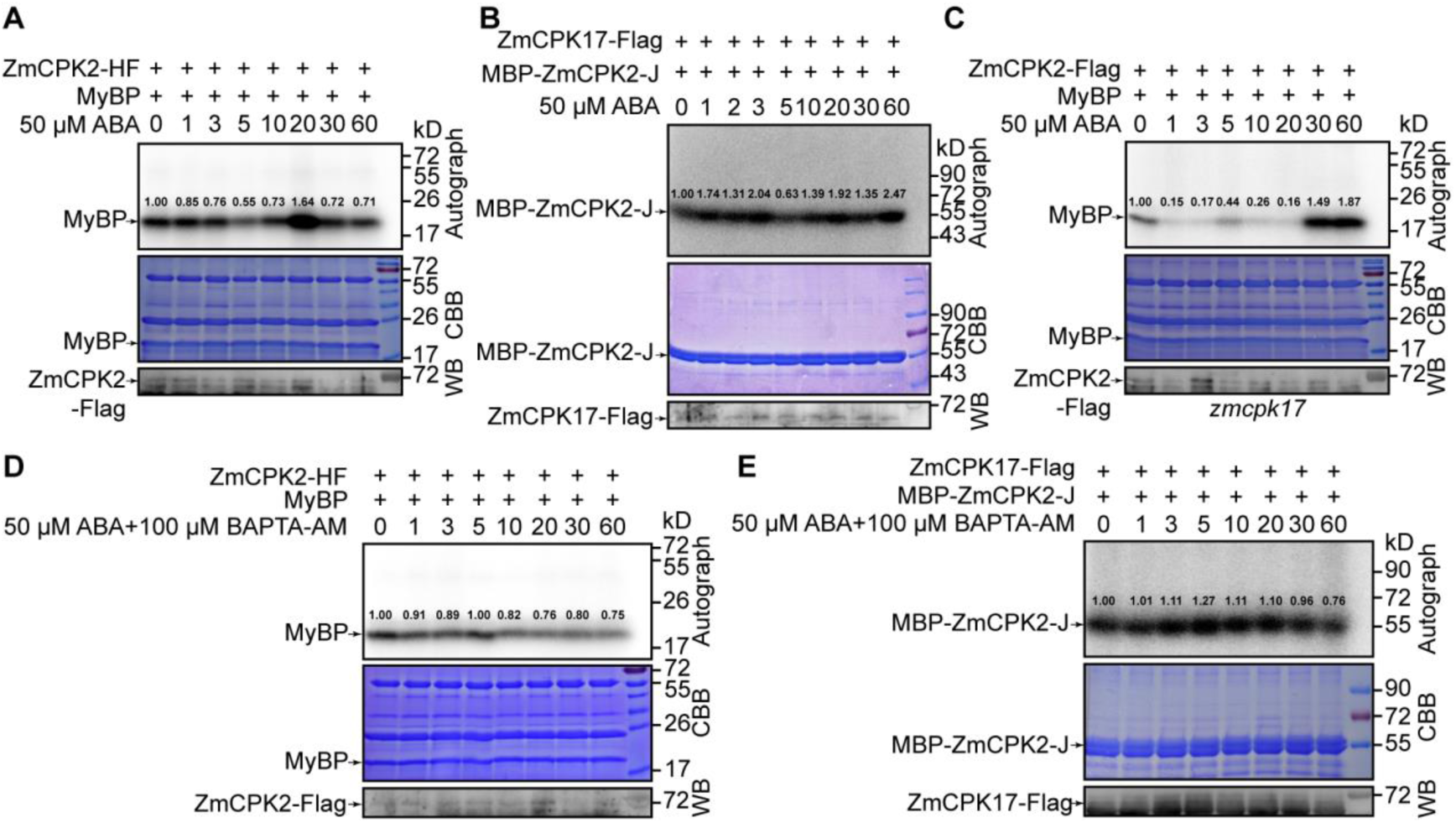
The activities of ZmCPK2 and ZmCPK17 are dynamically modulated by ABA. **(A)** The kinase activity of ZmCPK2 is dynamically modulated by ABA. The protoplasts were isolated from 10-d-old seedlings of *ZmCPK2-HF2,* and treated with 50 µM ABA for different times (0-60 min). ZmCPK2-HA-Flag was immunoprecipated (IPed) from total proteins with anti-Flag agarose. IPed proteins were subjected to *in vitro* phosphorylation assay using MyBP as a substrate. The autograph, CBB and the protein levels of ZmCPK2 in each sample (WB) were shown. The numbers represent the quantification of the phosphorylation signals of MyBP by the Image J software. **(B)** The kinase activity of ZmCPK17 is regulated by ABA. *Super: ZmCPK17-Flag* was transformed to maize protoplasts, which were treated with 50 μM ABA for different times (0-60 min). IPed proteins were subjected to *in vitro* phosphorylation assay using recombinant MBP-ZmCPK2-J as the substrate. The numbers represent the quantification of the phosphorylation signals of MBP-ZmCPK2-J by the Image J software. The autograph, CBB and the protein levels of ZmCPK17 in each sample (WB) were shown. **(C)** ABA-mediated ZmCPK2 activity in *zmcpk17*. *Super: ZmCPK2-Flag* was transformed to the protoplast isolated from 10-d-old seedlings of *zmcpk17*, which were treated with 50 μM ABA for different times (0-60 min). IPed proteins were subjected to *in vitro* phosphorylation assay using MyBP as the substrate. The numbers represent the quantification of the phosphorylation signals of MyBP. **(D)** The activity of ZmCPK2 is activated by ABA relying on calcium. The protoplasts were isolated from 10-d-old seedlings of *ZmCPK2-HF2,* and treated with 50 μM ABA and 100 μM BAPTA-AM (a cytosol calcium chelator) for different times (0-60 min). IPed proteins were subjected to *in vitro* phosphorylation assay using MyBP as the substrate. The numbers represent the quantification of the phosphorylation signals of MyBP. **(E)** ZmCPK17 activity is still activated by ABA without calcium. *Super: ZmCPK17-Flag* was transformed to maize protoplasts, which were treated with 50 μM ABA and 100 μM BAPTA-AM for different times (0-60 min). IPed proteins were subjected to *in vitro* phosphorylation assay using recombinant MBP-ZmCPK2-J as the substrate. The numbers represent the quantification of the phosphorylation signals of MBP-ZmCPK2-J.

We then measured the ZmCPK17 activity using a transiently expressing assay with 50 μM ABA treatment for different times. The expression vector carrying *Super: ZmCPK17-Flag* was transiently expressed in maize protoplasts. Then, ZmCPK17-Flag protein was immunoprecipiated and used for phosphorylation assay with MBP-ZmCPK2-J or general kinase substrate MyBP. Three independent experiments were performed with each substrate, and similar results were obtained (Figure 5B, Figure S7C, S7D, S7E). We noticed that although the degree of gene expression was somehow different at different time points, the changing tendency was similar. Similarly, we found that ZmCPK17 activity was also dynamically changed with ABA treatment in a time course: quickly induced at 1 - 3 min (please see three independent results for both MBP-ZmCPK2-J and general kinase substrate MyBP), then reduced at about 3-5 min, and induced again at 10 - 20 min, reduced at 20 - 30 min, then recovered again at 60 min. We then checked ZmCPK2 activity in *zmcpk17* mutant, and found that ZmCPK2 activity in *zmcpk17* mutant was reduced to the lower level at 1 - 5 min, then recovered to higher level after 30 - 60 min (Figure 5C, S7F, S7G). According to the quantitation of activity, we could notice that the dynamic pattern of ZmCPK2 activity seems likely opposite to that of ZmCPK17 during ABA treatment except for the early time point (Figure 5A, 5B). In the *zmcpk17* mutant, the induction peak time of ZmCPK2 activity at 10 - 30 min was disappeared, and the inhibition of ZmCPK2 was not observed at 30 - 60 min (Figure 5C), suggesting that dynamic pattern of ZmCPK2 activity is mediated by ZmCPK17. Meanwhile, we treated transiently expressing protoplasts with 50 μM ABA plus 100 μM BAPTA-AM, a cytosolic Ca^2+^ chelator (Garcia-Mata and Lamattina, 2007), for different times, and immunoprecipated proteins were used for phosphorylation assay. We found that dynamic patterns of both ZmCPK2 and ZmCPK17 activity were disturbed when Ca^2+^ was not available (Figure 5D, S7H, S7I, and 5E, S7J and S7K). We then used maize protoplasts transiently expressing the calcium reporter aequorin under the control of the *UBQ10* promoter to monitor the Ca^2+^ signature during ABA treatment (Ma et al., 2019; Mehlmer et al., 2012). When adding 100 μM BAPTA-AM to chelate cytosol Ca^2+^, we did not observe a clear Ca^2+^ change, indicating that the Ca^2+^ signature is imposed by ABA. However, we found that ABA treatment increased the cytosol Ca^2+^ to a high level at different times, but we could not find a defined or regular mode, probably because we used the transiently expressed protoplasts that are not uniform (Figure S7L). Thus, we quantified relative changes in cytoplasmic Ca^2+^ by calculating 20 peaks induced by three independent treatments versus 20 peaks in the control. The results showed that ABA significantly induced [Ca^2+^]_cyt_ (Figure S7M). Although ZmCPK2 is a Ca^2+^ independent kinase, its activity did not show any change when the protoplasts were treated with ABA and BAPTA-AM together (Figure 5D, S7H, S7I), suggesting that its dynamic activity indirectly relies on Ca^2+^ signaling. Although ZmCPK17 lost dynamic pattern in ABA combined with BAPTA-AM treatment, its activity was still activated and reached to a high level at about 5 min and to normal level at 60 min (Figure 5E, S7J, S7K), suggesting that both Ca^2+^ and ABA signaling modulate its activity. These results further indicate that ZmCPK17 modulates ZmCPK2 activity. As the clade A PP2Cs are negative factors that could directly interact with and inhibit some protein kinases such as CPKs (Chen et al., 2021; Geiger et al., 2010), we checked whether ZmCPK17 and ZmCPK2 are inhibited by some PP2Cs in maize. Yeast-two hybrid assay indicated that ZmCPK17 interacted with ZmPP2C3, 6, 9, 11, 12, and 15 (Figure S8A), and ZmCPK2 interacted with ZmPP2C11, 12, and 15 (Figure S8B), while ZmPP2C9, 11, 12, 15 interacted with ZmPLY8 in an ABA-dependent manner (Figure S8C). Thus, we chose ZmPP2C11 for further *in vitro* study. ZmCPK17 and ZmCPK2 were inhibited by ZmPP2C11, or ZmPP2C11 plus ZmPYL8, but not by ZmPYL8. However, addition of ABA released the inhibition of ZmPP2C11 plus ZmPYL8 on ZmCPK17 and ZmCPK2 (Figure S8D, S8E), suggesting that ZmCPK17 and ZmCPK2 can be both activated by releasing the inhibition of ZmPP2C11 through ABA-bound ZmPYL8. These results suggest that ZmCPK17 inhibits ZmCPK2 activity under ABA treatment in an opposite dynamic manner, which at least partially relies on Ca^2+^ signaling. Considering the change of Ca^2+^ oscillation during stress response, ZmCPK17 may be the output of Ca^2+^ signalling.

### ZmCPK17 affects the nuclear localization of ZmCPK2

Given that ZmCPK17 can inhibit ZmCPK2, we further investigated whether ZmCPK17 also influences the localization of ZmCPK2 using maize transiently expressing protoplast assay. ZmCPK2-, ZmCPK2^T60A^- or ZmCPK2^T60D^-GFP was found to locate in both cytoplasm and nucleus, while ZmCPK17-GFP mainly in the cytoplasm (Figure S9A). The AtMCM10 (Minichromosome maintenance protein 10)-mCherry was used as a nuclear location marker (Figure S9A) (Zhao et al., 2023). We also compared the localization intensity of ZmCPK2-GFP, and phosphomimetic form ZmCPK^T60D^-GFP, and non-phosphomimetic form ZmCPK2^T60A^-GFP in nucleus and cytoplasm, and found that ZmCPK^T60D^-GFP seemly reduced GFP signal in nucleus (Figure S9A, S9I). ABA treatment did not show a clear difference for the localization of either ZmCPK2-GFP or ZmCPK17-GFP (Figure S9B). ZmCPK2-GFP and ZmCPK17-mCherry co-located in the cytoplasm when they were transiently co-expressed in protoplasts (Figure S9C, S9F). Interestingly, the nuclear localization of ZmCPK2-GFP was significantly reduced when co-expressed with ZmCPK17-mCherry (Figure S9C, S9I). Under ABA treatment, we did not observe a clear difference of ZmCPK2 distribution when co-expressed with ZmCPK17, probably because ZmCPK2 or ZmCPK17 was quietly overexpressed, which could not be affected by ABA-enhanced ZmCPK17 or the basal activity of overexpressed ZmCPK7 is enough to fully phosphorylate ZmCPK2 (Figure S9C, S9F). When co-expressed with ZmCPK17-mCherry, the nuclear localization intensity of ZmCPK2^T60D^-GFP was reduced to a similar level as ZmCPK2-GFP, but ZmCPK2^T60A^-GFP was not apparently changed (Figure S9D, S9G, S9I), suggesting that phosphorylation of ZmCPK2 by ZmCPK17 reduces its nuclear localization. To verify whether the effect of ZmCPK17 on ZmCPK2 localization depends on its kinase activity, we co-expressed the kinase-dead form ZmCPK17^D173A^ (D173 is the active center site) with ZmCPK2, and found that the nuclear localization of ZmCPK2 was almost not affected (Figure S9E, S9H, S9I), suggesting that ZmCPK17 preventing the nuclear localization of ZmCPK2 requires its kinase activity. These results suggest that besides inhibiting ZmCPK2, ZmCPK17 can reduce ZmCPK2 nuclear localization through phosphorylation, thus further alleviating the roles of ZmCPK2 in nuclei.

### ZmYAB15 is a direct target of ZmCPK2

In order to identify the substrates of ZmCPK2, we used the kinase domain of ZmCPK2 (ZmCPK2-NK) as a bait to perform yeast two-hybrid screening from a maize cDNA library, and identified ZmYAB15 as a putative interaction protein. ZmYAB15 belongs to YABBY family. The yeast two-hybrid assay confirmed that ZmCPK2 interacted with ZmYAB15 (Figure S10A). Here, OST1 and ABI1 were used as a positive control (Deng et al., 2022). Next, we used the purified proteins isolated from maize protoplasts, performed Co-IP assay, and verified the interaction between ZmCPK2 and ZmYAB15 *in vivo* (Figure S10B). BiFC assay further demonstrated that ZmCPK2 interacted with ZmYAB15 in cytoplasm and nucleus (Figure S10C). We used GUS-YCE and GUS-YNE as negative controls. Interestingly, we also found that ZmCPK17 interacted with ZmYAB15 in yeast two-hybrid assay (Figure S10D), and ZmCPK17-YCE could also interact with ZmYAB15-YNE in both nucleus and cytoplasm in BiFC assay (Figure S10C). We further performed pull-down assay and found that ZmYAB15 interacted with ZmCPK17, which was not affected by Ca^2+^ (Figure S10E). ZmYAB15-GFP was well co-localized with ZmCPK2-mCherry in both nucleus and cytoplasm (Figure S10F), but only partially or weakly with ZmCPK17-mCherry mainly in cytoplasm (Figure S10G). We performed *in vitro* phosphorylation assay, and found that both ZmCPK2 and ZmCPK17 could phosphorylate ZmYAB15 (Figure 2B, S10H). To explore whether ZmCPK2 and ZmCPK17 phosphorylated different domains of ZmYAB15, we constructed the three domains of ZmYAB15, the Zinc finger domain (zf), the disordered domain (dis) and the HMG domain (hmg), into the recombinant vector to perform the phosphorylation assay. The results showed that ZmCPK2 only phosphorylated ZmYAB15-dis, ZmCPK17 could phosphorylate ZmYAB15-dis and ZmYAB15-hmg (Figure S10I). Because it is difficult to differentiate the phosphorylation ability imposed by ZmCPK2 or ZmCPK17 when combining two kinase together, we then co-expressed ZmCPK17 with ZmCPK2 in *E. coli* to see whether ZmCPK17 could affect the kinase activity of ZmCPK2. Indeed, we found that ZmCPK2 purified from *E. coli* co-expressing ZmCPK17 phosphorylated ZmYAB15 to a less level than that from *E. coli* only expressing ZmCPK2 (Figure S11A), which was consistent with the above results that ZmCPK17 inhibited the kinase activity of ZmCPK2. However, we found that when truncated form ZmCPK2-NKJ or ZmCPK2-NK was co-expressed with ZmCPK17, respectively, both of the purified truncated ZmCPK2 forms phosphorylated ZmYAB15 to a higher level than only expressing ZmCPK2-NKJ or ZmCPK2-NK (Figure S11B, S11C). These results suggest that ZmCPK2 C-terminus is an inhibition domain that can be further inhibited by ZmCPK17 phosphorylation, while C-terminal truncated forms might alter their structures, possibly favoring ZmCPK17 phosphorylation on ZmCPK2 N-terminus, thereby, activating rather than inhibiting their activity, which may explain why the two mutants *zmcpk2-3/-4* with deletion of 4 EF hands exhibited a similar drought sensitive phenotype as the OE lines. Meanwhile, ZmCPK17^D173A^ had no influence on the kinase activity of ZmCPK2 (Figure 4B, S11D). Considering that ZmCPK17 could reduce the nuclear location of ZmCPK2 through phosphorylation on T60, we detected the interaction between ZmYAB15 and phosphomimetic ZmCPK2 T60D or non-phosphomimetic ZmCPK2 in pull-down assay and yeast two-hybrid assay. The results consistently showed that different phosphorylation forms of T60 in ZmCPK2 had no obvious influence on their interaction (Figure S11E, S11F). Similarly, different phosphorylation forms of ZmCPK2 T342 did not influence on their interaction with ZmYAB15 (Figure S11G). Consistent with above mentioned results, these data suggest that ZmCPK17 could inhibit the activity of ZmCPK2 through phosphorylation, and ZmCPK17 also regulates ZmYAB15 in an ZmCPK2-independent way. ZmYAB15 should one of ZmCPK2 or ZmCPK17 targets in maze drought response.

### ZmYAB15 negatively regulates drought response

To further investigate that ZmYAB15 is one of candidate substrates of ZmCPK2, we obtained the overexpression transgenic lines and CRISPR/Cas9 mutants of *ZmYAB15* to analyze their drought phenotypes. The soil drought treatment of seedlings assay showed that *ZmYAB15-OE* displayed drought sensitive phenotype (Figure 6A). Detached leaf water loss confirmed the more water loss of *ZmYAB15-OE1/2* compared to the wild type (Figure 6B). Both *ZmYAB15-OE1* and *-OE2* had higher expression level of *ZmYAB15* comparing to the wild type as checked by qRT-PCR (Figure 6C). We obtained two *zmyab15* mutants with CRISPR/Cas9 technique. One mutation with a T insertion and another with an A insertion after T207 counting from the first putative start codon A (1)TG, both of which produce a frame shift after Leu69 (Figure S12A). Consistently, *zmyab15-1/-2* mutants displayed drought tolerant phenotype with higher relative leaf water content comparing with the wild type (Figure 6D, 6E). When the mutant, OE and wild type seedlings were planted in the same pots for drought stress treatment, *zmyab15-1/-2* showed most tolerant to drought stress, while *ZmYAB15-OE1/2* were most sensitive to drought stress (Figure 6F). We also compared the leaf temperatures and found that *ZmYAB15-OE2* seedlings had lower temperatures, while *zmyab15-1* mutant had higher temperatures than the wild type seedlings (Figure 6G, 6H), indicating that *ZmYAB15-OE2* transpires more water, while *zmyab15-1* transpires less water than the wild type. We also performed stomatal aperture assay and found the stomatal opening of both *ZmYAB15-OE1/2* and *zmyab15-1/-2* had no significant difference with LH244 without ABA treatment (Mock), while *ZmYAB15-OE1/2* exhibited less sensitivity to ABA-induced stomatal closure and *zmyab15-1/-2* more sensitivity comparing with the wild type (Figure 6I). We did not observe the difference in plant growth and development among these OE lines, mutants and the wild type seedlings under normal growth conditions (Figure S12B, S12C). We also observed the field drought treatment phenotype, and found that *ZmYAB15-OE* displayed more wilting phenotype in young leaves and earlier senescence in old leaves compared to LH244 (Figure S12D). From the field data, we found that *zmyab15* was shorter than LH244 (8.12%) in the height (Figure S12E), and the hundred grain weight was similar to LH244 (Figure S12F). Thus, ZmYAB15 negatively modulates drought response.

**Figure 6.**
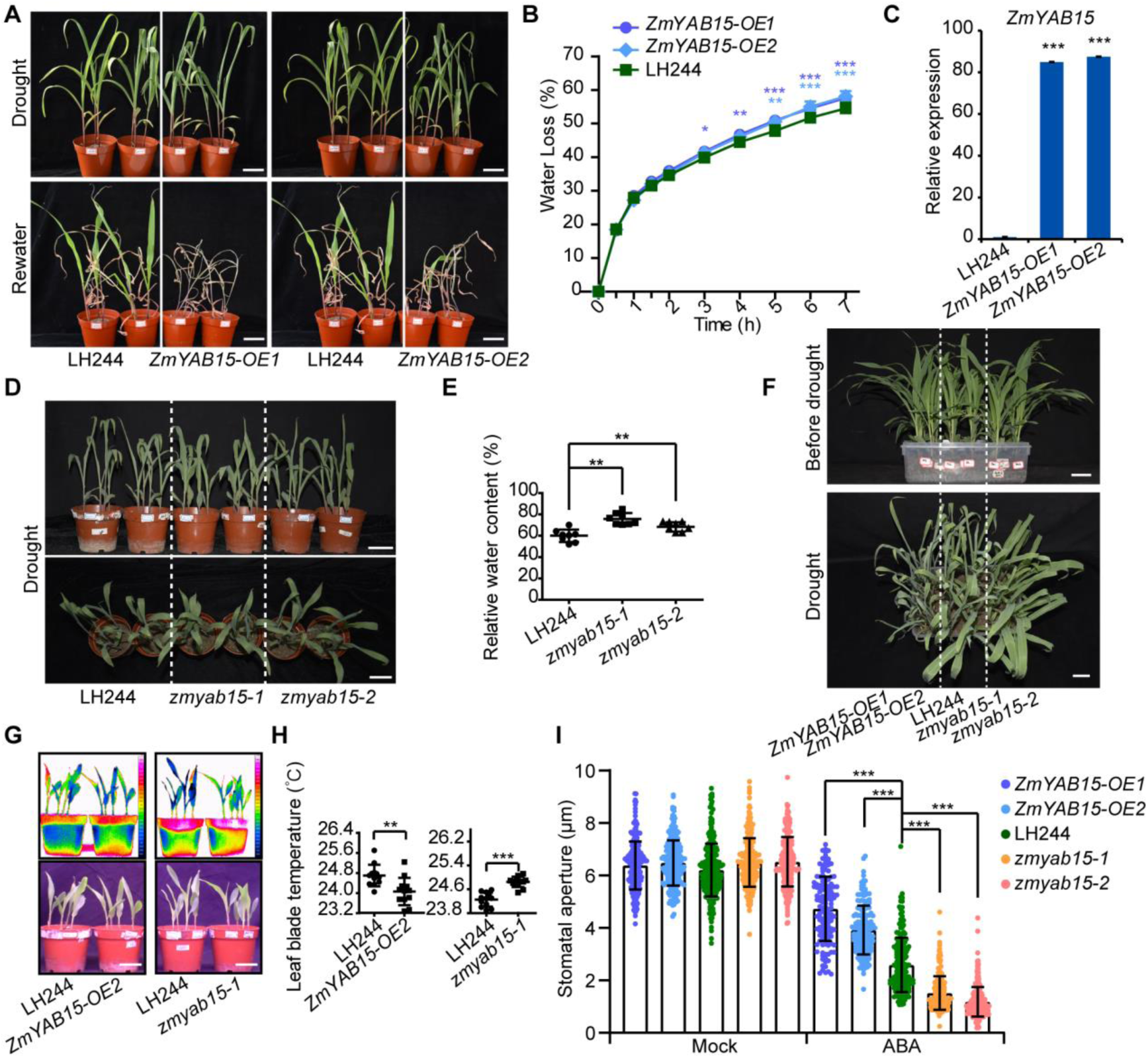
ZmYAB15 is a negative drought regulator. **(A)** Transgenic *ZmYAB15-OE1/2* seedlings are more sensitive to drought treatment compared with LH244. Maize seedlings were treated with drought stress and then rewatered. Bar = 5 cm. **(B)** Water loss of *ZmYAB15-OE1/2* and LH244. Error bars represent ± SD of three technical replicates from 12 second expanded leaves in one representative experiment. * Represents significant difference with LH244, * p<0.05, ** p<0.01, *** p<0.001, Student’s *t*-test. **(C)** Relative expression level analyses of *ZmYAB15* in transgenic *ZmYAB15-OE1/2* plants and LH244. Total RNAs were extracted from 9-d-old well-watered seedlings. Error bars represent ± SD of three technical replicates from one experiment and three experiments were performed with similar results. * Represents significant difference with LH244, *** p<0.001, Student’s *t*-test. **(D)** The CRISPR/Cas9 mutants *zmyab15-1/-2* are more tolerant to drought treatment compared with LH244. Pictures were taken from 12-d-old seedlings treated with drought stress. Bar = 5 cm. **(E)** Relative water content of leaves was calculated in **(D)**. The fresh weight, saturated fresh weight and dry weight of all cut leaves were measured for calculating the relative relative water. Each point represents a group of leaves from one seedling. Error bars represent ± SD from one experiment and three experiments were performed with similar results. * Represents significant difference with LH244, ** p<0.01, Student’s *t*-test. **(F)** Transgenic *ZmYAB15-OE1/2* seedlings are more sensitive and the CRISPR/Cas9 mutants *zmyab15-1/2* are more tolerant to drought treatment compared with LH244. All of the plants were planted in one tray and stopped water supply at 8-d-old. Bar = 5 cm. **(G)** The leaf temperature comparison of *ZmYAB15-OE2*, *zmyab15-1* and wild type. 8-d-old seedling with well watering were used for temperature measurement by a far-red CDD camera. **(H)** Statistical analysis of leaf temperature in **(G)**. Data are means ± SD of three replicates (eight leaves from one pot were measured per replicate) in one experiment. Four experiments were done with similar results. Asterisks indicate a significant difference compared with the wild type: **p < 0.01, ***p < 0.001, Student’s *t*-test. **(I)** Measurement of stomatal aperture with (ABA) or without (Mock) 12 µM ABA treatment. Three hundred stomatas of three leaves from three seedlings were measured in one experiment, and three independent experiments were done with similar results. Data are means ± SD. * represents significant difference with LH244, *** p<0.001, Student’s *t*-test.

In order to find the genes that are modulated by ZmYAB15, ZmCPK2 and ZmCPK17, we performed RNA sequencing experiments to profile the transcriptome of the *ZmCPK17-OE2*, *zmcpk2-1*, *zmyab15-1* and LH244 under normal condition and ABA treatment for 3 hours. We used *ZmCPK17-OE2* because *ZmCPK17* is a positive regulator of drought stress, and other two genes are negative regulators, thus we may identify their commonly regulated genes. The transcription levels of *ZmCPK2* and *ZmYAB15* were not regulated by ABA while *ZmCPK17* were a little upregulated after ABA treatment (3 h). *ZmNAC111* was used as ABA-induced marker gene (Figure S13A) (Mao et al., 2015). Total RNAs were extracted from the second leaves of 8-d-old seedlings and used for mRNA-seq library construction. We collected the leaves from three seedlings for each sample and sequenced two biological replicates on a NovaSeq 6000 platform (Illumina) and obtained more than 30 million paired-end clean reads for each replicate. RNA-seq reads were mapped back to the reference maize genome (B73 RefGen_v4, AGPv4). Differentially expression genes (DEGs) were selected with a false discovery rate (FDR) < 0.05. After 3 h ABA treatment, we identified 3286 DEGs in *ZmCPK17-OE2*, 2830 DEGs in *zmcpk2-1*, 3237 DEGs in *zmyab15-1* comparing with the wild type (Figure 7A). The expression of about 1070 genes was only changed in the wild type, not in three mutants (Figure 7A). 103 genes were co-regulated only in *ZmCPK17-OE2*, *zmcpk2-1* and *zmyab15-1* (Figure 7A), which include 60 co-upregulated genes and 32 co-downregulated genes.

**Figure 7.**
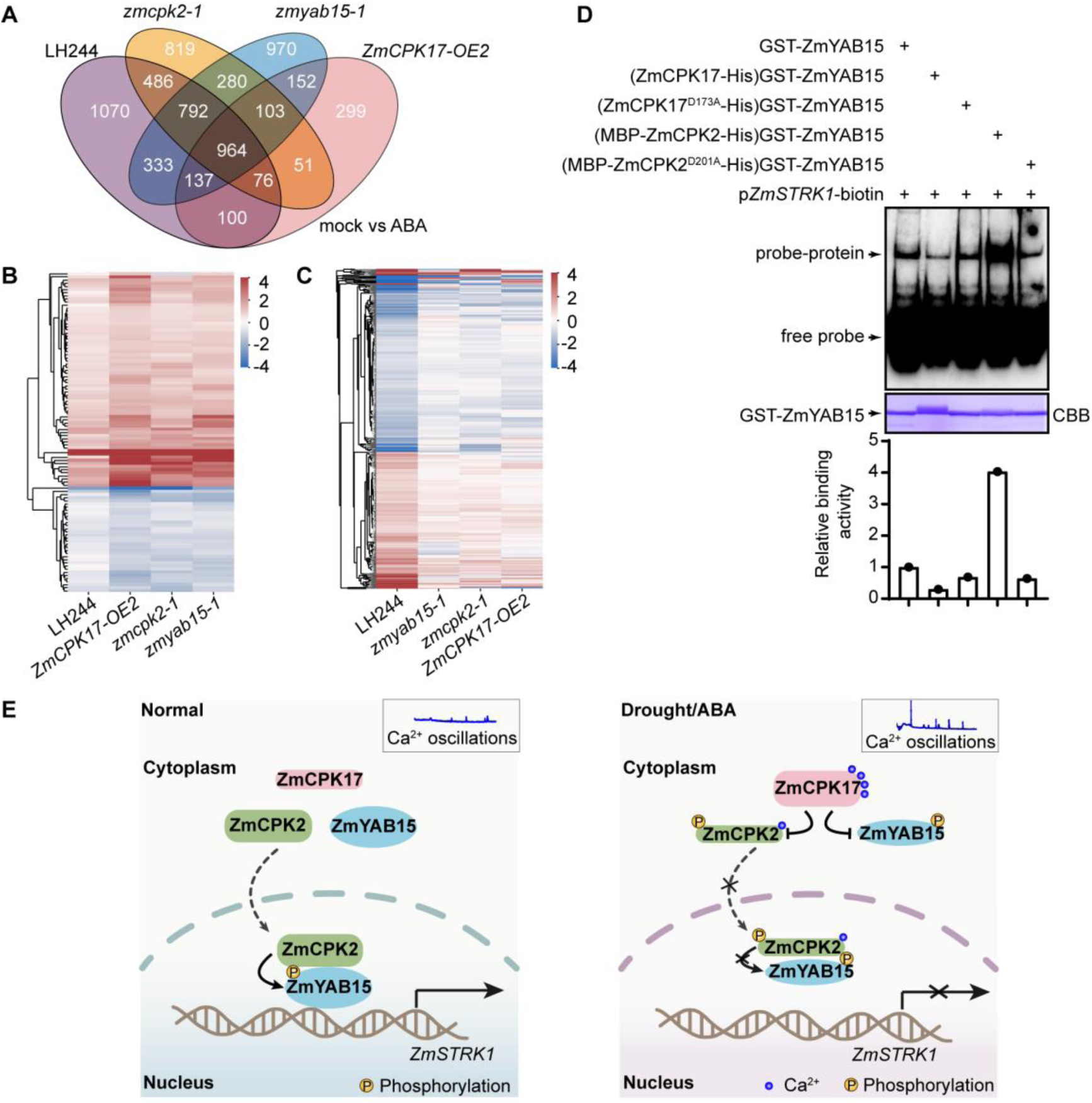
ZmCPK17 plays contrast roles with ZmCPK2 and ZmYAB15 in drought stress. **(A)** A Venn diagram shows the number of overlapped different expression genes in 9-d-old seedlings of LH244, *zmcpk2-1*, *zmyab15-1* and *ZmCPK17-OE2* treated with 50 µM ABA compared with no ABA treatment (mock). **(B)** The heat map shows the 103 genes only significantly changed in *ZmCPK17-OE2*, *zmcpk2-1* and *zmyab15-1*. The unit of expression is FPKM. **(C)** The heat map shows the 1070 genes only significantly changed in LH244 but not in *ZmCPK17-OE2*, *zmcpk2-1* and *zmyab15-1*. The unit of expression is FPKM. **(D)** The phosphorylation of ZmCPK17 weakens and ZmCPK2 enhances the DNA binding activity of GST-ZmYAB15 on the promoter of *ZmSTRK1* in EMSA assay. Recombinant ZmCPK17-His or MBP-ZmCPK2-His with or without kinase activity were co-expressed with GST-ZmYAB15 in the *E. coli* strain BL21 (DE3). GST-ZmYAB15 was purified from five different transforms and performed the EMSA assay. The protein amount was shown as CBB. The relative binding activity was measured by Image J and shown under the CBB. **(E)** A proposed model ZmCPK17-ZmCPK2-ZmYAB15 module for drought response. Under normal condition (Normal), ZmCPK2 interacts with and phosphorylates ZmYAB15 and could enhance the DNA binding activity of ZmYAB15 to activate *ZmSTRK1* expression; when drought or ABA triggers the Ca^2+^ oscillations (Drought/ABA), ZmCPK17 is activated and then phosphorylates and inhibits ZmCPK2, resulting in inactivating ZmYAB15 due to reduced phosphorylation by ZmCPK2, and/or increased phosphorylation by ZmCPK17, which would trigger drought stress response.

Further GO analysis showed these 103 co-regulated genes are involved in water and chemical response, oxoacid metabolic and single-organism process and so on (Figure S13B). We then compared the expression level of these co-regulated genes and made heat map analysis. Interestingly, we found that the expression levels of these co-upregulated genes in mutants were all higher and co-downregulated genes in mutants were all lower than LH244 after ABA treatment (Figure 7B). The upregulated genes are negatively regulated by ZmCPK2 and ZmYAB15, and positively regulated by ZmCPK17 under ABA treatment. These results strongly suggest that ZmCPK2 and ZmYAB15 play contrast roles with ZmCPK17 in regulating the expression of these common genes. We also compared the expression levels of 1070 genes only changed in LH244, and found that the expression levels of ABA-downregulated genes are lower in the wild type plants than mutants, while the expression levels of ABA-upregulated genes are higher in the wild type than mutants after ABA treatment (Figure 7C). GO analysis of these genes showed that they are involved in various processes, especially in biosynthetic processes (Figure S13C). These results suggest that one of main outputs imposed by these proteins under ABA treatment is the change of various metabolic processes.

Previous studies have reported that ZmYAB15 belongs to the YABBY transcriptional factor family (Juarez et al., 2004) and there were four predicted binding motifs of YABBY (Figure S13D) (Franco-Zorrilla et al., 2014; Shamimuzzaman and Vodkin, 2013). We found that ZmYAB15 could bind probe 2 and 4 that are rich in C (Figure S13E). We selected several candidate genes from the 103 genes, in which their promoters contain some putative YABBY binding cis-elements. We found that ZmYAB15 could weakly bind to the prompter of the co-upregulated gene *RD22* (BURP domain 22), and strongly bind to the co-downregulated gene *ZmSTRK1* (Salt Tolerance Receptor-like cytoplasmic Kinase 1), but not bind to the promoter of *HAK11* (High-affinity K^+^ transporter 11) (Figure S13F, S13G) in EMSA assay. ZmSTRK1 belongs to receptor-like cytoplasmic kinase family and is homologous to OsSTRK1 that positively regulates salt and oxidative stress tolerance (Vij et al., 2008; Zhou et al., 2018). We confirmed the expression of *ZmSTRK1* was inhibited by ABA treatment using qRT-PCR (Figure S13A). We further explored whether the phosphorylation of ZmYAB15 by ZmCPK2 or ZmCPK17 affects its binding activity and/or transcriptional activity. By co-expressing ZmYAB15 and ZmCPK2 or ZmCPK17 in *E. coli*, we found that ZmCPK17 inhibited the binding activity of ZmYAB15, while ZmCPK2 significantly activated the binding activity of ZmYAB15 in EMSA assay, while the kinase dead forms of ZmCPK17 and ZmCPK2 had no clear influence on ZmYAB15 binding ability (Figure 7D). Further dual-LUC assay with transiently expressing *pZmSTRK1:LUC* reporter and ZmYAB15-Flag effector in *N. benthamiana* leaves showed that ZmYAB15 activated the expression of *pZmSTRK1:LUC*, while the activation was enhanced in the presence of ZmCPK2 and a little inhibited by ZmCPK17 (Figure S13H, S13I). Meanwhile, the kinase dead forms of ZmCPK2 and ZmCPK17 had no influence on ZmYAB15 (Figure S13I). These results suggest that ZmYAB15 can directly activate *ZmSTRK1* expression, while ZmCPK2 phosphorylation can promote its binding activity, and ZmCPK17 phosphorylation can inhibit its binding activity. In the future study, the genetic analysis of ZmYAB15 with ZmCPK2 and ZmCPK17 is required to confirm their relationship in planta.

## DISCUSSION

As a key second message signal, Ca^2+^ plays multifaced roles in plants responding to both abiotic and biotic stresses (Dong et al., 2022). Many different Ca^2+^ binding proteins such CPKs have been extensively studied for their roles in drought stress in the past years, but the regulation mechanism among different CPKs is not well known (Dong et al., 2022). It has reported that many CPKs involved in the ABA-regulated stomatal movement with the stomatal apertures assay in excised leaves (Brandt et al., 2015; Li et al., 2022a; Mori et al., 2006). Ca^2+^ oscillations in different cell types or under different stress conditions has been known for a long time (Allen et al., 2000; Tian et al., 2020). A recent study used a powerful approach CDPK-FRET for tackling real-time live-cell decoding, and found that ABA induces an increase in cytosolic Ca^2+^-concentrations in the form of repetitive transients, and triggers a conformational change in Ca^2+^-sensitive AtCPK21 (Liese et al., 2023). However, whether Ca^2+^ oscillation would synergistically trigger downstream target oscillation is still not known. In this study, we propose a model for how Ca^2+^-dependent ZmCPK17 mediates Ca^2+^-independent ZmCPK2 activity in an dynamic manner for plants responding to drought stress and ABA signaling in maize, which may match the Ca^2+^ oscillation patterns in cells.

We found that ZmCPK2 physically interacts with ZmCPK17. The activity of ZmCPK2 was not influenced by supplying or depleting Ca^2+^, indicating that its activity is not dependent on Ca^2+^, while calcium could activate ZmCPK17. ZmCPK17 was able to phosphorylate and inhibit ZmCPK2, but ZmCPK2 could not phosphorylate ZmCPK17, indicating that ZmCPK2 is one of ZmCPK17 downstream direct targets. Under different concentrations of ABA treatment, cytosolic Ca^2+^ concentration would be triggered to reach a high level in a short time, then display prolonged oscillations (Allen et al., 2000; Dong et al., 2022; Islam et al., 2010), but the biological significance is not well known. ZmCPK17 activity could be induced at 1-3 min after ABA treatment in maize protoplast cells. More interestingly, ZmCPK17 activity was then dynamically changed: reduced to a lower level at 3 - 5 min, increased to a higher level at 10 - 20 min, reduced to a much lower level at 20 - 30 min, and then recovered to basal level at 60 min, which showed the similar form of repetitive transient conformation changes of AtCPK21 in *Arabidopsis* guard cells responding to ABA and the flagellin peptide flg22, which rely on Ca^2+^ oscillations (Liese et al., 2023). Correspondingly, ZmCPK2 activity in maize seedlings reduced to a lower level at 5 min and increased to a higher level than basal at 10 - 30 min, then reduced to a low level again at 30 - 90 min, suggesting that with short and long time ABA treatment, ZmCPK2 activity was more inhibited. It seems that ZmCPK17 activity is oppositely correlated with ZmCPK2 activity in a time delayed manner. It is very possible that ZmCPK17 directly inhibits ZmCPK2 activity *in vivo*. In a previous study on *Arabidopsis* guard cells, it was found that Ca^2+^ oscillation had different amplitudes and frequencies under different treatment conditions (Allen et al., 2000), which might also occur in different plants or cells. We found that the earliest activation of ZmCPK17 was observed at 1 - 3 min in protoplasts after 50 μM ABA treatment, and the correspondingly reduced ZmCPK2 activity was observed at about 5 min in seedlings and protoplasts (Figure 2I, 5A). It was speculated that the response of ZmCPK2 was slower than ZmCPK17. However, in *zmcpk17* loss-of-function mutant cells, we observed that ZmCPK2 activity was reduced to a lower level in 5 - 20 min, and the earliest activation of ZmCPK2 at 10 - 30 min after ABA treatment was disappeared, then the activity recovered to basal level after 30 min, and did not reduce any more at 60 min. These observations suggest that ZmCPK2 activity is inhibited at both medium and prolonged treatment by ZmCPK17. However, as a negative regulator for drought stress, how ZmCPK2 can be activated to a higher level than its basal activity is a very interesting observation. Also, the earliest inhibition of ZmCPK2 in the first 5 min still occurred in *zmcpk17* (Figure 5C). It is very likely that other unidentified factors should be involved in this process during Ca^2+^ oscillation of drought stress. ZmCPK2 was found to putatively interact with different ZmCPKs in yeast-two hybrid assay (Figure S3C). Although we did not further analyze these proteins due to the conditional limitation, it is possible that some of these putative proteins or other unidentified ones might play a role during early drought stress and/or ABA signaling together with ZmCPK2 and ZmCPK17. For example, ZmCPK2 might be activated or inhibited by other calcium-dependent CPKs, which needs further explored. These results suggest that ZmCPK17 and ZmCPK2 activity are dynamically changed likely accompanying with Ca^2+^ oscillations during ABA treatment. Although most of studies utilize guard cells as model system to study calcium signatures under different stress conditions (Liese et al., 2023), other cells most likely exhibit a similar responsive pattern as guard cells as early suggested (Schroeder et al., 2001). We suspect that dynamic changes of kinase activity might depend on Ca^2+^ oscillation and/or different regulation components, which would endow plant adaptation to drought stress more efficiently: suddenly braking or speeding up the signaling may not be good to plants.

The localization of a protein is crucial for its functions. ZmCPK17 is largely localized in the cytosol and ZmCPK2 in both cytosol and nucleus. However, we found that when co-expressing ZmCPK2 and ZmCPK17, ZmCPK2 was more detected in cytosol and co-localized with ZmCPK17, indicating that ZmCPK17 influences the localization of ZmCPK2. When the phosphorylation site ZmCPK2 T60 was mutated to A60, the unclear localization of non-phosphomimetic ZmCPK2^T60A^ was not further reduced when co-expressed with ZmCPK17. We found that ZmCPK17 had basal kinase activity in the transient protoplast assay, which might be high enough for phosphorylating ZmCPK2 for its retention in cytoplasm when ZmCPK17 was overexpressed. Thus, ABA-activated ZmCPK17 may not only interact with, phosphorylate and inhibit ZmCPK2, but also reduce ZmCPK2 nuclear localization, attenuating its effects in nucleus and further reducing its roles in nucleus.

We found that ZmCPK17 and ZmCPK2 could physically interact with and phosphorylate ZmYAB15 so as to regulate the gene expression of *ZmSTRK1*. Transgenic plants overexpressing either *ZmCPK2* or *ZmYAB15* were more sensitive while *zmcpk2-1* and *zmyab15* mutants were more resistant to drought stress and ABA than the wild type plants, while *ZmCPK17* exhibited oppositely, indicating that both ZmCPK2 and ZmYAB15 are negative regulators, and ZmCPK17 is a positive regulator in drought tolerance. YAB transcriptional factors have been found to be involved in plant growth, development and stress responses, but most of studies focused on plant growth and development (Zhang et al., 2020). For example, Shattering1 (Sh1) in sorghum and its two orthologs (ZmSh1-1 and ZmSh1-5.1+ZmSh1-5.2) in maize, one ortholog SH1 in foxtail millet (*Setaria italica*) and rice control seed shattering (Lin et al., 2012; Liu et al., 2022). SH1 acts as a transcription repressor to repress the expression of some genes in the abscission zone in foxtail millet (Liu et al., 2022). We also found that *zmyab15* had shorter height compared to LH244 at adult stage (Figure S12E). YABs contain a conserved zinc-finger domain (C2C2) and a YABBY domain that can bind to DNA (Zhang et al., 2020). In this study, we found that ZmYAB15 can bind to the cis-elements with C rich (Figure S13D, S13E), which is different to AtYAB1/FIL in *Arabidopsis* that could bind to AT rich cis-elements (Boter et al., 2015). Meanwhile, the upstream components of YAB factors are not well known by now. We found that ZmCPK2 could directly phosphorylate ZmYAB15, which could enhance the DNA transactivating activity of ZmYAB15 on the gene *ZmSTRK1* under normal conditions for maintaining and/or promoting plant growth and development. We also detected that ZmCPK17 could weakly interact with and strongly phosphorylate ZmYAB15, and the phosphorylation could weaken the DNA binding activity of ZmYAB15 when ZmCPK17 was activated (Figure 7D, S13I). Thus, we favor a model in which drought stress or ABA triggers Ca^2+^ oscillations that might correspondingly activate ZmCPK17, which then phosphorylates and inhibits ZmCPK2, resulting in inactivating ZmYAB15 due to reduced phosphorylation by ZmCPK2, and/or increased phosphorylation by ZmCPK17, which would turn off its roles on plant growth and development, but activate drought stress response (Figure 7D). Considering that *ZmCPK2-OE*, *ZmCPK17-OE* and *zmyab15* do not affect the hundred grain weight or plot yield (Figure S2I, S2J, S5H, S12F), and the height difference between *ZmCPK17-OE* and LH244 is not significant, while the height of *ZmCPK2-OE* and *zmyab15* may be shorter (Figure S2H, S5G, S12E), thus, they may be the good candidates to be used for improving drought resistance in maize breeding.

## METHODS

### Plant materials and growth conditions

All transgenic maize (*Zea mays* L.) plants and CRISPR/Cas9 mutants were obtained from the Center for Crop Functional Genomics and Molecular Breeding (Center), China Agricultural University (CAU), Beijing. For overexpressing lines, the coding sequences were cloned into pBCXUN vector based on the previous report (Chen et al., 2009). In the preliminary stage, the Center collected a gene list that included more than 700 genes, which might play important roles in various physiological processes from different research groups in CAU. *ZmCPK2* and *ZmCPK17* were included in the gene list. For CRISPR/Cas9 lines, the guide RNA targets were designed using CRISPR-P (http://crispr.hzau.edu.cn/CRISPR2/) and inserted into pBUE411 (Xing et al., 2014). All the constructs were transformed into inbred line LH244 by *Agrobacterium*-mediated transformation. The homozygous lines obtained by self-pollinated T3 were used in this study. The CRISPR/Cas9 mutants were examined using the primers listed in Table S1. The T-DNAs in CRISPR/Cas9 mutants were removed during the crossing.

For seedling stage analysis, maize seeds were planted in pots containing nutrient soil, vermiculite and Pindstrup soil mix (Denmark) (1:1:1), and the seedlings were grown in a glasshouse at 25 °C under a 14 h light/10 h dark photoperiod with c. 200 µmol m^-2^ s^-1^ photo density and 50% - 60% relative humidity. The pots with four holes at the bottom were placed on a tray and water was added to the tray to make sure that every pot absorbs the same amount of water. For adult stage analysis, maize seedlings were grown in the field with a waterproof cloth cover that can be moved to prevent water from raining if needed.

### Physiological phenotype analyses

For drought phenotype analyses of maize at seedling stage, four plump seeds were planted in one pot with soil mixture. Each transgenic line or CRISPR/Cas9 line and the control LH244 had 3 repeat pots, respectively in one tray. The seedlings were fully watered at 6-d-old and then stopped water supply until the transgenic line or CRISPR/Cas9 line had different leaf wilting phenotype comparing with LH244. Pictures were taken to record the phenotypes. The leaf fresh weight, saturated fresh weight (leaf fully soaked in water) or dry weight was obtained for calculating the relative water content (RWC, RWC = (fresh weight - dry weight)/(saturated fresh weight - dry weight)), or rewatered after 3 - 5 days to record the survival rates.

For growth and development phenotype analyses of maize seedlings, each transgenic line, CRISPR/Cas9 line or the control LH244 with 3 pots in one tray were well-watered until 12-d-old. Photos of seedlings were taken in pot, and then washed away the soil from the roots to weigh the whole plants in one pot.

For drought phenotype analyses of maize seedlings at adult stage, the transgenic lines, CRISPR/Cas9 mutants or the control LH244 seedings were sowed side by side with 10 individual seedlings per row, 25 cm between each seedling. The field was divided to two parts, one well-watered and the other stopped water supply at 20-d-old until the transgenic line or CRISPR/Cas9 line had different leaf wilting phenotype comparing with LH244. Pictures of well-watered plants and drought treatment plants were taken.

For water loss rate, the seedlings were grown until the second leaf totally expanded under well-watered condition. The second leaves were cut and weighted at different times. Four leaves were weighted in each replicate and three biological replicates were performed for one experiment.

For the infrared thermal imaging assay, 8-d-old seedlings grown in the greenhouse had water withheld for 2 days before being imaged with a VarioCAM HD camera (JENOPTIK). Leaf temperature was measured using IRBIS 3 professional software.

For stomatal aperture assay, the medium 1-cm piece of the first fully expanded leaf from 8-d-old maize seedlings was excised and immersed in stomatal opening solution buffer (10 mM MES-KOH (pH 5.7), 10 mM KCl, 50 µM CaCl_2_) in 5 mL transparent centrifuge tube. Flat the tubes with the abaxial surface of leaf blade towards the light for 4 h to open the stomata. Then, transfer the leaf to fresh opening solution buffer with 12 µM ABA for another 1-2 h. The assay was performed on the shelf of green house at 25°C and the vertical distance between leaf and light was 15 cm. Each line need 3 leaves for opening (Mock) and 3 leaves for ABA treatment after opening (ABA). All leaves were coated with colorless nail polish. After nail polish was dried out, the abaxial epidermis was peeled with a transparent scotch tape and photographed using an Olympus BX53 microscope (Olympus) with a 40-fold objective lens. Stomatal apertures were measured with the Image J software. About 300 stomatal apertures from each treatment were measure in one experiment. Error bars represent ± SD. Three biological replicates were performed and obtained the similar results.

### Plasmid construction

The full-length or truncated CDS of ZmCPK2, ZmCPK17 or ZmYAB15 was amplified and constructed into different vectors by double enzyme digestion, respectively. The pGADT7 and pGBKT7 were used for yeast two-hybrid assay (Zhu et al., 2020). pGEX4T-1, pET28a and pMAL-c5x were used for prokaryotic expression protein. *Super1300*-GFP/MYC/Flag/mCherry were used for transient expression in maize protoplasts or in *N. benthamiana* leaves. pSPYNE173 and pSPYCE (M) were used for BiFC assay. The mutagenic vectors were cloned form the GST-tagged constructs using the primers designed in The QuickChange Primer Design (https://www.agilent.com.cn/store/primerDesignProgram.jsp). All the primers were listed in Table S2.

### ABA treatment and protein purification

For *ZmCPK2-HF* seedlings, the 8-d-old plants under well-watered condition were removed from soil, and the soil on roots were washed away with water slowly. The roots were soaked in water containing 50 µM ABA for 0-1.5 h, and the second leaves (about 100 mg) were harvested and used for total protein isolation. Maize protoplasts were isolated and transfected according to the method described previous (Ma et al., 2012). The protoplasts transiently expressing different vectors (incubated overnight under darkness) were treated with 50 µM ABA for 0-1 h. The total proteins of seedlings or protoplasts were extracted with IP buffer (10 mM HEPES (pH 7.5), 100 mM NaCl, 1 mM EDTA, 10% Glyceral (v/v) and 1× protease inhibitor cocktail, 10 mM PMSF) and immunoprecipitated by anti-Flag agarose (A2220, Sigma-Aldrich) for 3 h at 4°C. The immunoprecititated proteins on agarose were washed three times with PBS buffer and used for *in vitro* phosphorylation assay. The signal bands was measured by Image J software and relative kinase activity was calculated by GraghPad Prism 8. Error bars represent ± SD.

### *In vitro* phosphorylation assay

To detect the phosphorylation of protein kinase on candidate substrates, *in vitro* phosphorylation assays were performed as described previously (Deng et al., 2022). For recombinant proteins, the constructs with GST, His or MBP tag were transformed into the *Escherichia coli* strain BL21 (DE3) and cultured at 18-22°C for 14-16 h. The total proteins were isolated and then purified using glutathione Sepharose 4B (GE Healthcare), Ni Sepharose (GE Healthcare) or amylose resin (NEB). The *in vitro* phosphorylation assay was performed using proteins purified from maize seedlings or protoplasts or 2 µg recombinant proteins as kinases and 5-10 µg universal substrate MyBP or recombinant proteins as substrates. The mixed proteins were incubated in kinase buffer (25 mM Tris-HCl (pH 7.5), 10 mM MgCl_2_ 50 µM ATP and 1 µCi of [γ-^32^P]ATP) in a 20 µl volume at 30 °C for 30 min. The reactions were stopped by adding 4 µl of 6× SDS loading buffer and the proteins were separated on 8%-15% SDS-PAGE. The phosphorylated signals were visualized by autoradiography (Typhon 9410 imager).

For Ca^2+^-dependent kinase assay, we obtained defined free Ca^2+^ concentrations with adding 5 mM EGTA for 0 nM; 5 mM EGTA and 1.306 mM CaCl_2_ for 20 nM, and 3 mM CaCl_2_ for 100 nM, and 3.55 mM CaCl_2_ for 200 nM, and 4.28 mM CaCl_2_ for 500 nM, and 4.61 mM CaCl_2_ for 1000 nM, and 4.865 mM CaCl_2_ for 3000 nM, and 4.921 mM CaCl_2_ for 5000 nM, and 4.967 mM CaCl_2_ for 10000 nM (calculated with https://somapp.ucdmc.ucdavis.edu/pharmacology/bers/maxchelator/CaMgATPEGTA-NIST.htm) (Brandt et al., 2012; Geiger et al., 2010).

### MST assay

Microscale thermophoresis (MST) assay was performed as previously described (Ding et al., 2022). The recombinant proteins GST-ZmCPK17, GST-ZmCPK2, GST-ZmCPK2-NKJ or GST-AtCPK3 was purified using glutathione Sepharose 4B, respectively, and the buffers were replaced by PBS buffer with 0.005% Tween-20 (PBST) using column A (NanoTemper Technologies). Next, 10 µM proteins (100 µL) were labeled with excess NHS NT-647 dye at a molar ratio of 1:5 for 30 min at room temperature in the dark. Free unlabeled dye was removed by column B, which was re-equilibrated with PBST buffer. Different concentrations CaCl_2_ were added to MST buffer (50 mM Tris-HCl (pH 7.4), 10 mM MgCl_2_, 150 mM NaCl, 5 mM EGTA and 0.05% Tween-20). To obtain defined free Ca^2+^ concentrations we added 4.9938 mM for 20 µM, 4.9585 mM for 10 µM, 4.904 mM for 5 µM, 4.805 mM for 2.5 µM, 4.63 mM for 1.25 µM, 4.2874 mM for 625 nM, 3.7523 mM for 312.5 nM, 3.003 mM for 156.25 nM, 2.1458 mM for 78.125 nM, 1.515 mM for 39.0625 nM, 0.913 mM for 19.5313 nM, 0.509 mM for 9.7656 nM CaCl_2_. MST buffer with different CaCl_2_ was mixed with the same amount of labeled protein (10 µL). After gently mixing proteins and CaCl_2_, the mixtures were loaded into capillaries (NanoTemper Technologies) and analyzed by NanoTemper Monilith Instrument (NT.115) (NanoTemper Technologies) with 20% LED power and 20% MST power. Dissociation constant (Kd) was calculated using MO. Affinity Analysis (v2.2.4).

### RNA-seq assays and Quantitative RT-PCR

For RNA-seq assay, total RNAs were extracted from the second fully expanded leaves of three 9-d-old seedlings treated with or without 50 µM ABA grown under well-watered conditions with HiPure Plant RNA Mini Kit (R4151; Magen). Two independent replicates were performed. RNA integrity was evaluated using a 2100 Bioanalyzer (Agilent Technologies). The RNA libraries were constructed and sequenced on an illumina nova 6000 at Novogenge (Beijing) and 150 bp paired-end reads were generated. The methods used for RNA-seq data analysis were described previously (Li et al., 2022b).Raw data (raw reads) in fastq format were subjected to quality control by Fastp (v.0.19.7) and the clean reads were mapped to the reference maize genome (B73 RefGen_v4, AGPv4) using HISAT2 (V.2.2.0) with default parameters (Trapnell et al., 2013). FPKM of each gene was calculated using Cufflinks (v.2.2.1). The differential expression analyses were performed using Cuffdiff (v.2.2.1) with default parameters. Significant DEGs were identified as those with FDR <0.05. The heatmap was drawn based on the expression levels of indicated genes and the genes are listed in Supplemental Data Set sheet 2-3. GO enrichment was performed with the accession numbers of overlapped significant DEGs via agriGO v2.0 (Tian et al., 2017). The raw data have been deposited in the NCBI Sequence. Read Archive is accessible through SRA accession code RPJNA944568 (https://www.ncbi.nlm.nih.gov/sra/PRJNA944568).

For quantitative RT-PCR, the integrity of total RNA was evaluated using agarose gel electrophoresis followed by reverse transcription with Maxima H Minus First-Strand cDNA Synthesis Kit (K1682; Thermo Scientific). Quantitative RT-PCR was performed with SYBR Green regent (Takara) on StepOne Plus Real-Time PCR system (Applied Biosystem). The maize polyubiqutin gene *Ubi* was used as an internal control. Primer sequences for quantitative RT-PCR were listed in Table S3. The experiments were independently repeated three times.

## ACCESSION NUMBERS

Sequence data from this article can be found in the Gramene data libraries under the following accession numbers:

ZmCPK2, GRMZM2G040743/Zm00001d029758/Zm00001eb022180; ZmCPK17, GRMZM2G463464/Zm00001d041871/Zm00001eb139060; ZmYAB15, GRMZM2G529859/Zm00001d017391/Zm00001eb248640; RD22, GRMZM5G800586/Zm00001d043538/Zm00001eb154060; ZmSTRK1, GRMZM2G166719/Zm00001d013858/Zm00001eb220340; ZmCPK6, GRMZM2G347047/Zm00001d051835; ZmCPK7, GRMZM2G032852/Zm00001d027480/Zm00001eb002180; ZmCPK9, GRMZM2G121228/Zm00001d034376/Zm00001eb060140; ZmCPK10, GRMZM2G353957/Zm00001d040996/Zm00001eb132720; ZmCPK11, GRMZM2G028926/Zm00001d034372/Zm00001eb060110; ZmCPK14, GRMZM2G035843/Zm00001d053016/Zm00001eb200920; ZmCPK15, GRMZM2G047486/Zm00001d004812/Zm00001eb091710; ZmCPK26, GRMZM2G154489/Zm00001d021623/Zm00001eb322620; HAK11, GRMZM2G005040/Zm00001d002333/Zm00001eb070910.

## ACKNOWLEDGMENTS

This research is supported by grants from the National Science Foundation of China (32030008 and 31921001) and the Beijing Outstanding University Discipline.

## AUTHOR CONTRIBUTIONS

Z.G. and X.H. conceived the study and designed the research. X.H. performed most of the experiments and analyzed the data. J.C. assisted for analyzing the RNA-seq data and Z.L. for LC-MS/MS analysis. Z.G. and X.H. wrote the manuscript. All other authors discussed the results and commented on the manuscript.

## Supplemental Figures

**Figure S1.**
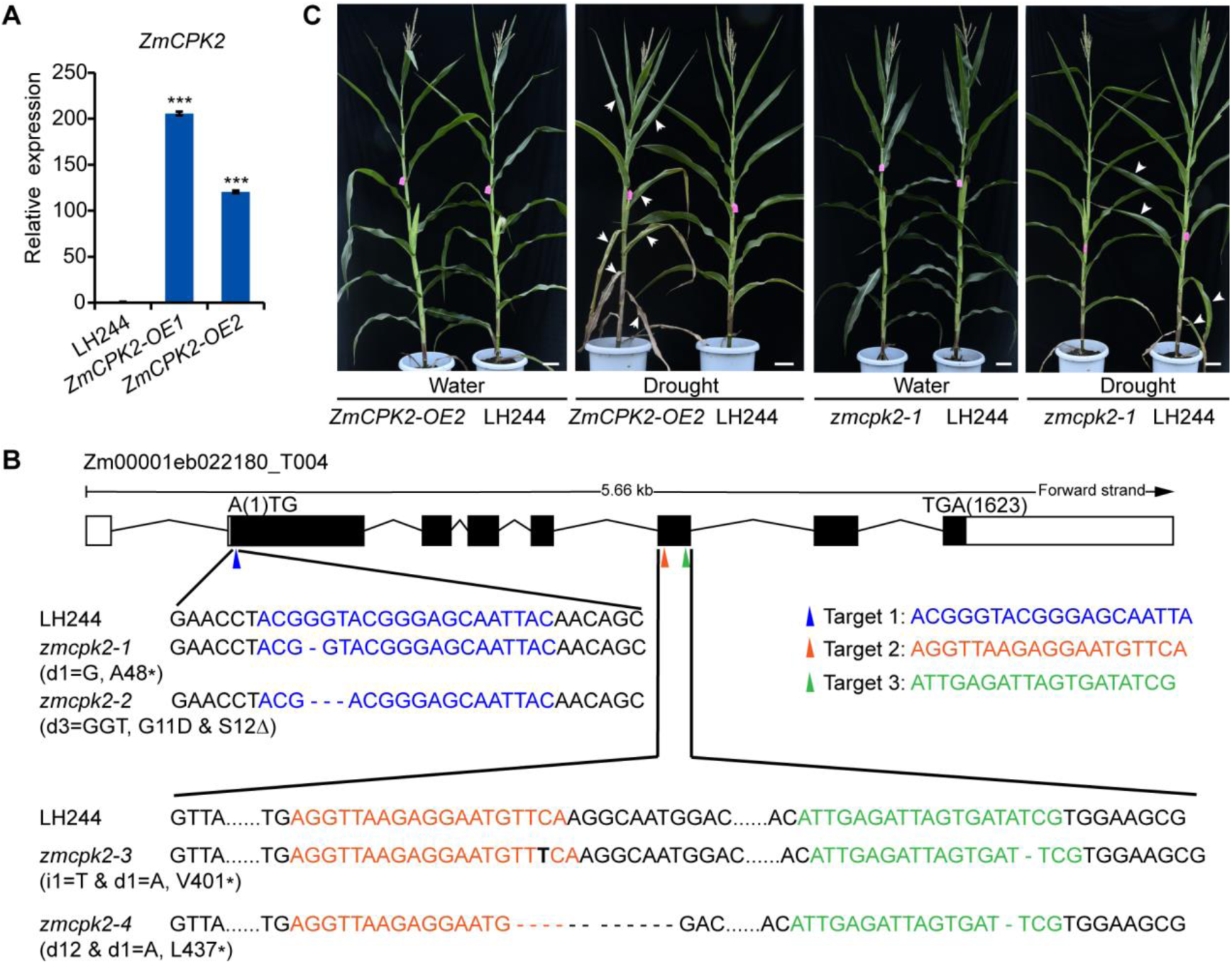
CRISPR/Cas9 mutation analysis of *ZmCPK2*. **(A)** Relative expression levels of *ZmCPK2* in transgenic plants *ZmCPK2-OE1/2* and LH244. Total RNAs were extracted from 9-d-old well-watered seedlings and used for qPCR. Error bars represent ± SD of three technical replicates from one experiment and three experiments were performed with similar results. * represents significant difference with LH244, *** p<0.001, Student’s *t*-test. **(B)** Detail information for the *zmcpk2-1/-2/-3/-4* mutations. The gene structure is shown in the top from Gramene database. Black boxes represent exon and black lines represent intron. The blue triangle indicates the location of the single guide RNA target in the first exon that creates the mutant *zmcpk2-1*, with a single nucleotide G deletion causing premature termination at Ala48, and the mutant *zmcpk2-2*, with three base deletion causing an amino acid Gly **(G)** changed to Asp (D) and an amino acid Ser (S) deletion. The orange and green triangles indicate the location of the double guide RNA targets in the fifth exon that created the mutant *zmcpk2-3*, with a single base T insertion and a single A deletion causing premature termination at Val401, and the mutant *zmcpk2-4*, with a 12 bp deletion and a single A deletion causing premature termination at Leu437. The nucleotide sequences surrounding guide RNA targets in mutants and LH244 are shown. **(C)** Transgenic *ZmCPK2-OE2* plants are more sensitive and *zmcpk2-1* plants are more tolerant to drought treatment compared with LH244 at the adult stage. The plants were grown in the field with a rain shelter. The control plants were well-watered (Water), while the drought-treated plants were stopped water supply at 20-d-old (Drought). 10 seedlings in one row showed the similar leaf senescence phenotypes. One representative seedling was replanted in a pot for taking pictures. Bar = 10 cm.

**Figure S2.**
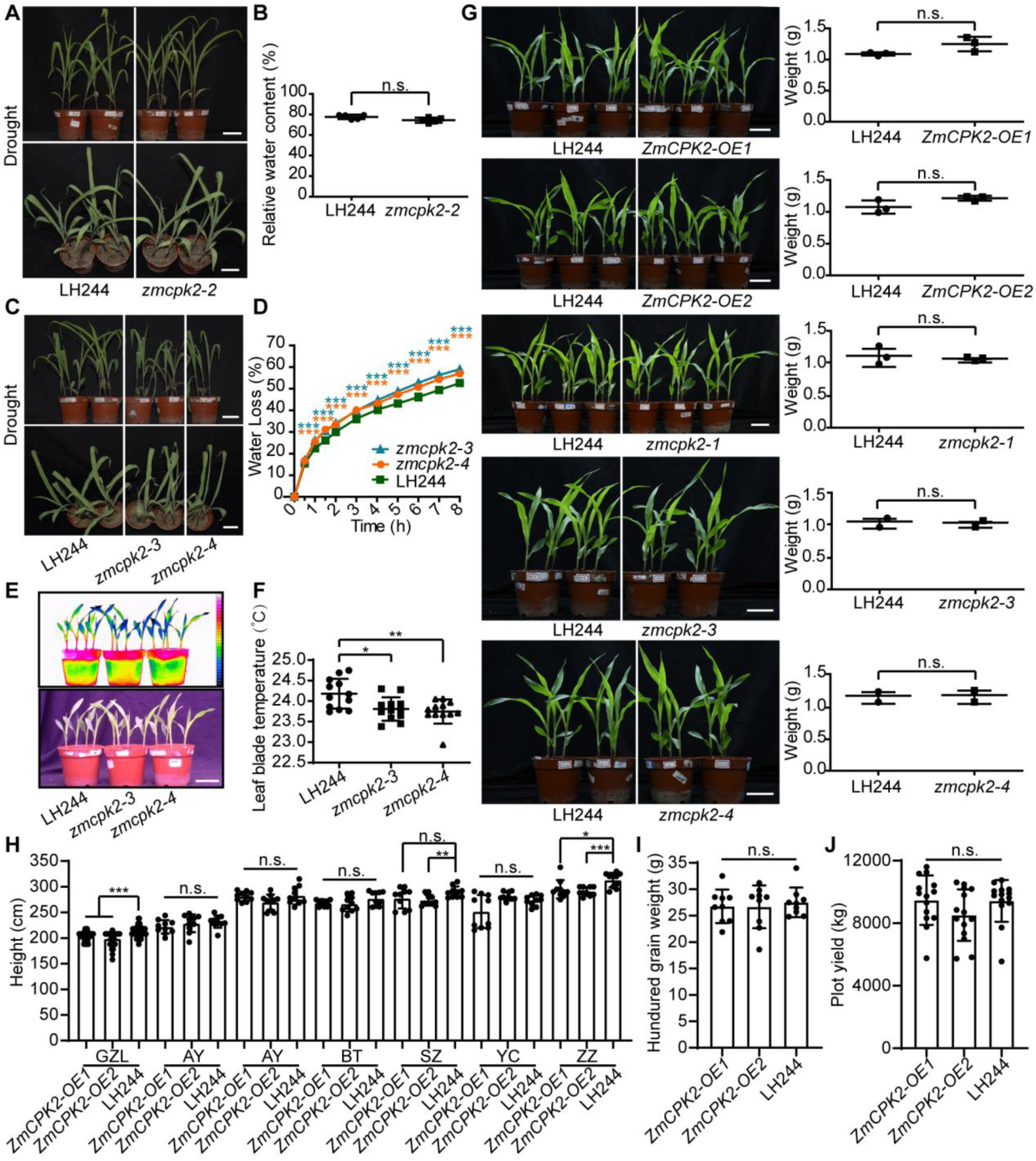
The growth and development phenotypes and drought phenotypes of *ZmCPK2-OE* line and *zmcpk2* mutants. **(A)** The CRISPR/Cas9 mutant *zmcpk2-2* has no significant drought phenotype compared with LH244. The pictures were taken for 14-d-old seedlings treated with drought stress. Bar = 5 cm. **(B)** Relative water content of leaves in **(A)**. The fresh weight, saturated fresh weight and dry weight of all cut leaves were measured for calculating the relative relative water. Each point represents a group of leaves from one seedling. Error bars represent ± SD from one experiment and three experiments were performed with similar results. n.s. represents no significant difference with LH244, Student’s *t*-test. **(C)** The CRISPR/Cas9 mutants *zmcpk2-3/-4* are more sensitive to drought treatment compared with LH244. The pictures were taken for 14-d-old seedlings treated with drought stress. Bar = 5 cm. **(D)** Water loss of *zmcpk2-3/-4* and LH244. Error bars represent ± SD of three technical replicates from 12 second expanded leaves in one representative experiment. Asterisks indicate a significant difference compared with the wild type: *** p < 0.001, Student’s *t*-test. **(E)** Comparison of the leaf temperatures of *zmcpk2-3/-4* and the wild type as determined by an infrared thermal imaging assay. 8-d-old seedlings grown in soil with well watering were used for temperature measurement with a far-red CCD camera. **(F)** Statistical analysis of leaf temperature in **(E)**. Data are means ± SD of three replicates (eight leaves from one pot were measured per replicate) in one experiment. Four experiments were done with similar results. Asterisks indicate a significant difference compared with the wild type: * p < 0.05, ** p < 0.01, Student’s *t*-test. **(G)** The growth and development phenotype analyses of different seedlings. 12-d-old seedlings of *ZmCPK2-OE1/2*, *zmcpk2-1/-2/-3/-4* or LH244 were grown in one pot under well-watered condition, respectively. The weights of total plants in one pot were recorded. Each point represented a weight data from one pot. Error bars represent ± SD from one experiment. n.s. represents no significant difference with LH244, Student’s *t*-test. **(H)** The height of *ZmCPK2-OE* was shorter than LH244 in some field sites. The capital letters in the abscissa represent different regions in China that performed field trials. GZL: Gongzhuling, China, 44° N, 124° E; AY: Anyang, China, 36° N, 114° E; BT: Baotou, China, 41° N, 110° E; SZ: Shangzhuang, Beijing, China, 40° N, 116° E; YC: Yinchuan, China, 38° N, 106° E; ZZ: Zhuozhou, China, 39° N, 116° E. The statistical analysis was done with Graphpad Prism 8. Error bars represent ± SD. n.s. represents t no significant difference with LH244. * represents significant difference with LH244, * p<0.05, ** p<0.01, *** p<0.001, Student’s *t*-test. **(I,J)** The hundred grain weight and plot yield of *ZmCPK2-OE* in the field. Error bars represent ± SD. n.s. represents no significant difference with LH244.

**Figure S3.**
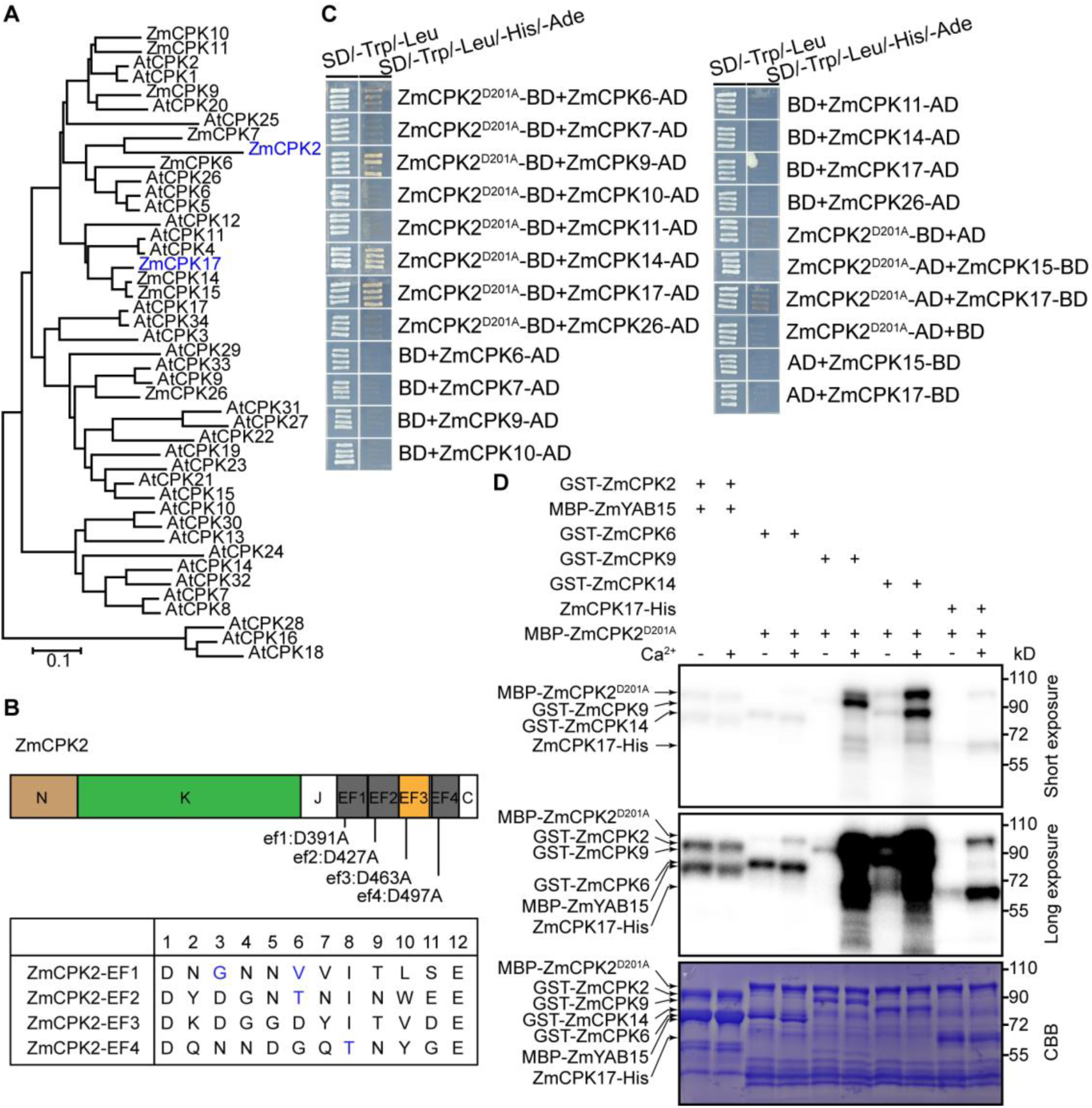
ZmCPK2 interacts with some ZmCPKs. **(A)** Phylogenetic relationships of ZmCPKs in Zea mays and AtCPKs in Arabidopsis. Phylogenetic tree was constructed based on the full-length protein sequences by MEGA 5.0. **(B)** The protein structure of ZmCPK2. The annotations were obtained from Gramene database. EF3 in pale brown is the only intact EF hand in ZmCPK2. N: N terminus, K: Kinase domain, J: Junction domain, EF: EF hands, C: C terminus, ef1/2/3/4 with the first Asp (D) in EF hand motif mutated to Ala (A) was shown. Four EF hand motifs in ZmCPK2 were shown, and the blue characters represent the unconservative amino acid. **(C)** The kinase dead form ZmCPK2^D201A^ interacts with some ZmCPKs in yeast two-hybrid assay. AD: Gal4 activation domain, BD: Gal4 DNA-binding domain. **(D)** ZmCPK6/9/14/17 phosphorylates the kinase dead MBP-ZmCPK2^D201A^ in Ca^2+^-dependent manner *in vitro*. Recombinant GST-ZmCPK6/9/14 and ZmCPK17-His were subjected to the phosphorylation assay with or without 1 µM free Ca^2+^. Recombinant GST-ZmCPK2 or GST-ZmYAB15 was used as the substrate for Ca^2+^-independent control. Long and short exposure of signals and CBB were shown.

**Figure S4.**
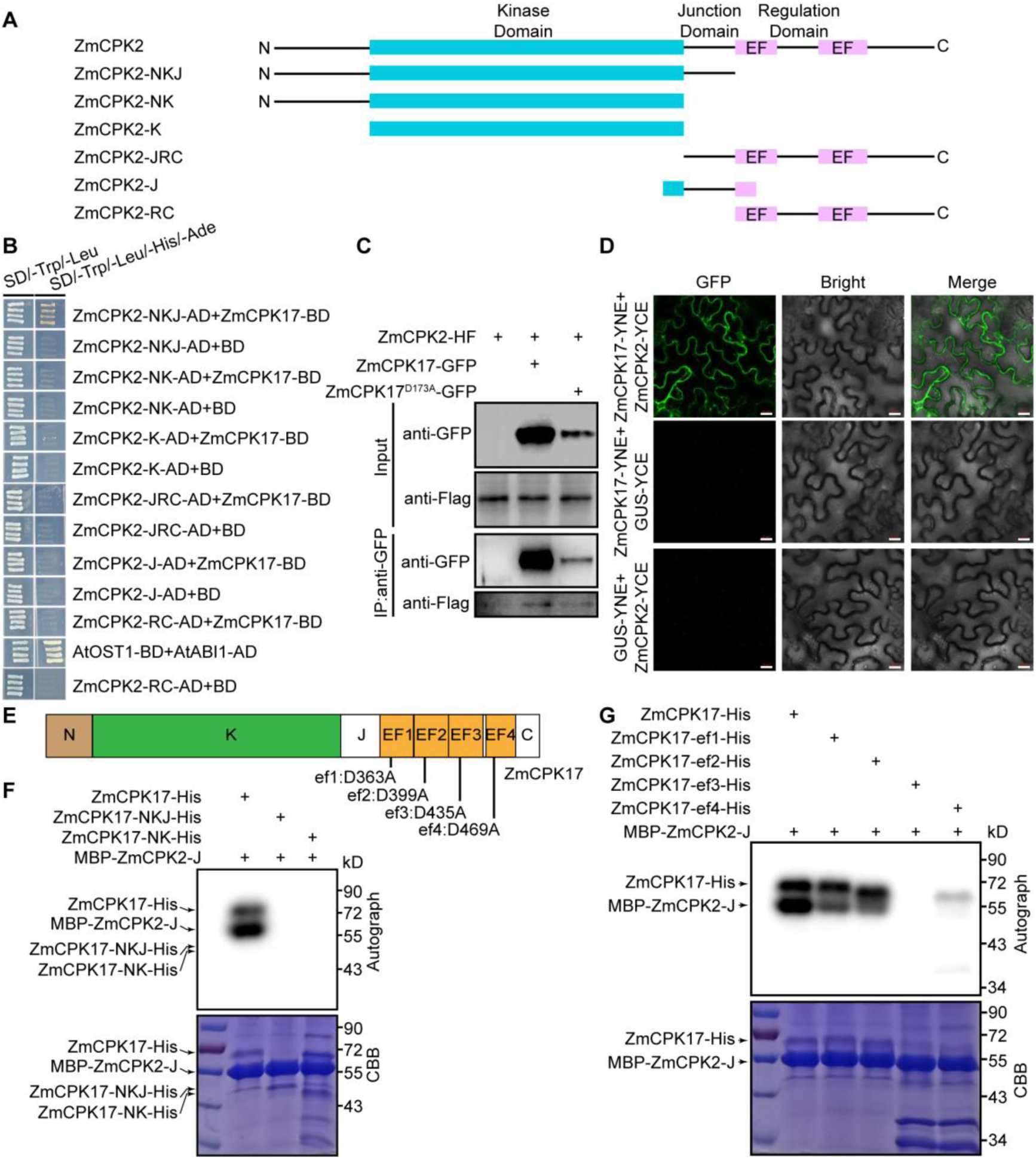
ZmCPK2 interacts with ZmCPK17. **(A)** The structure of ZmCPK2 protein and the diagrams of truncated proteins are shown. **(B)** The truncated form ZmCPK2-NKJ interacts with ZmCPK17 in yeast two-hybrid assay. The combination of AtABI1-AD and AtOST1-BD was used as a positive control. **(C)** ZmCPK2 interacts with ZmCPK17 by Co-IP assay in maize protoplasts. ZmCPK17-GFP and ZmCPK17^D173A^-GFP were transiently expressed in *ZmCPK2-HF2* protoplasts. Total proteins were isolated, and ZmCPK17-GFP or ZmCPK17^D173A^-GFP was immunoprecipated (IPed) with anti-GFP agarose, respectively. The IPed proteins were detected with anti-Flag or anti-GFP antibodies, respectively. **(D)** ZmCPK2 interacts with ZmCPK17 in *N. benthamiana* leaf cells shown by the BiFC assay. The YFP signal was visualized by a confocal microscopy. Bar =20 µm. GUS-YNE and GUS-YCE were used as negative controls. **(E)** The protein structure of ZmCPK17. The annotations were obtained from Gramene database. N: N terminus, K: Kinase domain, J: Junction domain, EF: EF hands, C: C terminus, ef1/2/3/4 with an Asp (D) mutation was shown. **(F)** Comparing the kinase activity of different truncated forms of ZmCPK17 *in vitro*. Recombinant ZmCPK17-His, ZmCPK17-His-NKJ or ZmCPK17-His-NK was subjected to the phosphorylation assay with MBP-ZmCPK2-J as the substrate, respectively. **(G)** Comparing the kinase activity of ZmCPK17 with different mutations in four EF hands. Recombinant ZmCPK17-His or ZmCPK17-ef1/2/3/4-His was subjected to the phosphorylation assay *in vitro* with MBP-ZmCPK2-J as the substrate, respectively.

**Figure S5.**
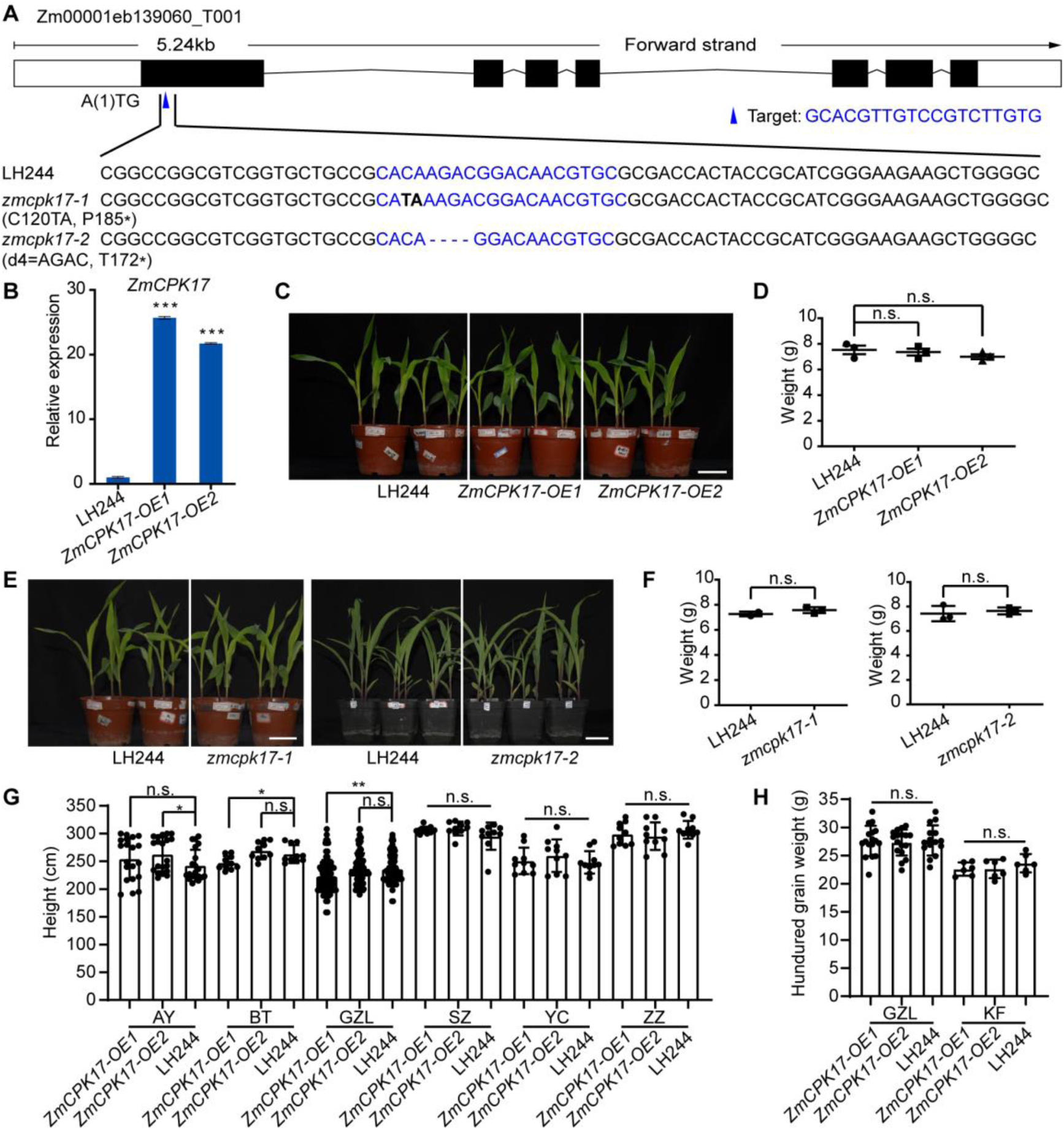
The growth and development phenotype of *ZmCPK17-OE* and *zmcpk17*. **(A)** Detail information for the *zmcpk17-1/2* mutations. The gene structure was shown in the top from Gramene database. Black boxes represent exon and black lines represent intron. The blue triangle indicates the location of the single guide RNA target in the first exon that creates the mutant *zmcpk17-1*, with C120 changing to TA, producing a frame shift and an early putative translation stop codon after Pro180. *ZmCPK17* with four bases deletion (AGAC) produces an early putative translation stop codon after Thr172 in mutant *zmcpk17-2*. **(B)** Relative expression level analyses of *ZmCPK17* in transgenic plants *ZmCPK17-OE1/2* and LH244. 8-d-old seedlings were well-watered and the total RNAs were extracted. Error bars represent ± SD of three technical replicates from one experiment. * represents significant difference with LH244, *** p<0.001, Student’s *t*-test. **(C, E)** The growth and development phenotype analyses of different seedlings. 12-d-old seedlings of *ZmCPK17-OE1/2* **(C)** and *zmcpk17-1/-2* **(E)** were separately grown with LH244 in one tray under well-watered condition. **(D, F)** The weights of total plants in **(C, E)**. Each point represented a weight data from one pot. Error bars represent ± SD from one experiment. n.s. represents no significant difference with LH244, Student’s *t*-test. **(G)** The height of *ZmCPK17-OE* compared with LH244 in the field. Error bars represent ± SD. n.s. represents no significant difference with LH244. * represents significant difference with LH244, * p<0.05, ** p<0.01, Student’s *t*-test. **(H)** The hundred grain weight of *ZmCPK17-OE* compared with LH244 in the field. KF: Kaifeng, China, 35° N, 114° E. Error bars represent ± SD. n.s. represents no significant difference with LH244. Student’s *t*-test.

**Figure S6.**
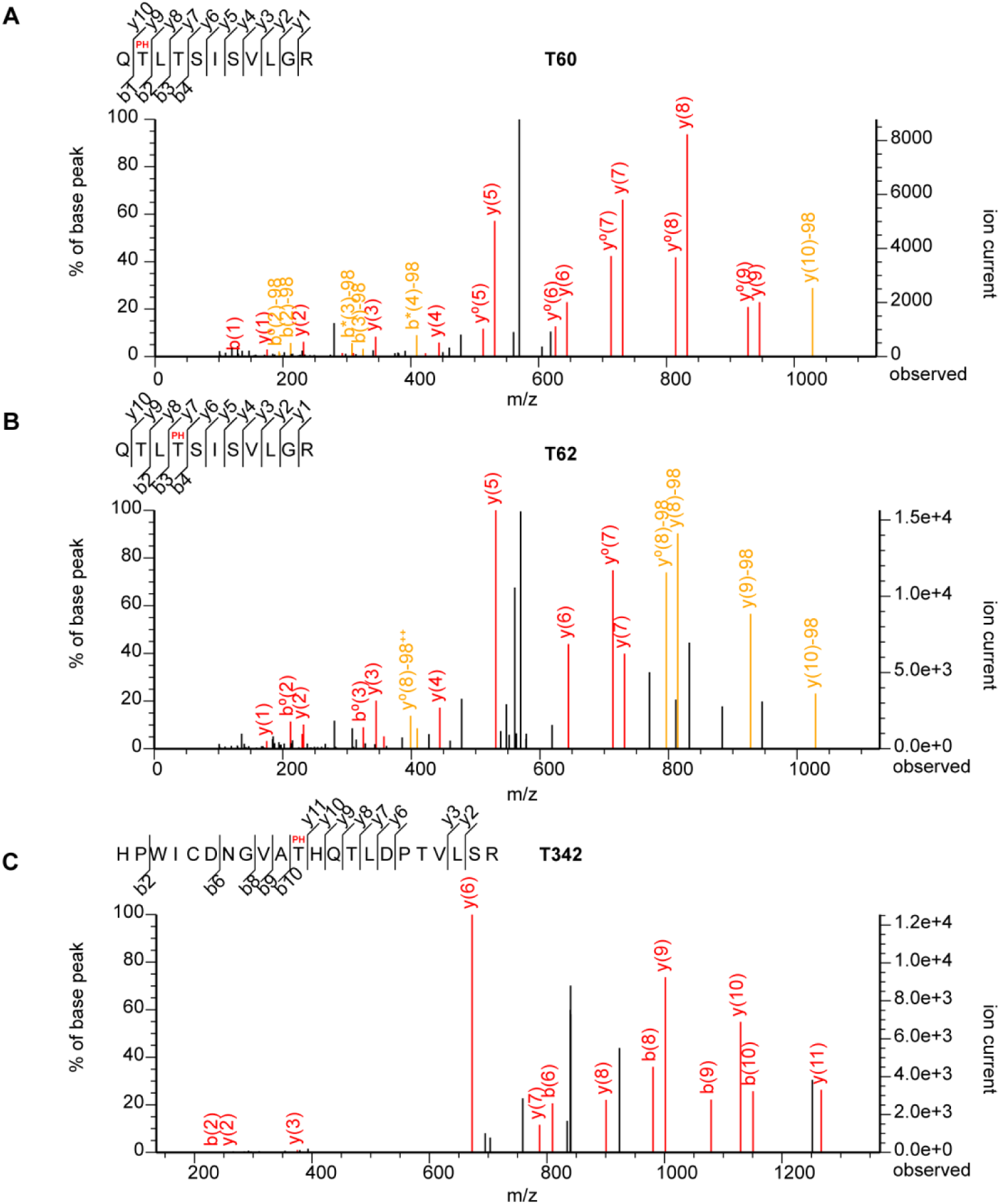
The candidate phosphorylation sites identified by LC-MS/MS. **(A-C)** Three candidate sites identified with phospho (PH) modification: T60 **(A)**, T62 **(B)** and T342 **(C)**.

**Figure S7.**
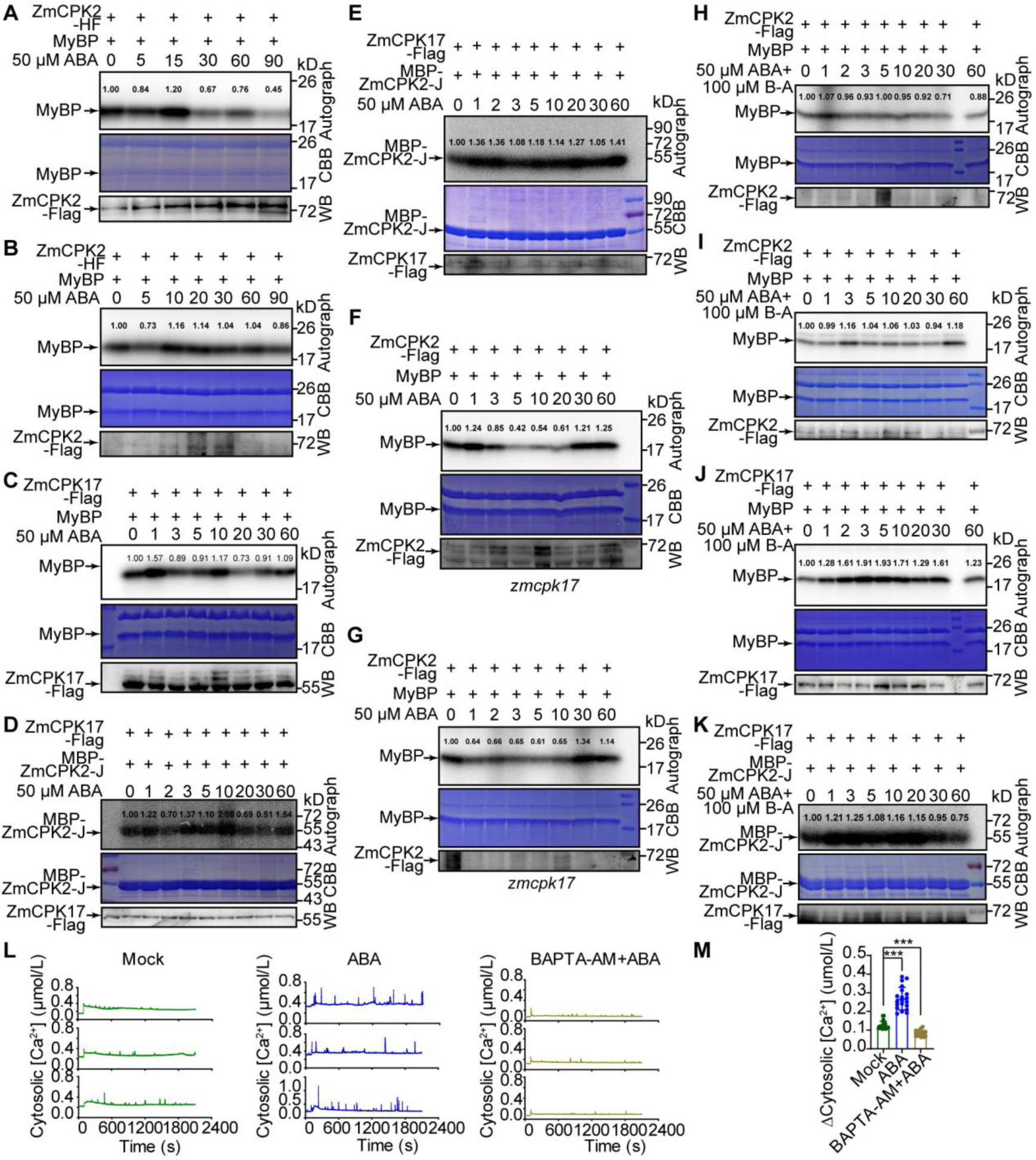
The activities of ZmCPK2 and ZmCPK17 are dynamically modulated by ABA. **(A, B)** The other two repeats of the kinase activity of ZmCPK2 regulated by ABA as shown in Figure 2E. The total proteins were extracted from 8-d-old seedlings of *ZmCPK2-HF2* treated with 50 µM ABA for different times (0-90 min). ZmCPK2-HA-Flag (ZmCPK2-HF) was immunoprecipated (IPed) from total proteins with anti-Flag agarose. IPed proteins were subjected to *in vitro* phosphorylation assay using MyBP as a substrate. The autograph, CBB and the protein levels of ZmCPK2 in each sample (WB) were shown. The numbers represent the quantification of the phosphorylation signals of MyBP by the Image J software. **(C)** The kinase activity of ZmCPK17 regulated by ABA similar to the data shown in Figure 5B. *Super: ZmCPK17-Flag* was transformed to maize protoplasts and treated with 50 μM ABA for different times (0-60 min). IPed proteins were subjected to *in vitro* phosphorylation assay using MyBP as the substrate. The numbers represent the quantification of the phosphorylation signals of MyBP by the Image J software. **(D, E)** The other two repeats of the kinase activity of ZmCPK17 regulated by ABA as shown in Figure 5B. *Super: ZmCPK17-Flag* was transformed to maize protoplasts and treated with 50 μM ABA for different times (0-60 min). IPed proteins were subjected to *in vitro* phosphorylation assay using recombinant MBP-ZmCPK2-J as the substrate. The numbers represent the quantification of the phosphorylation signals of MBP-ZmCPK2-J by the Image J software. **(F, G)** The other two repeats of ABA-mediated ZmCPK2 activity in *zmcpk17* as shown in Figure 5C. *Super: ZmCPK2-Flag* was transformed to the protoplast isolated from 10-d-old seedlings of *zmcpk17* and treated with 50 μM ABA for different times (0-60 min). IPed proteins were subjected to *in vitro* phosphorylation assay using MyBP as the substrate. The numbers represent the quantification of the phosphorylation signals of MyBP. **(H, I)** The other two repeats of the activity of ZmCPK2 regulated by ABA and BAPTA-AM as shown in Figure 5D. *Super: ZmCPK2-Flag* was transformed to the protoplast isolated from 10-d-old seedlings of LH244 and treated with 50 μM ABA and 100 μM BAPTA-AM (B-A) for different times (0-60 min). IPed proteins were subjected to *in vitro* phosphorylation assay using MyBP as the substrate. The numbers represent the quantification of the phosphorylation signals of MyBP. **(J, K)** The other two repeats of the activity of ZmCPK17 regulated by ABA and BAPTA-AM as shown in Figure 5E. *Super: ZmCPK17-Flag* was transformed to maize protoplasts and treated with 50 μM ABA and 100 μM BAPTA-AM (B-A) for different times (0-60 min). IPed proteins were subjected to *in vitro* phosphorylation assay using MyBP (**J**) or recombinant MBP-ZmCPK2-J (**K**) as the substrate. The numbers represent the quantification of the phosphorylation signals of the substrate. **(L)** Time-course analysis of [Ca^2+^]_cyt_ dynamics in protoplasts with no treatment (Mock) or 50 µM ABA (ABA) or 100 µM BAPTA-AM and ABA (BAPTA-AM+ABA). The luminescence was recorded for 2600 s (100 s for resting luminescence, 2000 s for stimulated luminescence and 500 s for discharged luminescence) at 1 s interval. Three independent repeats were shown for each treatment. **(M)** Quantitation of different changes in cytosolic Ca^2+^ (Δ [Ca^2+^]_cyt_ (µmol/L)) in **(L)**. 20 peaks induced in three treatments (Mock, ABA, BAPTA-AM) against to 20 peaks in the rest were calculated. Data represent means ± SD from three independent experiments. *** p<0.001, Student’s *t*-test.

**Figure S8.**
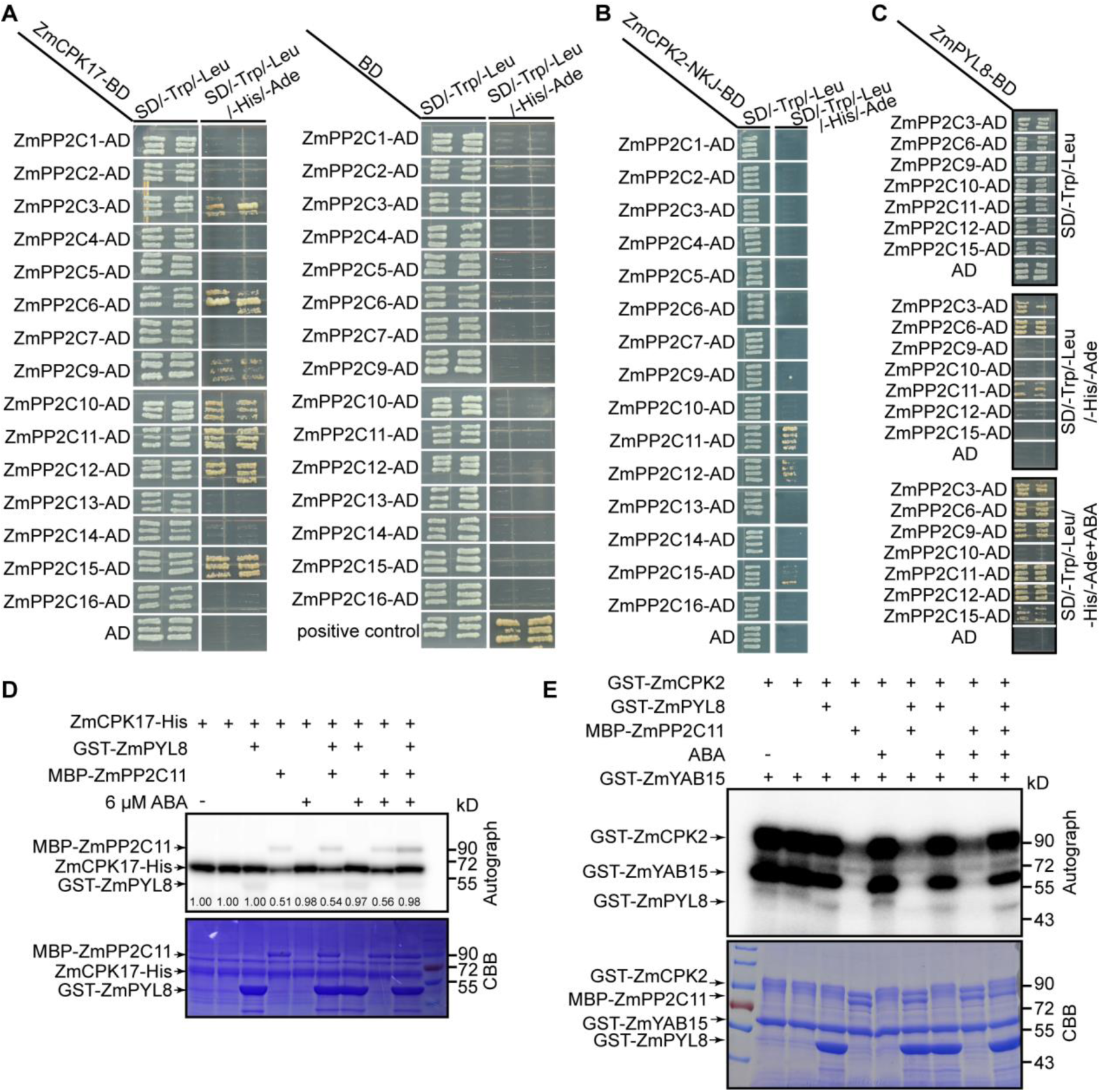
ABA-ZmPYL8 releases the inhibition of ZmPP2C12 on ZmCPK17 and ZmCPK2 *in vitro*. **(A)** ZmCPK17 interacts with ZmPP2C3/6/9/11/12/15 in yeast two-hybrid assay. The combination of AtABI1-AD and AtOST1-BD was used as a positive control. **(B)** ZmCPK2 interacts with ZmPP2C11/12/15 in yeast two-hybrid assay. **(C)** ZmPP2C9/11/12/15 interact with ZmPYL8 in an ABA-dependent manner in yeast two-hybrid assay, while ZmPP2C3/6 interacts with ZmPYL8 in an ABA-independent manner. **(D)** ZmPP2C12 inhibits the activity of ZmCPK17, and ABA activates ZmCPK17 through releasing the inhibition of ZmPP2C12. Recombinant ZmCPK17-His with or without MBP-ZmPP2C12, GST-ZmPYL8 and 6 µM ABA were subjected to the phosphorylation assay. The autograph and CBB were shown. The numbers represent the quantification of the phosphorylation signals of ZmCPK17 by the Image J software. **(E)** ZmPP2C12 inhibits the activity of ZmCPK2, and ABA activates ZmCPK2 through releasing the inhibition of ZmPP2C12. Recombinant GST-ZmCPK2 with or without MBP-ZmPP2C12, GST-ZmPYL8 and 6 µM ABA were subjected to the phosphorylation assay.

**Figure S9.**
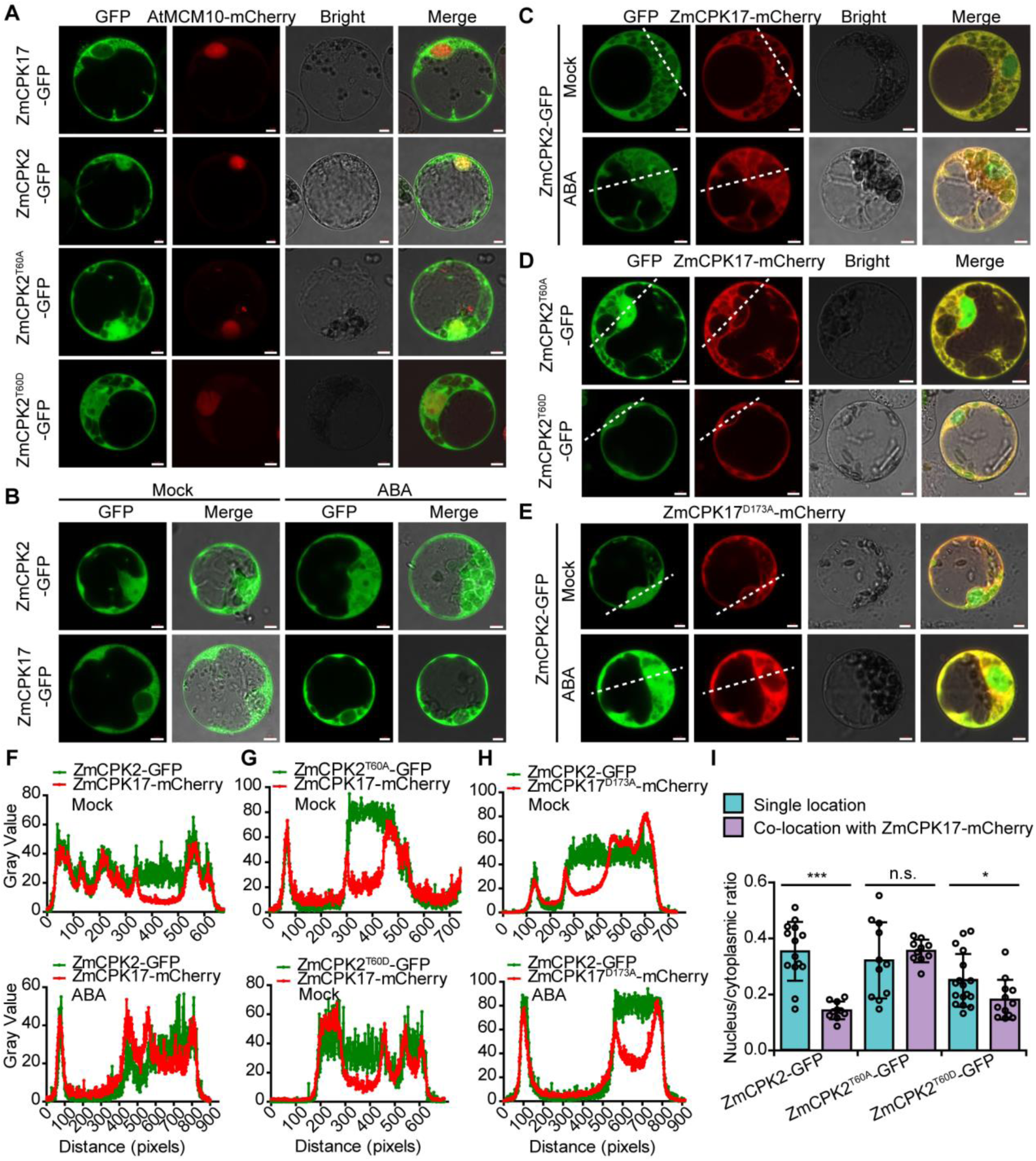
The ZmCPK2 T60 phosphorylation attenuates its nuclear localization. **(A)** Subcellular location of ZmCPK2 with different phosphorylation forms of T60 and ZmCPK17. ZmCPK2-, ZmCPK2^T60A^- or ZmCPK2 ^/T60D^-GFP and ZmCPK17-GFP were cotransformed to protoplasts to detect the subcellular locations, respectively. AtMCM10-mCherry was co-expressed to mark the nucleus. Bar = 5 µm. **(B)** Subcellular location of ZmCPK2 and ZmCPK17 under ABA treatment. ZmCPK2-GFP and ZmCPK17-GFP were transiently expressed in protoplasts that treated with 0 µM (Mock) or 50 µM ABA (ABA). Bar = 5 µm. **(C)** Co-location of ZmCPK2 and ZmCPK17 with or without ABA treatment. ZmCPK2-GFP and ZmCPK17-mCherry were transiently co-expressed in protoplasts that treated with 0 µM (Mock) or 50 µM ABA (ABA). Bar = 5 µm. **(D)** Co-locations of ZmCPK17 and ZmCPK2^T60A^ or ZmCPK2^T60D^. ZmCPK17-mCherry and ZmCPK2^T60A^ –GFP or ZmCPK2^T60D^-GFP were co-expressed, respectively, to detect the co-location. Bar = 5 µm. **(E)** Co-locations of ZmCPK2 and ZmCPK17^D173A^ with or without ABA treatment. ZmCPK2-GFP and ZmCPK17^D173A^-mCherry were co-expressed in protoplasts that treated with 0 µM (Mock) or 50 µM ABA (ABA). Bar = 5 µm. **(F)** The co-location signals of GFP and mCherry in **(C)** were measured by Image J, white dotted lines marked the measured locations. Bar = 5 µm. **(G)** The co-location signals of GFP and mCherry in **(D)** were measured by Image J, white dotted lines marked the measured locations. Bar = 5 µm. **(H)** The co-location signals of GFP and mCherry in **(E)** were measured by Image J, white dotted lines marked the measured locations. Bar = 5 µm. **(I)** The ratios of nucleus to cytoplasmic of GFP signals in **(A, C, D)**. Each circle in the bar chart represents one protoplast cell. The GFP signals in nucleus and cytoplasmic were measured by Image J and the ratios were calculated. * p<0.05, *** p<0.001, n.s. represents no significant difference. Student’s *t*-test.

**Figure S10.**
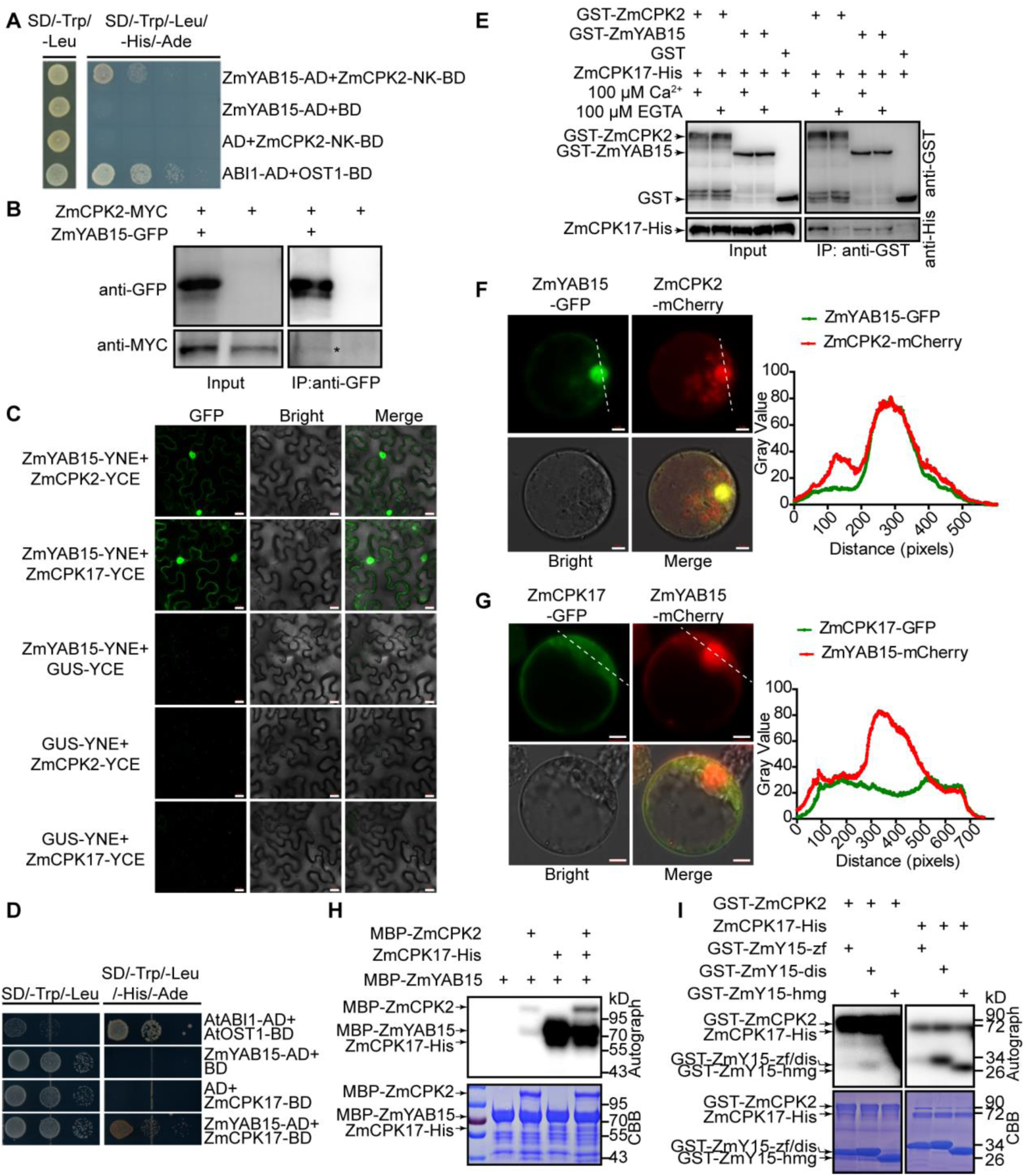
Both ZmCPK2 and ZmCPK17 interact with and phosphorylate ZmYAB15. **(A)** The truncated form ZmCPK2-NK interacts with ZmYAB15 in yeast two-hybrid assay. The combination of AtABI1-AD and AtOST1-BD was used as the positive control. **(B)** The interaction between ZmCPK2 and ZmYAB15 shown by Co-IP assay in maize protoplasts. ZmCPK2-MYC and ZmYAB15-GFP were transiently co-expressed in protoplasts of LH244. The immunoprecipated proteins with anti-GFP antibodies were used for immunobloting with anti-MYC or anti-GFP antibodies. * indicates the immunoprecipated ZmCPK2-MYC. **(C)** The interaction between ZmCPK2 or ZmCPK17 and ZmYAB15 shown by BiFC assay in *N. benthamiana* leaf cells. The YFP signals were visualized by a confocal microscopy. GUS-YNE and GUS-YCE were used as negative controls. Bar =20 µm. **(D)** ZmCPK17 interacts with ZmYAB15 in yeast two-hybrid assay. **(E)** The effects of Ca^2+^ and EGTA on the interactions between ZmCPK17 and ZmCPK2 or ZmYAB15 shown by the pull-down assay. Recombinant ZmCPK17-His, GST-ZmCPK2 and GST-ZmYAB15 were subjected to the assay and immunoprecipitated with GST agarose. Immunoblotting was conducted with anti-GST and anti-His antibodies. The empty GST was used as the negative control. **(F, G)** Co-location of ZmYAB15 and ZmCPK2 or ZmCPK17. ZmYAB15-GFP and ZmCPK2-mCherry or ZmYAB15-mCherry and ZmCPK17-GFP were co-expressed in maize protoplasts. The GFP and mCherry signals were visualized by a confocal microscopy and measured by Image J. White dotted lines marked the measured locations. Bar = 5 µm. **(H)** ZmYAB15 is phosphorylated by ZmCPK2 and ZmCPK17. Recombinant ZmCPK17-His, MBP-ZmCPK2 and MBP-ZmYAB15 were subjected to the phosphorylation assay. The autograph and CBB were shown. **(I)** ZmCPK2 and ZmCPK17 phosphorylate different domains of ZmYAB15. Recombinant GST-ZmCPK2 and ZmCPK17-His were subject to the phosphorylation assay with different truncated forms of GST-ZmYAB15 (GST-ZmY15) as substrates. zf: the Zinc finger domain; dis: the disordered domain; hmg: the HMG domain.

**Figure S11.**
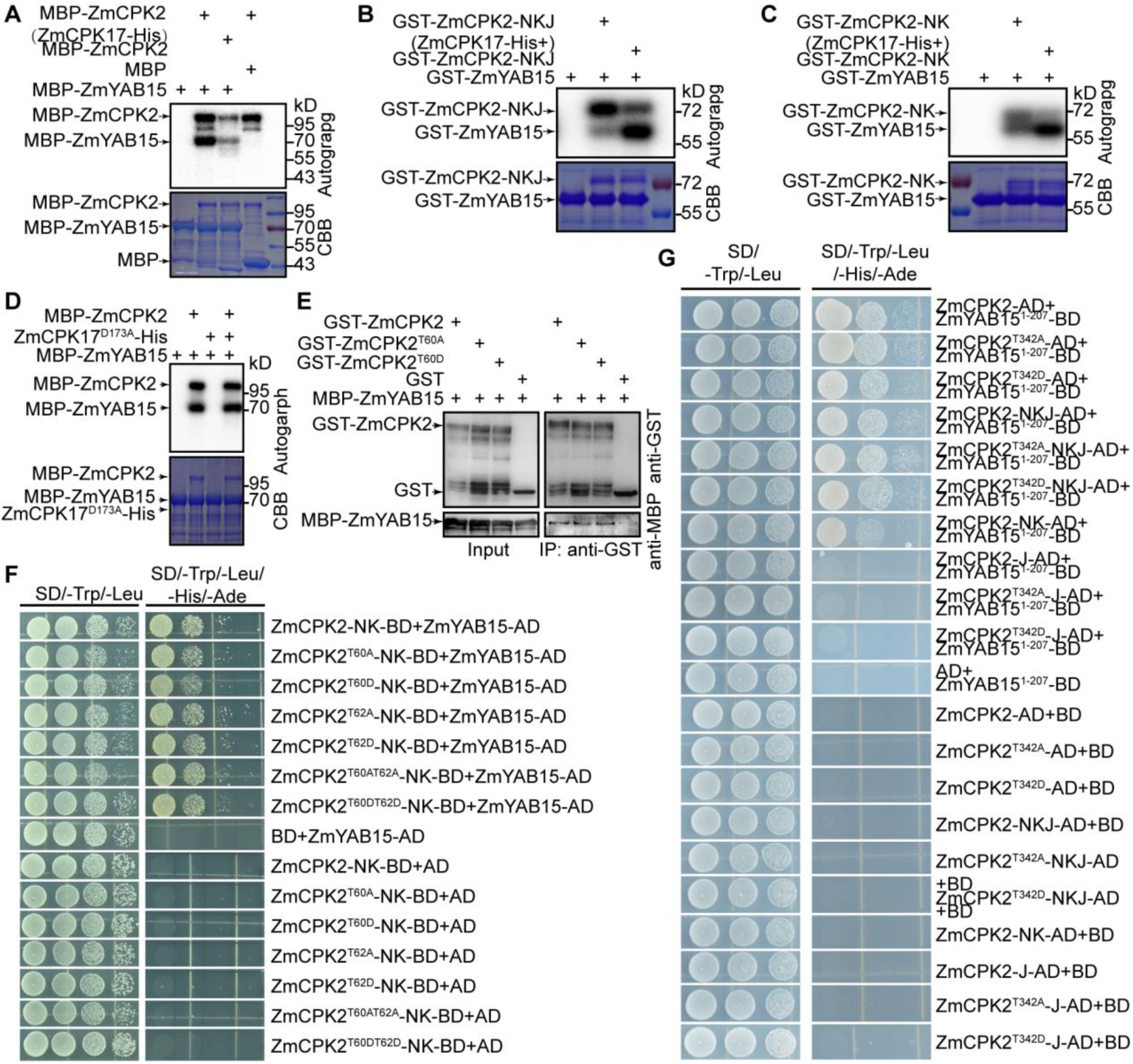
The phosphorylation of ZmCPK2 do not affect its interaction with ZmYAB15. **(A)** ZmCPK17 inhibits the kinase activity of ZmCPK2 on ZmYAB15. Recombinant ZmCPK17-His and MBP-ZmCPK2 were co-expressed or only ZmCPK2 was expressed in the *Escherichia coli* strain BL21 (DE3). MBP-ZmCPK2 purified from two transforms, respectively, was used for kinase assay. Recombinant MBP-ZmYAB15 was used as the substrate, and MBP as the negative substrate. **(B)** ZmCPK17 enhances the kinase activity of ZmCPK2-NKJ on ZmYAB15. Recombinant ZmCPK17-His and GST-ZmCPK2-NKJ were co-expressed or only ZmCPK2 was expressed in the *Escherichia coli* strain BL21 (DE3). GST-ZmCPK2-NKJ purified from two transforms, respectively, was used for kinase assay. Recombinant GST-ZmYAB15 was used as the substrate. **(C)** ZmCPK17 enhances the kinase activity of ZmCPK2-NK on ZmYAB15. Recombinant ZmCPK17-His and GST-ZmCPK2-NK were co-expressed or only ZmCPK2 was expressed in the *Escherichia coli* strain BL21 (DE3). GST-ZmCPK2-NK purified from two transforms, respectively, was used for kinase assay. Recombinant GST-ZmYAB15 was used as the substrate. **(D)** The effect of the kinase dead form ZmCPK17^D173A^ on the kinase activity of ZmCPK2 shown by *in vitro* phosphorylation assay. Recombinant ZmCPK17^D173A^-His, MBP-ZmCPK2 and MBP-ZmYAB15 were subjected to the assay. The autograph and CBB were shown. **(E)** ZmCPK2 T60A/D does not affect the interaction between ZmCPK2 and ZmYAB15 as shown by pull down assay. Recombinant GST-ZmCPK2, GST-ZmCPK2^T60A^ or GST-ZmCPK2^T60D^ and MBP-ZmYAB15 were subjected to the assay and immunoprecipitated with GST agarose, respectively. Immunoblotting was conducted with anti-GST and anti-MBP antibodies. **(F)** ZmCPK2 T60A/D and/or T62A/D does not affect the interaction between ZmCPK2 and ZmYAB15 in yeast two-hybrid assay. **(G)** ZmCPK2 T342A/D do not affect the interaction between different truncated forms of ZmCPK2 and ZmYAB15^1-207^ in yeast two-hybrid assay. ZmYAB15^1-207^ represents the first 1-207aa of ZmYAB15.

**Figure S12.**
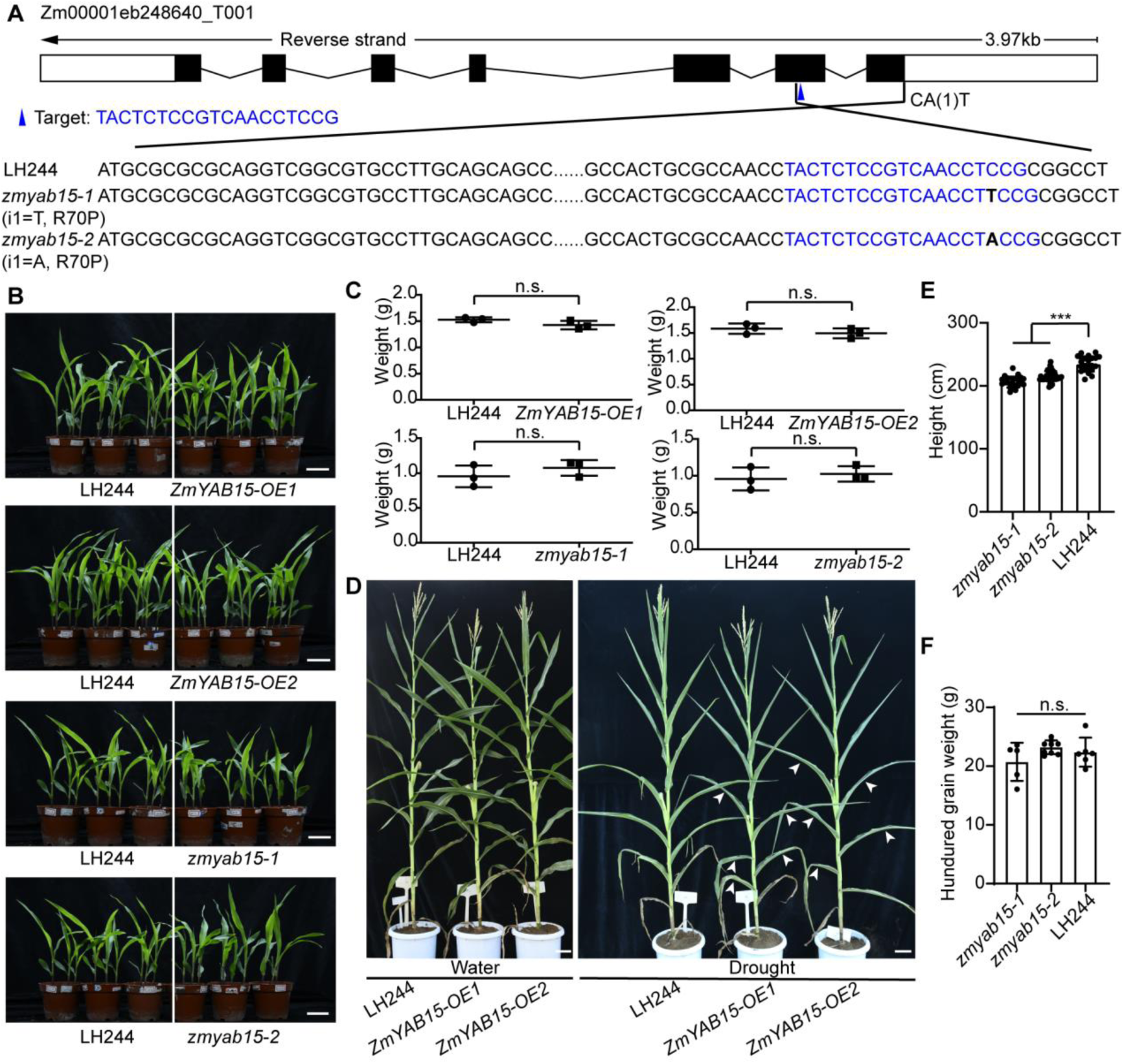
The growth and development phenotypes of *ZmYAB15-OE* and *zmyab15* plants. **(A)** Detail information for the *zmyab15-1/-2* mutations. The gene structure is shown in the top from Gramene database. Black boxes represent exon and black lines represent intron. The blue triangle indicates the location of the single guide RNA target in the second exon that creates two mutants *zmyab15-1/-2*, with a single base T/A insertion, both causing a frame shift after Leu69. **(B)** The growth and development phenotypes of *ZmYAB15-OE* and *zmyab15* plants. 12-d-old seedlings of *ZmYAB15-OE1/2*, *zmyab15-1/-2* or LH244 were separately grown in one pot under well-watered condition. **(C)** The weights of total plants in **(B)**. Each point represented a weight data from one pot. Error bars represent ± SD from one experiment. n.s. represents no significant difference with LH244, Student’s *t*-test. **(D)** Transgenic *ZmYAB15-OE1/2* plants are more sensitive to drought treatment compared with LH244 at the adult stage. The plants were grown in the field and the water group was well-watered (Water) while the drought group was stopped water supply at 20-d-old (Drought). Bar = 10 cm. **(E)** The height of *zmyab15* was shorter than LH244 in the field. Error bars represent ± SD. * Represents significant difference with LH244, *** p<0.001, Student’s *t*-test. **(F)** The hundred grain weight of *zmyab15* in the field. Error bars represent ± SD. n.s. Represents there was no significant difference with LH244, Student’s *t*-test.

**Figure S13.**
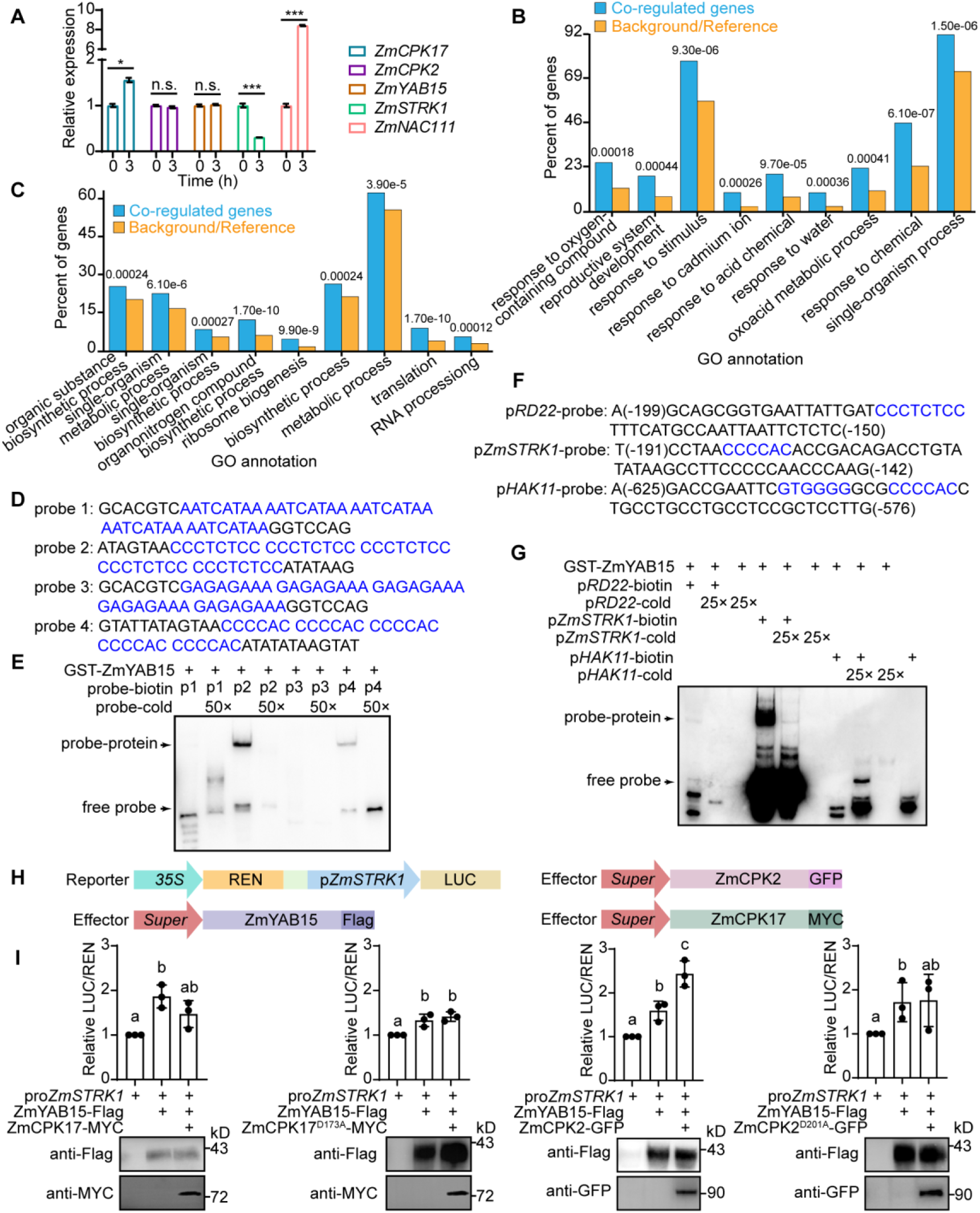
DNA binding activity of ZmYAB15 was regulated by ZmCPK2 and ZmCPK17. **(A)** Relative expression level analyses of *ZmCPK17*, *ZmCPK2, ZmYAB15* and *ZmSTRK1* in LH244 with ABA treatment for 0 h or 3 h. 8-d-old seedlings were well-watered and the total RNAs were extracted. Error bars represent ± SD of three technical replicates from one experiment. *ZmNAC111* was used as an ABA-induced marker gene. * Represents significant difference, * p<0.05, *** p<0.001, n.s. Represents there was no significant difference, Student’s *t*-test. **(B)** GO annotation analysis of ZmCPK2, ZmYAB15 and ZmCPK17 co-regulated genes. The abscissa represents a part of GO terms with significant enrichment, the ordinate represents the ratio of the genes enriched in the GO term to the total number of genes, the numbers represent the p value of the significance analysis compared to the reference genome. **(C)** GO annotation analysis of genes only changed in LH244. **(D)** The predicted probes used in EMSA. The blue letters represent four different candidate YABBY protein binding motifs that were reported previously. **(E)** EMSA shows that recombinant GST-ZmYAB15 binds C-rich probes *in vitro*. The upper and lower arrows indicate the bound probe and the free probe, respectively. Competitive experiment was performed using high concentration (50×) of cold probes. **(F)** The probe sequences used in **(G)**, which were 50 bp fragments including cis-element in the promoter of candidate genes. The numbers in parentheses showed the upstream location of probes from ATG. **(G)** EMSA shows that recombinant GST-ZmYAB15 binds to the promoter of *RD22* weakly and *ZmSTRK1* strongly *in vitro*, but not bind to the promoter of *HAK11*. Competitive experiment with high concentration (25×) of cold probes showed the binding specificity. **(H)** Sketch map of the reporter and the effector used in **(I)**. **(I)** Dual-LUC assay shows the effect of ZmCPK17 and ZmCPK2 on the transactivational activity of ZmYAB15. The constructs p*ZmSTRK1*:LUC, ZmYAB15-Flag, ZmCPK2-GFP, ZmCPK2^D201A^-GFP, ZmCPK17-MYC and ZmCPK17^D173A^-MYC were co-transformed in *N. benthamiana* leaves and 35S:*REN* was used as internal control. Relative LUC/REN ratio indicated the expression level of *ZmSTRK1.* Error bars represent ± SD of three independent experiments, and single letters indicates significant difference identified by unpaired *t*-test. The protein level of ZmYAB15-Flag, ZmCPK2^WT/D201A^-GFP and ZmCPK17^WT/D173A^-MYC were detected with anti-Flag, anti-GFP or anti-MYC antibodies as shown below.

## Supplemental Tables

**Table S1.**
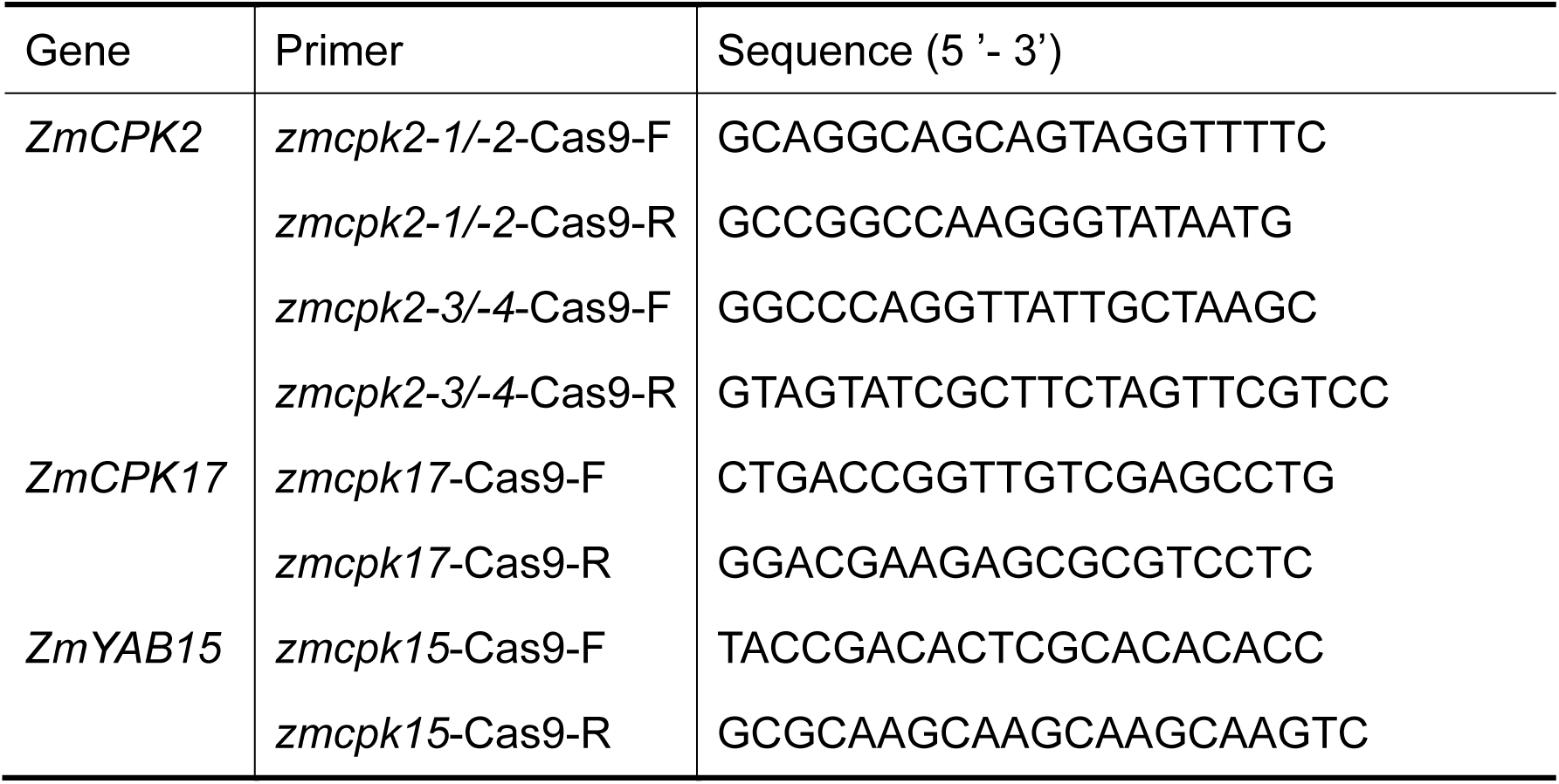
Primers used for genotyping.

**Table S2.**
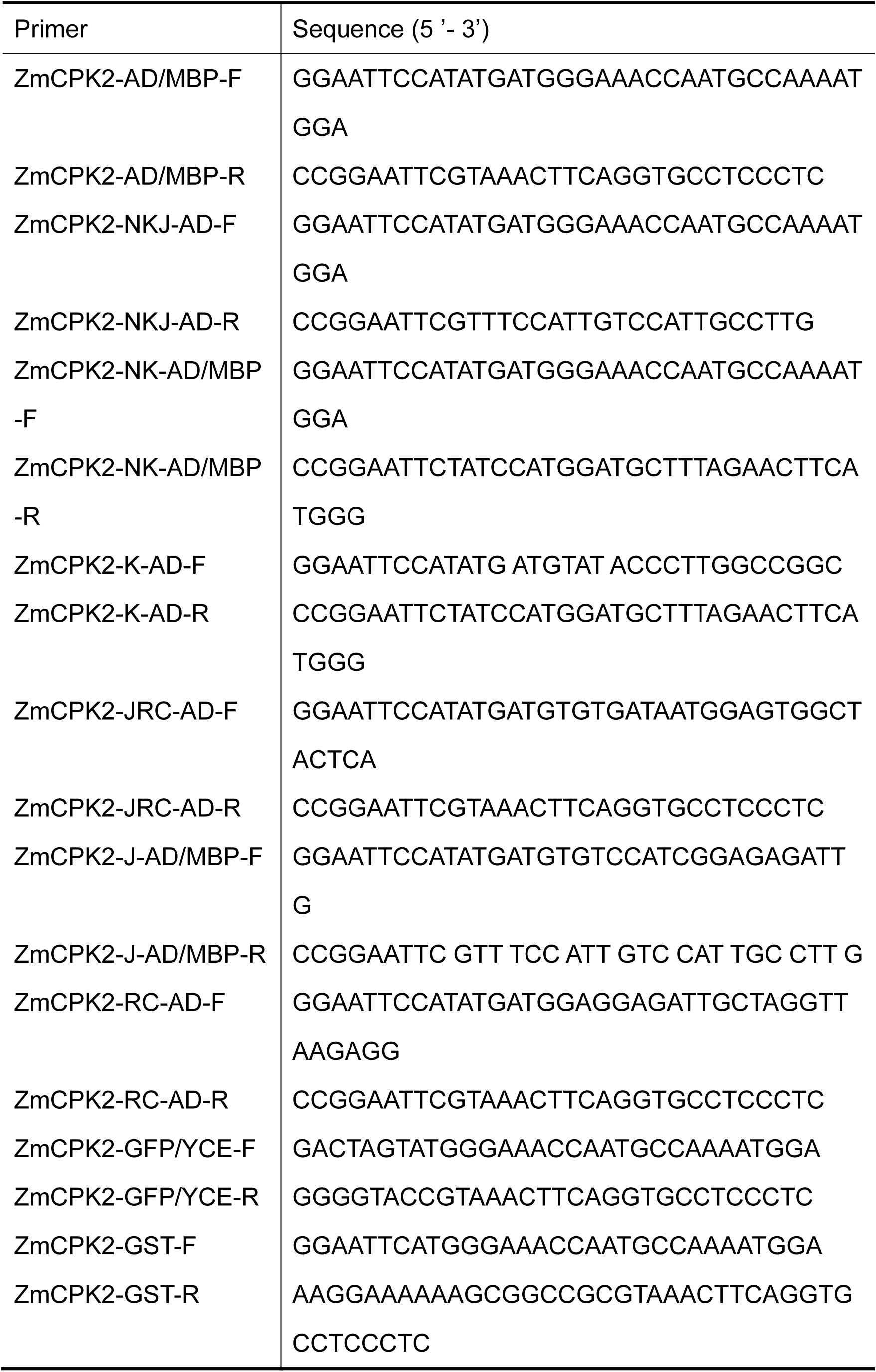
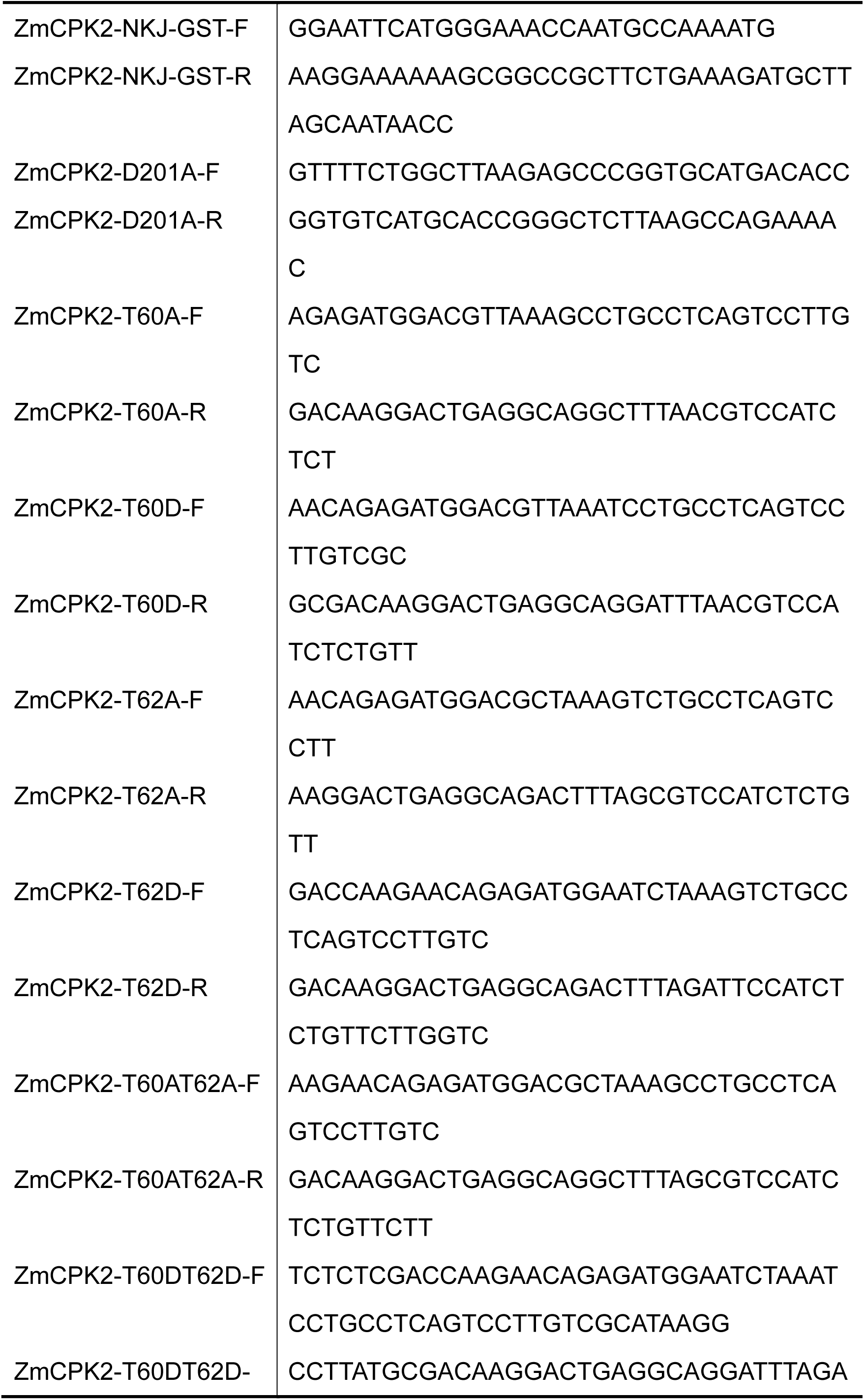
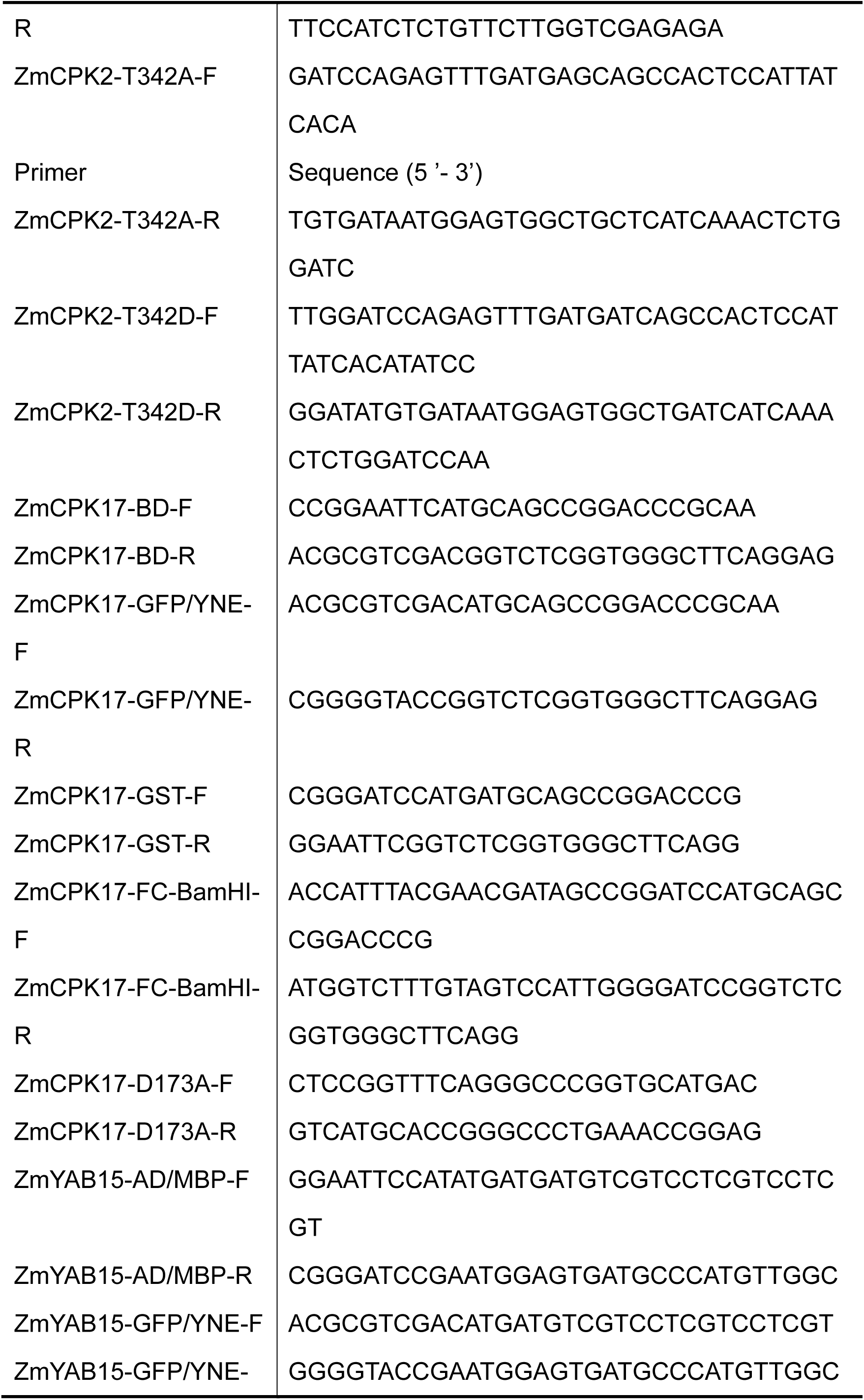
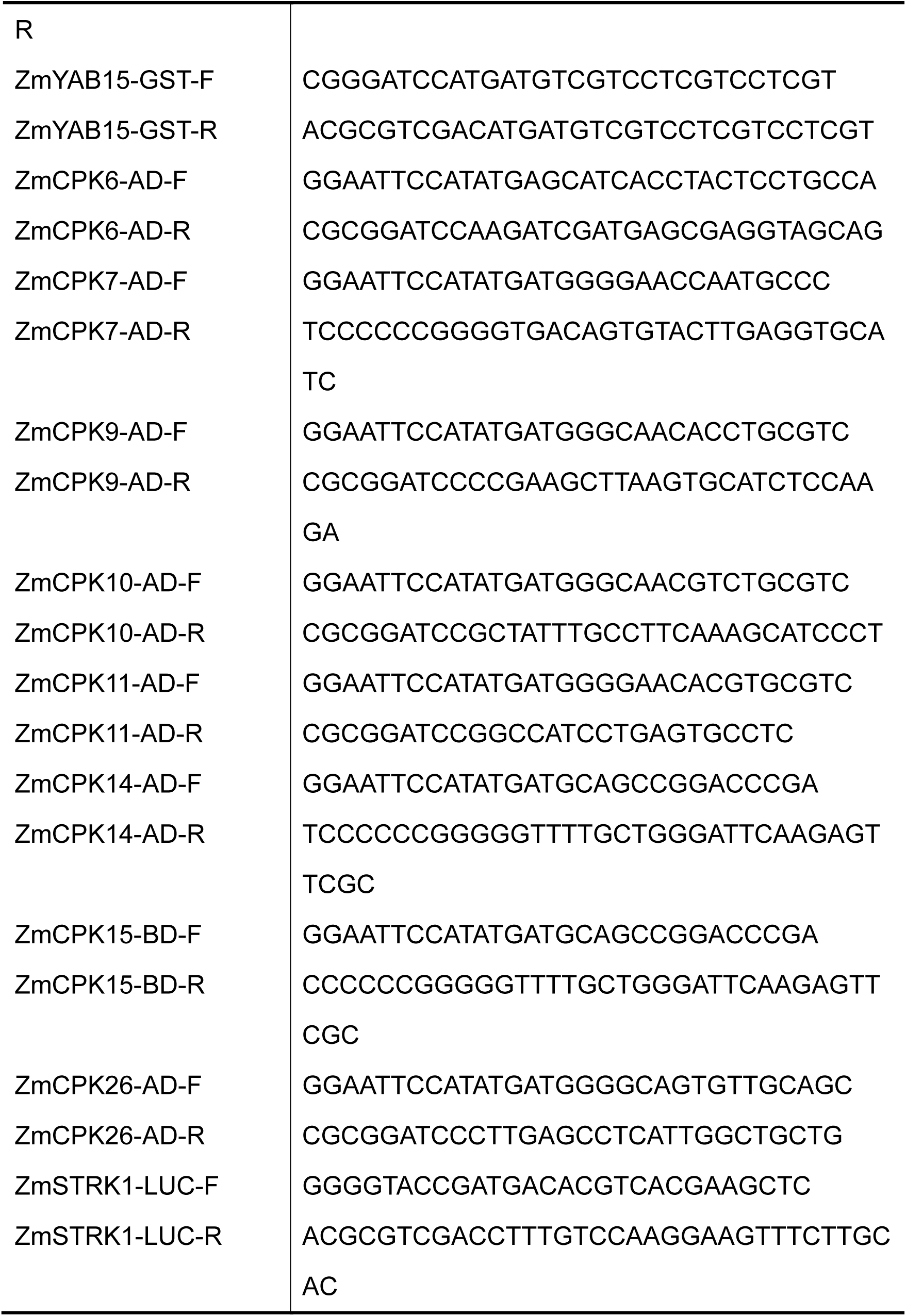
Primers used for plasmid construction.

**Table S3.**
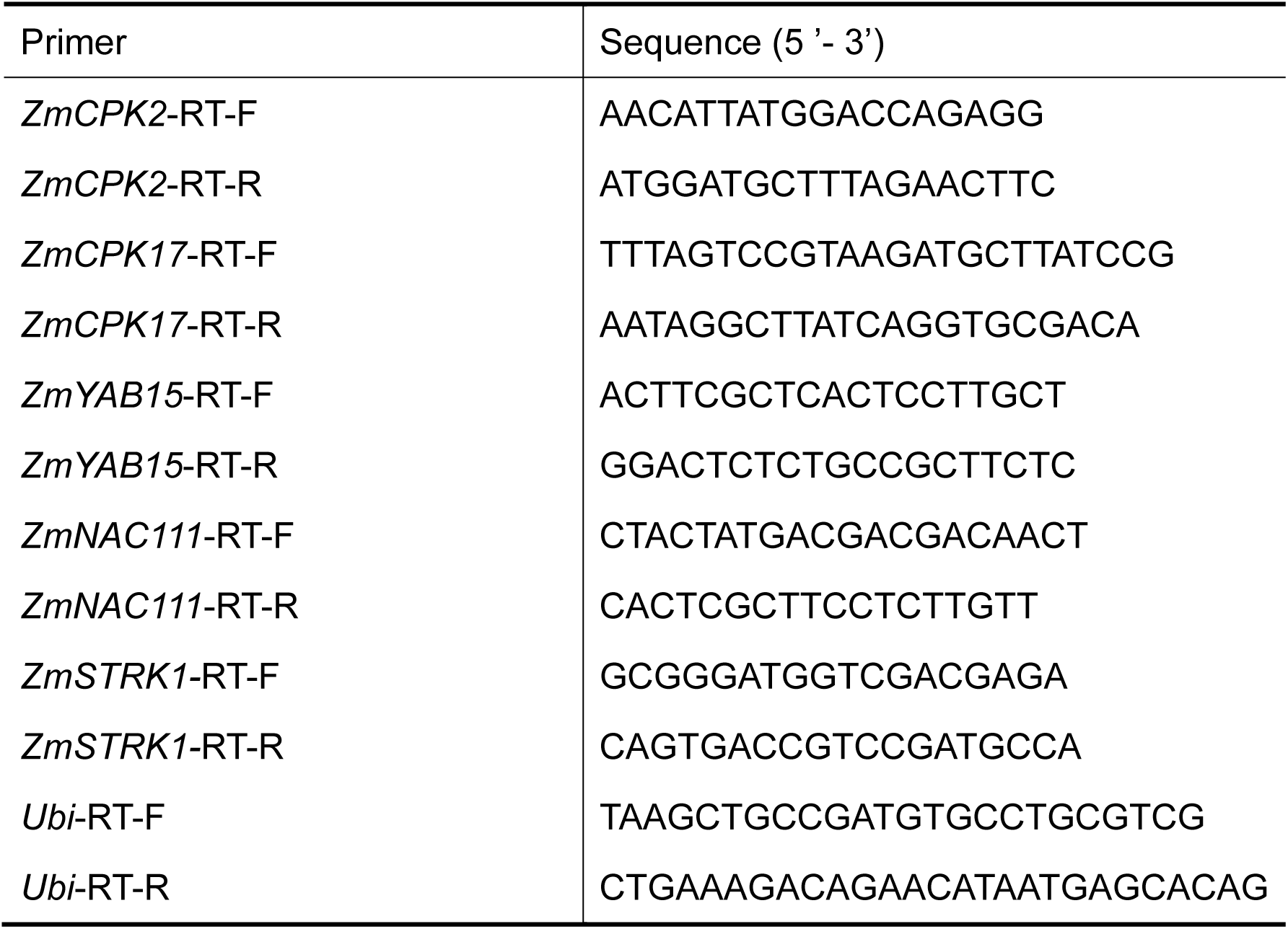
Primers used for qRT-PCR.

## Supplemental Experimental Procedures

### LC-MS/MS assay

To identify the phosphorylation site of ZmCPK2 by ZmCPK17 *in vitro*, 25 µg GST-ZmCPK2^D201A^ and 5 µg GST-ZmCPK17 purified proteins were incubated in 50 µl of protein protein kinase reaction buffer containing 25 mM Tris-HCl (pH 7.5), 10 mM MgCl_2_, 100 µM CaCl_2_ and 50 µM ATP at 30°C for 30 min. The reaction products were digested with trypsin (pH 8.5) at 37°C for 12 h and analyzed by LC-MS/MS.

### Yeast two-hybrid assay

Constructs were co-transformed into Gold yeast cells. Then the transfected yeast cells were placed on synthetically defined (SD)/-Leu/-Trp medium and cultured at 30°C for 3 d. The empty vectors pGADT7 and pGBKT7 were used as negative control, the vectors AtABI1-AD and AtOST1-BD were co-transformed as positive control. The interaction was determined on SD/-Ade/-His/-Leu/-Trp medium following the manufacturer’s protocol (Takara).

### BiFC assay

Construct of ZmCPK2-YCE, ZmYAB15-YNE or ZmCPK17-YNE/YCE was transformed into *Agrobacterium* GV3101, respectively, and two of them were co-injected into *Nicotiana benthamiana* leaves with the silencing suppressor P19 strain (Waadt and Kudla, 2008). The plants were grown in the greenhouse for 48 h, YFP signals were observed under confocal laser-scanning microscopy (ZEISS710, Carl Zeiss) with exciting light 488 nm.

### Co-IP assay

*Super1300*:*ZmCPK2-MYC* and *Super1300*:*ZmYAB15-GFP* were co-transformed into LH244 protoplasts, and *Super1300*:*ZmCPK17-GFP* vector was transformed into *ZmCPK2-HF* protoplasts. The protoplasts were incubated at 22°C for 14 h in darkness. Total proteins were extracted with IP buffer and incubated with anti-GFP agarose (gta-20, Chromo Tek) for 3 h. The immunoprecipitated samples were washed 3 times with PBS buffer and subjected to immunoblot analysis. The immunoprecipitated proteins were detected with anti-GFP (D089; TransGen Biotech), anti-MYC (G045; TransGen Biotech) or anti-Flag (F3165, Sigma-Aldrich) antibodies, respectively.

### Protein subcellular location

To detect the subcellular location and co-location of proteins, the constructs of *Super1300:ZmCPK2-GFP/mCherry*, *Super1300:ZmCPK17-mCherry/GFP* and *Super1300*:*ZmYAB15-GFP/mCherry* and *Super1300*:*AtMCM10-mCherry* were transformed into maize protoplasts individually or together, and incubated at 22°C for 14 h in darkness. GFP and mCherry fluorescence were observed by confocal microscopy (ZEISS710, Carl Zeiss) with exciting light 488 nm and 561 nm, respectively. GFP and mCherry signals were measured by Image J.

### Phylogenetic analysis

The amino acid sequences were obtained from Ensembl genome database (http://plants.ensembl.org/Zea_mays/Info/Index). Multiple sequences were aligned using Clustal W (Larkin et al., 2007), and the alignments were saved as mega format. A phylogenetic tree was built with the MEGA 5 software using the neighbor-joining method (Hall, 2013).The accession numbers of ZmCPKs as reported before were listed in Supplemental Data Set sheet 1 (Kong et al., 2013).

### Aequorin-based cytosolic Ca^2+^ measurements

For aequorin assay, maize protoplasts from LH244 were transiently expressed *UBQ10*:YFP-aequorin (Mehlmer, 2012) and used for ABA-induced [Ca^2+^] increase measurement. The procedure of quantitative cytosolic Ca^2+^ measurement was modified according to the previous study (Ma et al., 2019). Briefly, after 12 h incubation in darkness, the protoplasts were collected and divided into three 1.5 ml tubes equally. And then 100 µL solution including 4 mM MES, (pH 5.7), 0.6 M mannitol, 4 mM KCl, 20 µmol/L coelenterazine, with or without 50 µM BAPTA-AM was added to each tube, respectively, and the tubes were kept in the dark for 4 h at 22 °C. Signal values of luminescence were recorded by a luminometer (GLOMAX, Promega). The tube was set on the luminometer measurement chamber. Following 100 s of measurement (1 s interval) for resting luminescence, 100 µL incubation buffer (4 mM MES, (pH 5.7), 0.6 M mannitol, 4 mM KCl,) with or without 100 µM ABA was added into the tube and mixed gently. Then, the stimulated luminescence was recorded at 1 s interval for another 2000 s. The remaining aequorin was discharged by adding 100 µL discharge buffer (1 M CaCl_2_, 20% (v/v) ethanol) and the signal values were recorded for a further 500 s. The quantification of [Ca^2+^]_cyt_ (µmol/L) was calculated as described before (Rentel and Knight, 2004). At least 4 repeats for each treatment were tested. Δ [Ca^2+^]_cyt_ (µmol/L) represents intensity of the peaks in response to stimulus, which was calculated by comparing the three treatment (Mock, ABA, BAPTA-AM+ABA) induced [Ca^2+^]_cyt_^peak^ against to the rest [Ca^2+^]_cyt_^peak^. Each experiment chose 20 peaks from the rest and the treatment to indicate the maximum changes of [Ca^2+^]_cyt_. Three independent experiments were calculated.

### *In vitro* pull-down assay

Recombinant protein of GST-ZmCPK2, GST-ZmYAB15, GST or ZmCPK17-His was expressed in *E. coli* strain BL21 (DE3) and cultured at 18 °C for 14-16 h, respectively, and was purified using glutathione Sepharose 4B or Ni Sepharose. *In vitro* pull-down assays were performed as described previously (Ma et al., 2021). In brief, purified protein GST-ZmCPK2, GST-ZmYAB15 or GST (10 µg) was separately incubated with glutathione Sepharose 4B for 2 h in a tube with 100 µl PBS buffer (0.05% Nonidet P-40) at 4°C. After a brief centrifugation at 1000 g for 3 min at 4°C, the buffer was removed and 1 µg of ZmCPK17-His protein and 100 µl fresh PBS buffer (0.05% Nonidet P-40) with 100 µM CaCl_2_ or EGTA were added to the glutathione Sepharose 4B. The mixtures were rotated for 2 h at 4°C and washed five times with 1 ml PBS (1% Triton X-100) each time to remove nonspecifically bound proteins. The proteins were subjected to immunoblot analysis with anti-GST or anti-His antibodies (TransGen Biotech).

### EMSA assay

Electrophoretic mobility shift assay (EMSA) was performed as previous described (Zhu et al., 2020). The recombinant protein GST-ZmYAB15 was purified with Glutathione Sepharose 4B. The 5’-biotin-labeled DNA primers were made by 10 min of 100°C denaturation and renaturation in room temperature. Each 20-µL volume of binding reaction contained 2 nM - 200 nM of biotin-probe and 2.5 µg protein and was incubated at 27°C for 20 min. The reaction products were stopped by adding 5 × native sample loading buffer and separated by 6.5% (v/v) native polyacrylamide gel electrophoresis. The binding signal on gel was transferred onto a nitrocellulose membrane and detected using the LightShift Chemiluminescent EMSA Kit (Thermo Fisher Scientific) according to the manufacturer’s instructions.

### Dual-luciferase (dual-LUC) activity assay

The dual-LUC transcriptional activity assay was performed as described previously (Li et al., 2022). The *ZmSTRK1* promoter fragment (-2 kb from ATG site) was cloned to the pGreenII 0800-LUC vector system (p*ZmSTRK1*:LUC), which includes a firefly luciferase (*LUC*) gene driven by the promoter of a target gene and a *Renilla* luciferase (*REN*) gene driven by the constitutive 35S promoter as an internal control. Construct of p*ZmSTRK1*:LUC, *Super1300*:ZmYAB15-Flag, *Super1300*:*ZmCPK2-GFP, Super1300*:*ZmCPK2^D201A^-GFP*, *Super1300*:*ZmCPK17-MYC* and *Super1300*:*ZmCPK17^D201A^-MYC* was transformed into *Agrobacterium* GV3101, respectively, and different combination of them were co-injected into *Nicotiana benthamiana* leaves with the silencing suppressor P19 strain. After 48 h in the normal growth condition, 50 µM ABA was injected into the leaves expressed Super1300:ZmCPK17-MYC for 1 h. Total protein were extracted from the leaves using dual-LUC assay reagents (Promega). The relative LUC/REN ratio was used to measure *ZmSTRK1* promoter activity, as detected using a GloMax 20/20 luminometer. The protein levels were detected by western blot with anti-Flag, anti-GFP or anti-MYC antibodies.

